# Evidence for a reproductive sharing continuum in cooperatively breeding mammals and birds: consequences for comparative research

**DOI:** 10.1101/2023.05.05.539509

**Authors:** Yitzchak Ben Mocha, Tal Dahan, Yuqi Zou, Michael Griesser, Shai Markman

## Abstract

Extreme reproductive skew occurs when a dominant female/male monopolises reproduction within a group of multiple sexually mature females/males. It is sometimes considered an additional, restrictive criterion in the definition of cooperative breeding (i.e., when conspecifics provide parental care to other group members’ offspring). However, datasets that use this restrictive definition to classify species as cooperative breeders have two critical shortcomings. First, reproductive skew is systematically overestimated by including groups with a single sexually mature female/male when calculating the reproductive output of “dominant” females/males. Second, a lack of reporting on which species are classified based on limited data prevents accounting for uncertainty in classification. Considering these shortcomings, we show that extreme reproductive skew in multi-female and multi-male groups only occurs rarely in species previously classified as cooperative breeders using restrictive definitions (11 mammal species, 12 bird species). We further provide updated datasets on reproductive sharing in multi-female/male groups of cooperatively breeding mammals and birds that allow accounting for classification uncertainty. Our results demonstrate a reproductive sharing continuum even among those cooperatively breeding species argued to exhibit extreme reproductive skew. At the practical level, these findings call for significant changes in datasets that classify species by social systems. At the conceptual level, we suggest that reproductive skew should not be a defining criterion of cooperative breeding.

## 1. Introduction

Cooperative breeding is a reproductive system where conspecifics provide parental care to the offspring of other group members [i.e., alloparental care 1,2]. For instance, in a range of species, alloparents provide systematic allonursing [3,4], allofeeding [5,6], babysitting [7] and transference of offspring between locations [8,9]. In some cases, alloparents care for others’ offspring while not reproducing themselves, leading to reproductive skew within social groups [e.g., common mole-rat:, 10, Florida scrub jay:, 11].

Within-group reproductive skew is of fundamental importance in cooperative-breeding research because it is sometimes considered a defining criterion of cooperative breeding [e.g., 12–14]. According to this restrictive definition, species that engage in alloparental care should be divided into communal and cooperative breeders [15,16]. Communal breeding species engage in alloparental care while group members are said to share reproduction to some extent. In contrast, cooperative breeding species engage in alloparental care, but within-group reproduction is argued to be extremely skewed in favour of a single female and/or male [15,16]. Comparative studies that apply this restrictive definition to classify species binary, as cooperative breeders or not, are therefore, critically dependent on the correct identification of species exhibiting both alleged traits of cooperative breeding: alloparental care and extreme female and/or male reproductive skew [17].

Datasets that classify mammalian and avian societies according to the restrictive definition of cooperative breeding, though, have two crucial shortcomings: (i) they systematically overestimate the prevalence of extreme reproductive skew across species, and (ii) they do not indicate species whose classification of social system is based on limited sample size or data that is potentially biased for different reasons. Thereby, preventing accounting for uncertainty in the classification. Below, we discuss these shortcomings and their consequences for comparative research of cooperative breeding.

### 1.1. Shortcoming I: Overestimation of reproductive skew

Ideally, within-group reproductive skew should be calculated with one of the indices controlling for factors affecting reproductive skew besides social suppression [e.g., group size, age, kinship:, 18,19]. These indices, though, require detailed longitudinal data [18,20] that are rarely available for wild animal populations. To overcome this deficiency of data, comparative studies that classify numerous species use relatively rough proxies of reproductive skew. For instance, the percentage of offspring born to the most dominant females in social groups, or the percentage of groups with a single breeding female within a population [e.g., 21,22]. Critically, these percentages are often calculated from all sampled groups, including those with a single sexually mature female/male (hereafter “single-female groups” or “single-male groups”). This approach, however, runs contrary to the fundamental rationale of determining reproductive skew. That is, measuring the divergence from equal reproductive sharing within a group of *potentially* reproductive individuals of a specific sex [16,18,20,23–25]. Measuring reproductive “skew” or “suppression” is, therefore, only meaningful where the reproductive potential of at least one group member can be compromised (i.e., group size ≥2 sexually mature individuals of the sex in question; hereafter “multi-female groups” or “multi-male groups”) [16,26–28].

### 1.2. Shortcoming II: Controlling for the certainty of species classification

A second important shortcoming is that, despite the paucity of parentage data in many species, current datasets often do not indicate species whose classification is based on limited sample size or potentially biased data [e.g., data from experimental populations; see also 17]. Comparative studies that use these datasets are therefore forced to treat the classification of all species with the same degree of confidence, despite the availability of statistical methods to account for the uncertainty of some species’ classification.

### 1.3. Revisiting the underlying biological assumption of the restrictive definition of cooperative breeding

Given these two shortcomings, we revisited the underlying biological assumption of the restrictive definition of cooperative breeding (i.e., restricting cooperative breeding to species with alloparental care and extreme reproductive skew in females and/or males). Namely, we tested whether there is a quantitative distinction between the reproductive skew of communal and cooperative breeding species [22]. To the best of our knowledge, the only empirical support for this assumption comes from a comparative study on mammals claiming that dominant females in cooperative breeding species produce 88-100% of offspring (N = 26 species), while in other social mammals dominant females produce only 8-69% of offspring [N = 20 species, Figure 1, 22]. However, if this underlying biological assumption of the restrictive definition is incorrect, we predicted that species with alloparental care would rather show a continuum of reproductive sharing [29,30].

**Figure 1:**
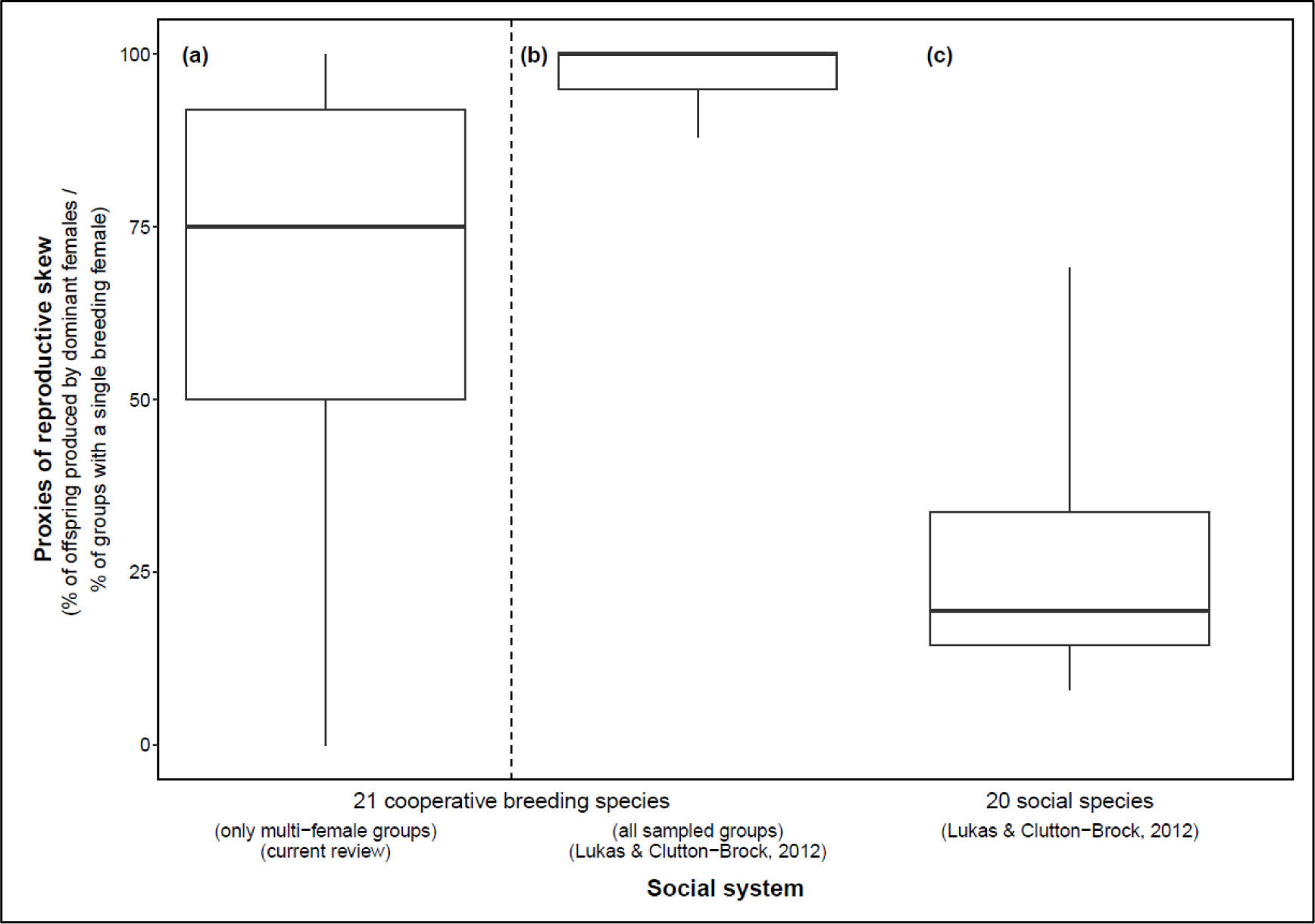
Proxies of female reproductive sharing in social mammals (amended from [22]). The reproductive output of dominant females in (b) cooperative breeding species (N = 21 species) and (c) “social” species (N = 20 “communal breeding” [i.e. where most adult females breed regularly and share care such as allonursing or feeding offspring] and “social breeding” species [i.e., where females live in groups and are neither cooperative nor communal breeders]) as reported and defined by Lukas & Clutton-Brock [22]. (a) The reproductive output of dominant females in the same 21 cooperative breeding species as in (b) while only considering multi-female groups. We excluded five species that were classified as cooperative breeders by Lukas & Clutton-Brock [22] because we could not find data on reproductive sharing (*Canis aureus*, *Castor fiber*, *Cryptomys ochraceocinereus*) or alloparental care (*Peromyscus californicus*, *Peromyscus polionotus*) in these species (see Table S1). Excluding these species made the difference between (a) and (b) less extreme as the range of reproductive skew reported for these five species by Lukas & Clutton-Brock [22] was 100%. Horizontal bars represent the median, boxes the 25% and 75% quartile and vertical lines indicate the minimum and maximum values.

To this end, we re-estimated the degree of reproductive sharing in 41 mammal and 37 bird species that were classified as exhibiting alloparental care and extreme female and/or male reproductive skew in four frequently used datasets [12,16,21,22]. In doing so, we only consider (where possible) social groups with multiple sexually mature females and/or males. To facilitate future research, we provide estimations for the certainty of the social classification of each species (e.g., based on sample size).

## 2. Methods

### 2.1. Revisited datasets

In mammals, we revisited the classification of 41 species that were classified as exhibiting alloparental care and an extreme female reproductive skew in at least one of three datasets: Raihani & Clutton-Brock [21] (7 species), Lukas & Clutton-Brock [22] (34 species) and Federico *et al*. [16] (13 species). Following the curators of these datasets, we only examined the female reproductive skew for mammals.

In birds, we revisited the classification of 37 species that were classified as exhibiting alloparental care and extreme female reproductive skew [23 species: Raihani & Clutton-Brock, 21] or as species exhibiting alloparental care and helpers that do “not breed or had zero-to-limited opportunities to breed” [36 species: Cornwallis et al., 12]. As Cornwallis and colleagues did not indicate the sex of helpers, for birds, we examined reproductive skew among females and males. Yet, the common eider (*Somateria mollissima*) was excluded from the male dataset as it was only classified as a cooperative breeder by Raihani & Clutton-Brock [21] who examined female reproductive skew.

### 2.2. Literature review

For each of the above-mentioned mammal and bird species, we implemented three search steps for data on reproductive sharing. First, we reviewed the citations that were used to support the species’ classification in the datasets that classified this species. Second, if this reference indeed included relevant data on parentage, we used Google Scholar to identify studies that cited this supporting reference. The title and/or abstract of each paper found in these searches were reviewed and relevant papers were further examined. Third, to ensure a comprehensive review of the literature on each species, we searched Web of Science using the following search command) TOPIC = English species name* AND (extra pair* OR extrapair* OR extra-pair* OR cooperative breeding* OR alloparental care* OR helpers* OR social system* OR parentage* OR maternity* OR paternity* OR genetic* OR DNA* OR reproduction*)(OR (TOPIC = Latin species name * AND (extra pair* OR extrapair* OR extra-pair* OR cooperative breeding* OR alloparental care* OR helpers* OR social system* OR parentage* OR maternity* OR paternity* OR genetic* OR DNA* OR reproduction*)). The title and/or abstract of each paper found in these searches were reviewed and relevant papers were examined. This procedure was repeated until no more new papers were found. Unfourtantly, our review was limited to publications in the English language.

In addition, we revisited the occurrence of alloparental care in each of examined species. Since the focus of this work is the prevalence of extreme reproductive skew, we used a relaxed definition of alloparental and considered any type of direct or indirect alloparental care [2]. Note that some scholars may reject indirect helping behaviours as alloparental care [e.g., indirect thermoregulation by non-parents in Alpine marmots:, 31] and, consequently, also the classification of these species as cooperative breeders [see discussion in 32]

Only primary literature describing wild populations was considered relevant for our review. We did not consider secondary literature including reviews and data from captive groups or introduced populations. The species were reviewed during the years 2021-2023, and our results, therefore, reflect the literature published before 2021.

### 2.3. Proxies of reproductive skew

A proxy of reproductive skew was calculated from each study that included relevant data. This resulted in species with multiple data points representing different populations, different study periods of the same population or different samples (e.g., samples of groups in which helpers were/were not related to the opposite sex breeder, see for example the splendid fairy-wren in Table S2). We calculated two proxies of reproductive skew. First, we summed the number of offspring produced by the most dominant female/male across all multi-female/multi-male groups and calculated the percentage of this number out of the total number of offspring produced in multi-female/multi-male groups, respectively (hereafter the “parentage-skew” proxy).

Second, we calculated the percentage of groups with a single breeding female/male (within a multi-female/multi-male group) out of the total number of multi-female/multi-male groups (hereafter the “skew-of-groups” proxy). One advantage of the skew-of-groups proxy is providing an estimation of reproductive sharing even when the dominance rank of breeders is unknown. As a rule, we calculated the skew-of-groups proxy out of the total number of years during which all groups sampled (i.e., from multiple samples of the same social groups over several seasons/years; hereafter “group-years”). Nonetheless, when data was detailed enough to infer that group composition remained similar across years, we used the number of unique groups (and not group-years) to calculate the skew-of-groups proxy (e.g., apostlebird in Table S2). This latter approach enables detecting reproductive sharing that occurs over multiple years [33,34]. For example, when different female group members reproduce across different years, as is often the case in mammal species that breed year-round [e.g., common marmosets in Brazil:, 35].

In addition, where data was detailed enough, the following factors were excluded from our calculations: (i) extra-group parentage [since restrictive definitions of cooperative breeding concern within-group reproductive skew:, 36], (ii) groups without at least one adult male and female (since these groups are not functionally reproductive units), (iii) groups without at least one helper that was unrelated to the opposite sex breeder [i.e., groups with incest limitation were excluded; see discussion in 37], and (iv) clutches/litters with only one sampled offspring (since shared maternity/paternity cannot be detected in these clutches/litters). See Tables S1-S2 for specific details of the samples used in each species.

### 2.4. Binary classification of species exhibiting extreme reproductive skew or not

We classified whether each species exhibits a female (for mammals and birds) and/or male (for birds only) extreme reproductive skew (yes/no). As a conservative approach, we used the more relaxed threshold applied by curators of the examined datasets [21,22]. According to this threshold, species exhibit an extreme female/male reproductive skew if the most dominant female/male in each group produced >90% of the offspring in their groups (i.e., the parentage-skew proxy) or if >90% of the groups had a single breeding female/male (i.e., the skew-of-groups proxy).

#### 2.4.1. Species with ambiguous proxies

For five mammal (Arctic fox: *Vulpes lagopus*, Ethiopian wolf: *Canis simensis*, saddle−back tamarin: *Saguinus fuscicollis*, cotton−top tamarin: *Saguinus oedipus*, dhole: *Cuon alpinus*) and seven bird species (Grey-crowned babbler: *Pomatostomus temporalis*, Tasmanian native hen: *Tribonyx mortierii*, Sociable weaver: *Philetairus socius*, Laughing kookaburra: *Dacelo novaeguineae,* Arabian babbler: *Turdoides squamiceps,* Brown jay: *Cyanocorax morio,* Apostlebird: *Struthidea cinerea*) different studies provided proxies that were below and others above the threshold of extreme reproductive skew (Figures 2-4). In the binary classification of these species as exhibiting extreme reproductive skew or not, we prioritised data according to the following rules (see details per species in Tables S1-S2):

(i) Data from a sample that included only multi-female groups/multi-male groups override data from a mixed sample of single and multi-female/multi-male groups (hereafter “mixed sample”). As a rule, data from mixed samples were only considered if a sample including solely multi-female or multi-male groups was not found or was very small.

(ii) When dominance ranks were known for both proxies, data for the parentage-skew proxy override data for the skew-of-groups proxy.

(iii) Genetic evidence override behavioural or physiological evidence of parentage (e.g., lactation, the number of placental scars or abnormally large clutch size).

(iv) Studies with larger sample sizes override studies with smaller sample sizes.

(v) Reproductive suppression may occur at any reproductive stage from interference with copulation [38,39] until infanticide [39,40]. We hence prioritised data on the reproductive output in later reproductive stages that are less likely to be aborted in the following order: number of weaned/fledged offspring, number of offspring, and number of pregnant females. For a valid comparison between dominant and subordinate group members, per a study, we only considered the type of reproductive output that was available for both dominance categories.

**Figure 2.**
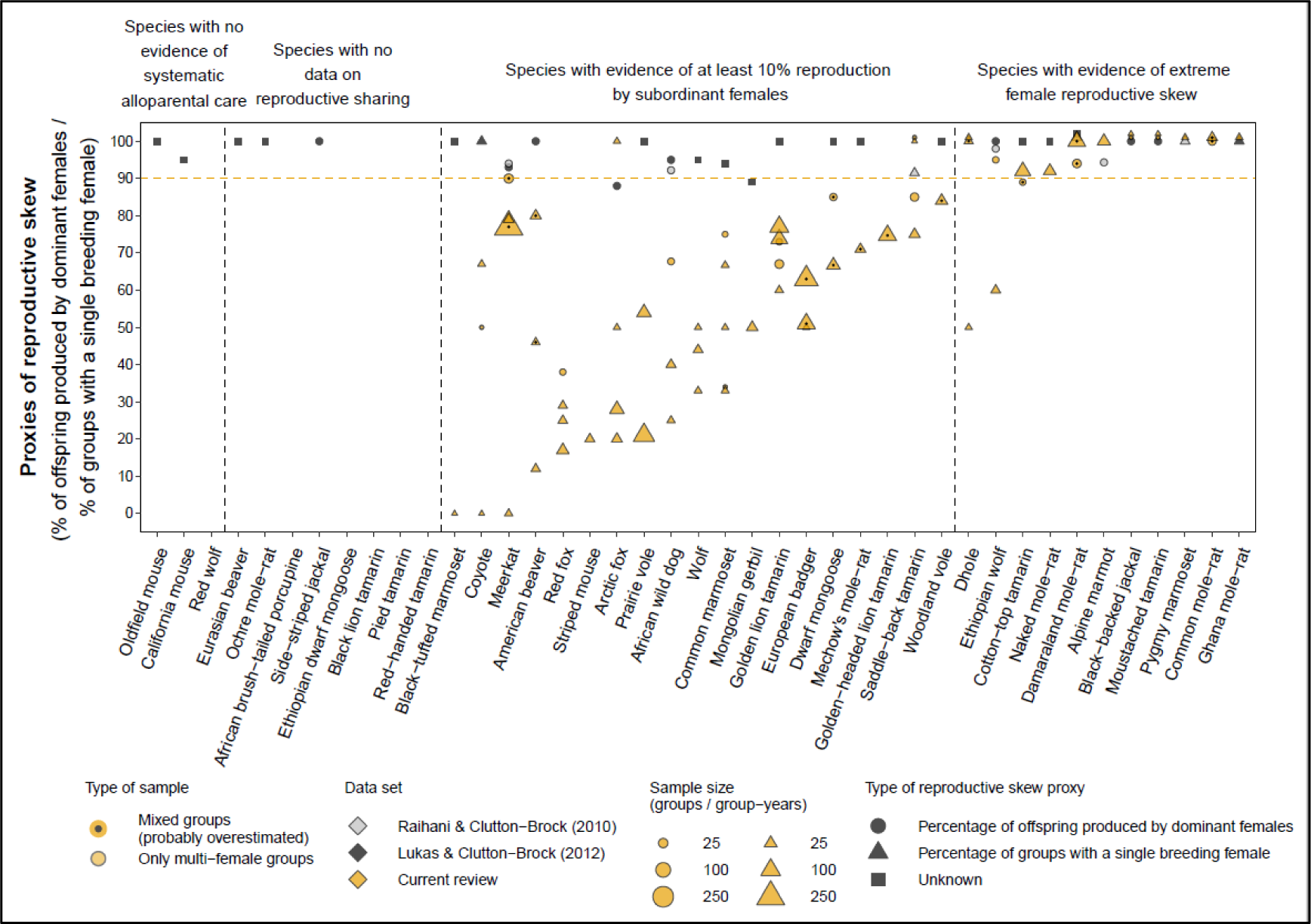
Within-group female reproductive sharing in the 41 mammal species classified as exhibiting alloparental care and an extreme female reproductive skew in Raihani & Clutton-Brock [21] and/or Lukas & Clutton-Brock [22] and/or Federico *et al*. [16] datasets. The orange line represents Raihani & Clutton-Brock [21] minimum threshold for the extreme female reproductive skew of >90% reproductive output by dominant females. Mixed samples may include groups with a single female and, thus, likely overestimate reproductive skew. Multiple data points for the same species represent different sample types, study periods or populations. For species where we estimated minimum and maximum possibilities of reproductive sharing, the mean value is presented (see Table S1 for details). Data from Raihani & Clutton-Brock [21] and Lukas & Clutton-Brock [22] are not adjusted for sample size as the authors did not report sample size. Species without data points are species for which we could not find data and/or the examined datasets did not include quantitative values for their classification. For clarity, overlapping data points for the same species were separated by placing one of them one percent higher.

**Figure 3.**
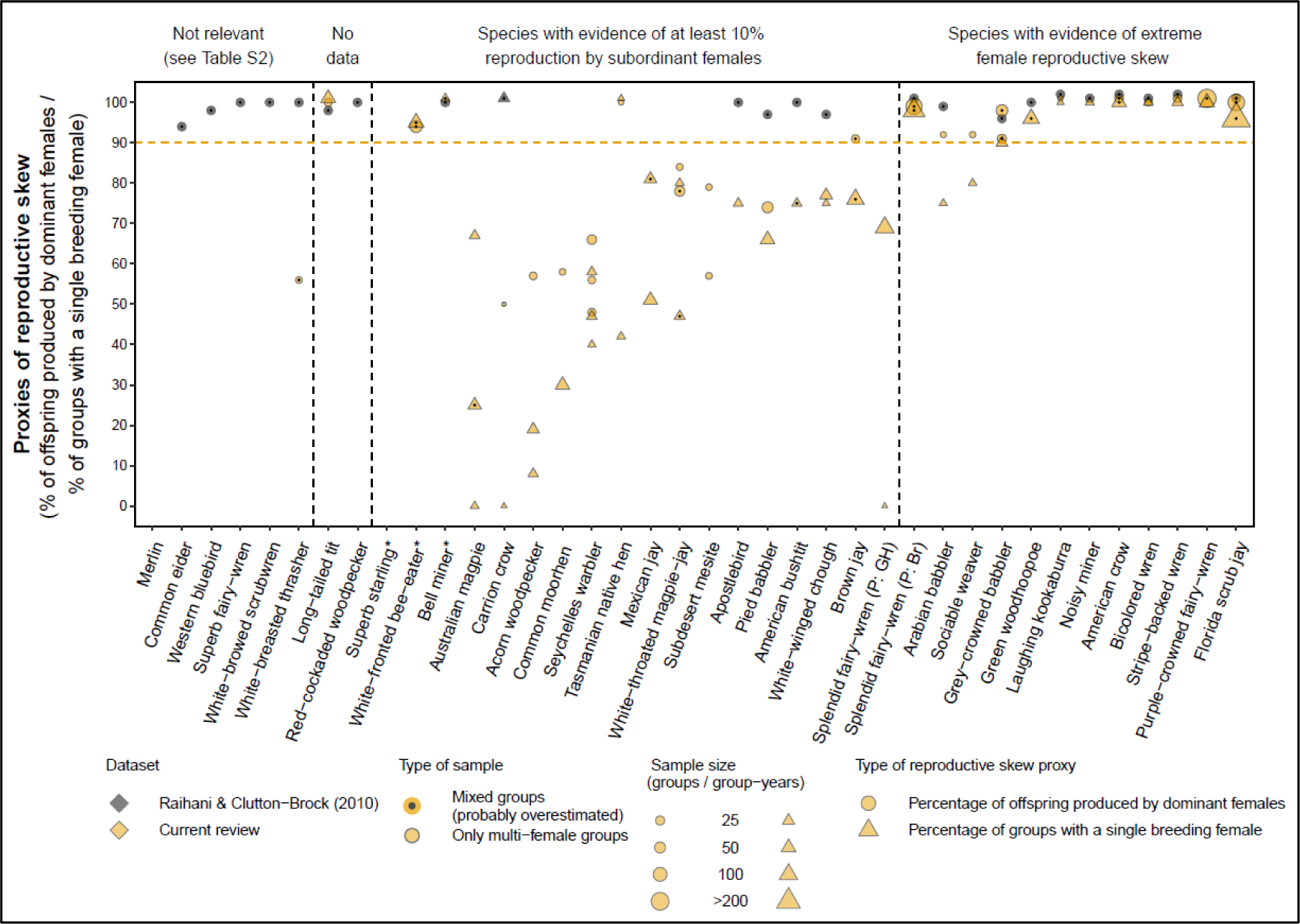
Within-group female reproductive sharing in the 37 bird species classified as exhibiting alloparental care and an extreme female reproductive skew in Cornwallis *et al.*, [12] and/or Raihani & Clutton-Brock [21]. The orange line represents Raihani & Clutton-Brock’s [21] minimum threshold for extreme female reproductive skew. Mixed samples may include groups with a single female and, thus, likely overestimate reproductive skew. Multiple data points for the same species represent different samples, study periods or populations. Data from Raihani & Clutton-Brock [21] are not adjusted for sample size. Species without data points are species for which we could not find data and/or the examined datasets did not provide quantitative values for their classification. For clarity, overlapping data points for the same species were separated by placing one of them one percent higher. Species with an asterisk are species living in multi-level societies in which the basic social unit consists of multiple breeding pairs (see Table S2 for details). Splendid fairy−wren is represented twice as two populations of the species exhibit different degrees of reproductive skew (see Table S2 for details).

**Figure 4.**
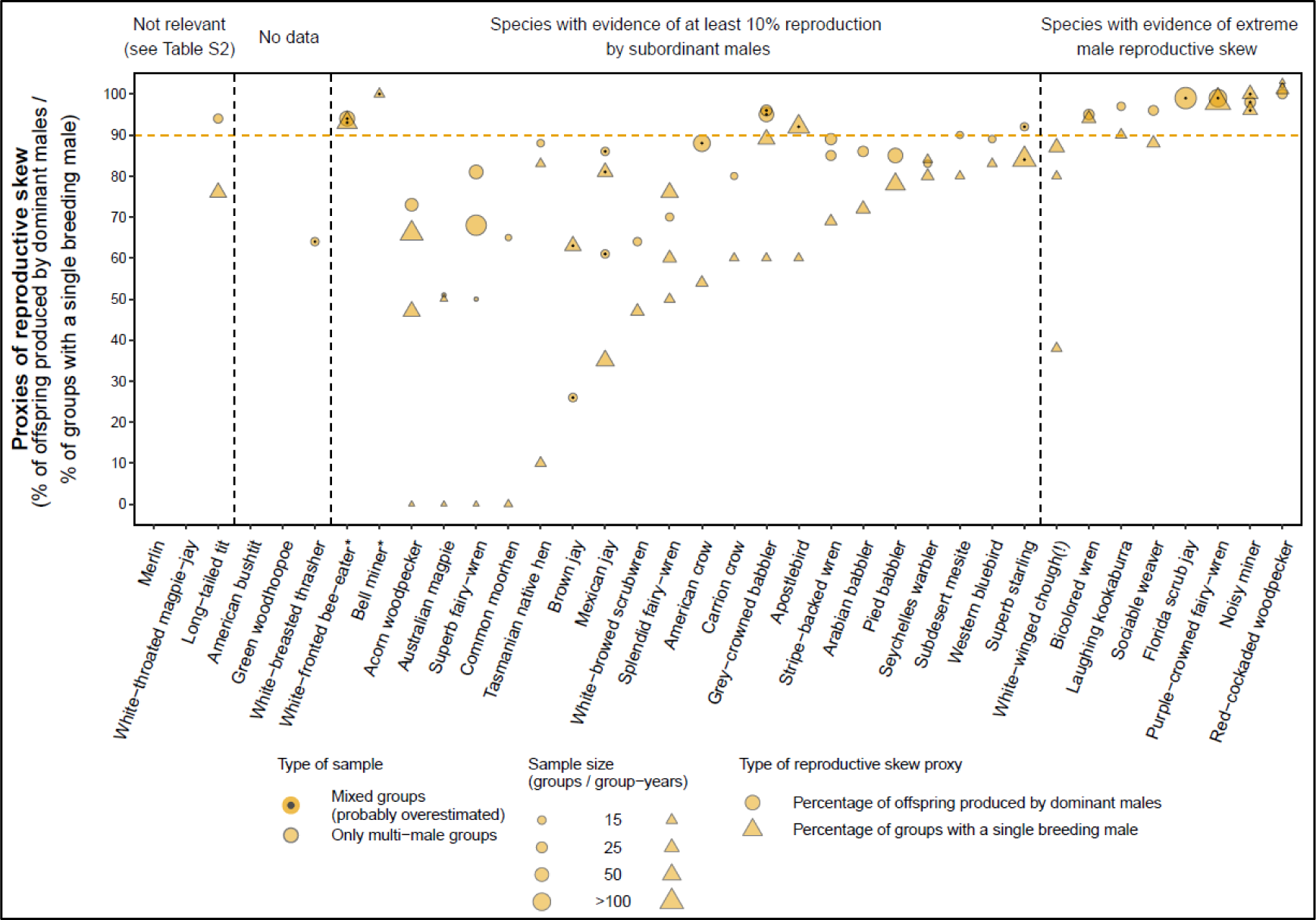
Within-group male reproductive sharing in the 36 bird species classified as exhibiting alloparental care and an extreme reproductive skew in Cornwallis *et al.* [12]. The orange line represents Raihani & Clutton-Brock’s [21] minimum threshold for extreme reproductive skew. Mixed samples may include groups with a single male and, thus, likely overestimate reproductive skew. Multiple data points for the same species represent different samples, study periods and/or populations. Species without data points are species for which we could not find data. For clarity, overlapping data points for the same species were separated by placing one of them one percent higher. Species with an asterisk are species living in multi-level societies in which the basic social unit consists of multiple monogamous breeding pairs (see Table S2 for details). White−winged chough is marked with an exclamation mark (!) because although reproduction is shared in newly formed groups, it is usually monopolised in socially established groups (see Table S2 for details).

### 2.5. Evaluation of the certainty of species classification

We evaluated the strength of evidence for each species’ classification qualitatively and quantitatively. Qualitatively, we indicated the type of sample used in each study (i.e., samples consisting only of multi-female groups versus mixed samples of single-female and multi-female groups). For the reasons described in the introduction and discussion, we assumed that skew proxies that are calculated from mixed samples likely overestimate the reproductive skew.

Quantitatively, we indicated the “combined” sample size for each species’ classification. Namely, we combined the sample sizes from all studies used to classify a species and categorised this combined sample size as “anecdotal” (i.e., ≤5 groups or ≤10 group-years), “limited” (i.e., 6–15 groups or 11–30 group-years), or “substantial” (i.e., ≥16 groups or ≥31 group-years). For species with conflicting evidence regarding the extent of reproductive skew (see section 2.4.1), we only considered the sample size of the studies used for our final classification of the species.

## 3. Results

### 3.1. Mammals

We found a total of 140 studies with relevant data on maternity of the 41 mammal species examined (a median of 3 studies per species; range: 0–9 studies per species). Eleven of these species were excluded from our classification. Three species were excluded due to a lack of evidence of alloparental care (Table S1 and Figure 2), and eight species were excluded because we could not find data on maternity in wild populations or confirm the occurrence of multi-female groups in these species (Table S1 and Figure 2). The datasets that originally classified these eleven species as cooperative breeders did not include supporting references for their classification (N = 4 species) or we were unable to verify the cited information in the references provided (N = 7 species).

#### 3.1.1. Extreme female reproductive skew in mammals

For the remaining 30 mammal species, we found evidence for extreme female reproductive skew and alloparental care in 11 species (Table S1 and Figure 2). Yet, only two of the species that we classified as having extreme female reproductive skew were classified based on substantial sample size from multi-female groups (Tables 1 and S1).

### 3.2. Birds

We found 109 studies with relevant data on the parentage of the 37 bird species examined (a median of 3 studies per species; range: 1–6). Eight and six out of these 37/36 bird species were excluded from our classification of female/male reproductive skew, respectively. Six and three species were excluded as not relevant to the question of reproductive skew, respectively (mostly due to a lack of evidence of alloparental care by sexually mature female/male helpers; see Figures 3-4 and Table S2 for details). Two and three additional species were excluded because we could not find data on maternity/paternity in wild populations (Figures 3-4 and Table S2) and we were unable to verify the information cited by the datasets that classified these species as cooperative breeders.

#### 3.2.1. Extreme female reproductive skew in birds

For the remaining 29 bird species in our female dataset, we found evidence for alloparental care and extreme female reproductive skew in 12 bird species (including splendid fairy-wren in which there was no extreme reproductive skew in another population; Figure 3 and Table S2). Only three species that we classified as having extreme female reproductive skew were classified based on a substantial combined sample size from multi-female groups (Tables 2 and S2).

#### 3.2.2. Extreme male reproductive skew in birds

For the remaining 30 bird species in the male dataset, we found evidence of extreme male reproductive skew and alloparental care in 8 bird species (including white-winged chough for which extreme reproductive skew was dependent on whether it was a newly established group or not; Figure 4 and Table S2). Only two of the species that we classified as exhibiting extreme male reproductive skew were classified based on a substantial combined sample size from multi-male groups (Tables 3 and S2).

## 4. Discussion

This study revisits the underlying biological assumption of restrictive definitions of cooperative breeding. Namely, we tested whether there is a quantitative distinction between the reproductive skew exhibited by communal and cooperative breeding species [22; Figure 1]. After accounting for biases in how reproductive skew was assessed in previous datasets, we report two main findings. First, even among the cooperative mammal and bird species classified as having extreme reproductive skew, reproductive sharing occurs along a wide range. This range of reproductive sharing in these so-called “cooperative breeding” species largely overlaps with the scope of reproductive sharing among so-called “communal breeding” species (Figure 1). Second, depending on the rigour of the data, only 2-11 mammal and 1-12 bird species would qualify as cooperative breeders according to the restrictive definition of cooperative breeding. Below, we discuss the implications of these findings.

### 4.1. Ignoring group composition when measuring proxies of reproductive skew results in empirical and logical faults

(1) Frequently used datasets on mammalian and avian social systems classified 41 mammal species and 37 bird species as exhibiting alloparental care and an extreme within-group reproductive skew to the degree of virtual breeding monopolisation by a single female and/or male [12,16,21,22]. However, proxies of reproductive skew in these datasets were based on all sampled groups regardless of whether reproduction could, by definition, be skewed in them or not. Nevertheless, including groups in which reproductive skew is meaningless (e.g., single-female groups) is contrary to the fundamental rationale of within-group reproductive skew [16,23–25,28] and to how it is empirically measured [e.g., 36,41–45].

(2) Ignoring the social composition of groups when measuring proxies of reproductive skew may be problematic at the logical and empirical levels. At the logical level, including single-female and/or single-male groups in datasets may undermine the rationale of studies examining the causes or consequences of reproductive skew. The restrictive definition of cooperative breeding links between alloparental care and extreme reproductive skew by requiring alloparental care to be provided by potentially, but not actually, reproducing individuals [e.g., 12,13]. Nevertheless, this combination of alloparental care by non-reproducing female/male helpers is absent in single-female / single-male groups [16,25]. For example, Raihani & Clutton-Brock [21] tested whether viviparity (i.e., the maintenance of developing embryos inside the female’s reproductive tract) reduces the prevalence of reproductive suppression of subordinate females. To this end, they compared the prevalence of female reproductive monopolisation in mammals (i.e., viviparous species) versus birds (i.e., oviparous species in which embryos develop outside the female body). However, regardless of whether species are viviparous or oviparous, “dominant” females in single-female groups cannot suppress female helpers’ reproduction because there are no other females in their group. Since cooperative species differ in the prevalence of groups with helpers [32], considering single-female groups in such an analysis may thus obscure the true relationship between reproductive suppression, viviparity and oviparity.

(3) At the empirical level, including single-female or single-male groups in reproductive skew proxies inflates the assessment of the overall reproductive share of dominant females/males (Figures 1-4). Indeed, such a sampling bias was evident in several cooperative-breeding species. For example, the female reproductive skew was considered extreme for golden lion tamarins [*Leontopithecus rosalia*:, 46] and African wild dogs [*Lycaon pictus*:, 47] before studies that focused on groups with multiple sexually mature females revealed shared maternity in these groups and, consequently, a significant lower skew at the species level [27,46]. This inflation is particularly problematic in populations with a high proportion of single-female and/or single-male groups [as it is the case in many populations of cooperatively breeding species; reviewed by 32], which will be classified as exhibiting extreme reproductive skew even if reproduction is shared within the few multi-female and/or multi-male groups existing in the population.

(4) Even the percentages reported here probably overestimate the reproduction output of dominant adults for the following reasons. First, in the absence of more accurate data for some species, we calculated skew proxies from mixed samples that may include single-female groups/single-male groups (Tables 1-3). Second, group members may not reproduce simultaneously in species that breed throughout the year. Short-term studies that only assessed the number of reproductive females/males during the study period [e.g., via excavating of mole-rats colonies:, 48,49], may thus overlook asynchronous reproductive sharing [50,51]. The skew-of-groups proxy is, therefore, prone to overestimation when it is based on short-term studies that do not assess the dominance rank of breeders.

**Table 1.** Certainty of the classification of mammal species as exhibiting extreme female reproductive skew or not. The number of species according to the type of sample and combined sample size based on which we classified each species (N = 30 species, see Table S1 for details on each species).

| Combined sample size: | Type of sample |  |
| --- | --- | --- |
|  | A mixed sample of groups with a single female and groups with multiple females | Only groups with multiple sexually mature females |
| anecdotal or limited<br>(i.e., 1–15 groups or 1–30 group-years) | 1 | 14 |
| Substantial<br>(i.e., ≥16 groups or ≥31 group-years) | 6 | 9 |

**Table 2.** Certainty of the classification of bird species as exhibiting extreme female reproductive skew or not. The number of species according to the type of sample and combined sample size on which we based each species’ classification (N = 29 species, yet the table includes 30 entries as splendid fairy-wren was considered twice according to the different populations with different degrees of reproductive skew; see Table S2 for details on each species).

| Combined sample size: | Type of sample |  |
| --- | --- | --- |
|  | A mixed sample of groups with a single female and groups with multiple females | Only groups with multiple sexually mature females |
| anecdotal or limited<br>(i.e., 1–15 groups or 1–30 group-years) | 2 | 9 |
| Substantial<br>(i.e., $\geq 16$ groups or $\geq 31$ group-years) | 7 | 12 |

**Table 3.** Certainty of the classification of bird species as exhibiting extreme male reproductive skew or not. The number of species according to the type of sample and combined sample size on which we based each species’ classification (N = 30 species, see Table S2 for details on each species).

| Combined sample size: | Type of sample |  |
| --- | --- | --- |
|  | A mixed sample of groups with a single male and groups with multiple males | Only groups with multiple sexually mature males |
| anecdotal or limited<br>(i.e., 1–15 groups or 1–30 group-years) | 1 | 11 |
| Substantial<br>(i.e., ≥16 groups or ≥31 group-years) | 5 | 13 |

### 4.2. Inflation of within-group reproductive skew results in overestimating the prevalence of species exhibiting extreme reproductive skew

(5) False-positive classifications of species as exhibiting extreme reproductive skew lead, in turn, to overestimating the number of species that are qualified as cooperative breeders by restrictive definitions. Indeed, after partly accounting for this bias, this study shortens the lists of species with evidence of alloparental care and extreme reproductive skew by at least 73% for mammals and 54% for birds, depending on the criteria of rigorous data.

(6) This significant change in the number and identity of species that can be classified as cooperative breeders according to restrictive definitions has two important consequences. First, there may be a need to repeat comparative studies that used previous datasets [see also 17]. Second, the very small number of cooperative breeding species according to restrictive definitions (2–11 in mammals and 1-17 in birds) reduces the ability to carry out statistically meaningful analyses, thereby limiting insights into the evolutionary causes and consequences of alloparental care [see also 33]. It, thus, also questions the practicality of the restrictive definitions of cooperative breeding.

(7) Our results demonstrate a continuum of within-group reproductive sharing even in the group of species that were claimed to exhibit extreme reproductive skew (Figure 1). These findings question the underlying biological assumption of restrictive definitions regarding a quantitative gap between the reproductive skew of communal and cooperative breeding species [Figure 1, 22]. At the same time, we provide the first systematic evidence supporting Sherman and colleagues [33] and Riehl’s [30,52] proposal that within-group reproductive sharing among cooperative-breeding species is better seen as a continuum, in which the few species with an extreme reproductive skew represent one end, rather than a qualitatively different category.

We, therefore, join the majority of behavioural ecologists [e.g., 1,2,53–58] and evolutionary anthropologists [e.g., 59,60] in support of a more inclusive approach that defines cooperative breeding as the provision of alloparental care regardless of the reproductive status of alloparents.

### 4.3. Curating datasets on cooperative breeding species

(8) This work seeks to facilitate the constant improvement of datasets on cooperative breeding species. We, therefore, provide estimations for the strength of the evidence supporting each species’ classification (Tables S1-S2). These estimations should direct research efforts towards species that require further study and facilitate statistical control for data uncertainty. The latter is particularly important due to the high proportion of species whose classification is based on mixed samples or limited sample sizes (Tables 1-3). Nonetheless, the issue of data uncertainty is often ignored by curators of similar datasets [17, see also 61].

(9) This study demonstrates that a comprehensive review of the primary literature, and fine filtering of those groups that are relevant to the question addressed in a given study, are no less crucial than the statistical analysis of this data. These laborious reviewing tasks require qualified readers and extensive time that should not be underestimated. We thus join Schradin [17] in suggesting that, in the absence of these resources, meta-analyses should be based on smaller and more accurate datasets, rather than larger but less accurate ones. If this suggestion were to be accepted, small high-quality datasets could later be merged into comprehensive ones. Thereby, ensuring a better understanding of cooperative breeding in the longer perspective.

## Acknowledgements

We are grateful to Carel van Schaik for insightful discussions about an earlier version of this manuscript.

## References

1. Stacey PB, Koenig WD. 1990 Introduction. In Cooperative Breeding in Birds: Long-term Studies of Ecology and Behavior (eds PB Stacey, WD Koenig), pp. ix–xviii. Cambridge: Cambridge University Press.

2. Jennions MD, Macdonald DW. 1994 Cooperative breeding in mammals. Trends Ecol Evol 9, 89–93. (doi:10.1016/0169-5347(94)90202-X)

3. Isler K, van Schaik CP. 2012 Allomaternal care, life history and brain size evolution in mammals. J Hum Evol 63, 52–63. (doi:10.1016/j.jhevol.2012.03.009)

4. Sargeant EJ, Wikberg EC, Kawamura S, Fedigan LM. 2015 Allonursing in white-faced capuchins (*Cebus capucinus*) provides evidence for cooperative care of infants. Behaviour 152, 1841–1869. (doi:10.1163/1568539X-00003308)

5. Koenig WD, Dickinson JL, editors. 2004 Ecology and evolution of cooperative breeding in birds. Cambridge: Cambridge University Press.

6. Ostreiher R. 1997 Food division in the Arabian babbler nest: adult choice or nestling competition? Behavioral Ecology 8, 233–238. (doi:10.1093/beheco/8.2.233)

7. Gilchrist JS, Russell AF. 2007 Who cares? Individual contributions to pup care by breeders vs non-breeders in the cooperatively breeding banded mongoose (*Mungos mungo*). Behav Ecol Sociobiol 61, 1053–1060. (doi:10.1007/s00265-006-0338-2)

8. Ben Mocha Y, Mundry R, Pika S. 2019 Joint attention skills in wild Arabian babblers (*Turdoides squamiceps*): A consequence of cooperative breeding? Proceedings of the Royal Society B: Biological Sciences 286:1900, 1–10. (doi:10.1098/rspb.2019.0147)

9. Heymann EW. 1990 Social behaviour and infant carrying in a group of moustached tamarins, *Saguinus mystax* (primates: Platyrrhini: Callitrichidae), on Padre Isla, Peruvian Amazonia. Primates 31, 183–196. (doi:10.1007/BF02380940)

10. Bishop JM, Jarvis JUM, Spinks AC, Bennett NC, O’Ryan C. 2004 Molecular insight into patterns of colony composition and paternity in the common mole-rat *Cryptomys hottentotus hottentotus*. Mol Ecol 13, 1217–1229. (doi:10.1111/j.1365-294X.2004.02131.x)

11. Townsend AK, Bowman R, Fitzpatrick JW, Dent M, Lovette IJ. 2011 Genetic monogamy across variable demographic landscapes in cooperatively breeding Florida scrub-jays. Behavioral Ecology 22, 464–470. (doi:10.1093/beheco/arq227)

12. Cornwallis CK, West SA, Davis KE, Griffin AS. 2010 Promiscuity and the evolutionary transition to complex societies. Nature 466, 969–972. (doi:10.1038/nature09335)

13. Blumstein DT, Armitage KB. 1999 Cooperative Breeding in Marmots. Oikos 84, 369–382.

14. Ligon DavidJ, Burt BrentD. 2004 Evolutionary origins. In Ecology and evolution of cooperative breeding in birds (eds WD Koenig, JL Dickinson), pp. 5–34. Cambridge: Cambridge University Press.

15. Clutton-Brock TH. 2006 Cooperative breeding in mammals. In Cooperation in primates and humans: mechanisms and evolution (eds PM Kappeler, CP van Schaik), pp. 173–189. Berlin Heidelberg, Germany: Springer-Verlag.

16. Federico V, Allainé D, Gaillard J, Cohas A. 2020 Evolutionary Pathways to Communal and Cooperative Breeding in Carnivores. Am Nat 195, 1037– 1055. (doi:10.1086/708639)

17. Schradin C. 2017 Comparative studies need to rely both on sound natural history data and on excellent statistical analysis. R Soc Open Sci 4, 1–4.

18. Ross CT, Jaeggi A V., Borgerhoff Mulder M, Smith JE, Smith EA, Gavrilets S, Hooper PL. 2020 The multinomial index: A robust measure of reproductive skew: The Multinomial Index. Proceedings of the Royal Society B: Biological Sciences 287:202020. (doi:10.1098/rspb.2020.2025)

19. Haydock J, Koenig WD. 2003 Patterns of Reproductive Skew in the Polygynandrous Acorn Woodpecker. Am Nat 162, 277–289.

20. Kutsukake N, Nunn CL. 2006 Comparative tests of reproductive skew in male primates: The roles of demographic factors and incomplete control. Behav Ecol Sociobiol 60, 695–706. (doi:10.1007/s00265-006-0213-1)

21. Raihani NJ, Clutton-Brock TH. 2010 Higher reproductive skew among birds than mammals in cooperatively breeding species. Biol Lett 6, 630– 632.

22. Lukas D, Clutton-Brock T. 2012 Cooperative breeding and monogamy in mammalian societies. Proceedings of the Royal Society B: Biological Sciences 279, 2151–2156. (doi:10.1098/rspb.2011.2468)

23. Pamilo P, Crozier RH. 1996 Reproductive skew simplified. Oikos 75, 533–535.

24. Tsuji K, Tsuji N. 1998 Indices of reproductive skew depend on average reproductive success. Evol Ecol 12, 141–152. (doi:10.1023/A:1006575411224)

25. Montgomery TM, Pendleton EL, Smith JE. 2018 Physiological mechanisms mediating patterns of reproductive suppression and alloparental care in cooperatively breeding carnivores. Physiol Behav 193, 167–178. (doi:10.1016/j.physbeh.2017.11.006)

26. Richardson DS, Jury FL, Blaakmeer K, Komdeur J, Burke T. 2001 Parentage assignment and extra-group paternity in a cooperative breeder: The Seychelles warbler (*Acrocephalus sechellensis*). Mol Ecol 10, 2263–2273. (doi:10.1046/j.0962-1083.2001.01355.x)

27. Spiering PA, Somers MJ, Maldonado JE, Wildt DE, Gunther MS. 2010 Reproductive sharing and proximate factors mediating cooperative breeding in the African wild dog (*Lycaon pictus*). Behav Ecol Sociobiol 64, 583–592. (doi:10.1007/s00265-009-0875-6)

28. Harrington F, Paquet P, Ryon J, Fentress J. 1982 Monogamy in wolves: a review of the evidence. In Wolves of the world: perspectives of behavior, ecology, and conservation (eds FH Harrington, PC Paquet), pp. 209–222. Park Ridge, New Jersey: Noyes Publications. (doi:10.1016/0376-6357(84)90073-1)

29. Lacey E, Sherman PW. 1997 Cooperative Breeding in Naked Mole-Rats: Implications for Vertebrate and Invertebrate Sociality. In Cooperative breeding in mammals (eds N Solomon, J French), pp. 267–301. New York: Cambridge University Press.

30. Riehl C. 2013 Evolutionary routes to non-kin cooperative breeding in birds. Proceedings of the Royal Society B: Biological Sciences 280.1772, 1–7. (doi:10.1098/rspb.2013.2245)

31. Hackländer K, Möstl E, Arnold W. 2003 Reproductive suppression in female Alpine marmots, Marmota marmota. Anim Behav 65, 1133–1140. (doi:10.1006/anbe.2003.2159)

32. Ben Mocha Y, Scemama de Gialluly S, Markman S. In press. What is cooperative breeding in mammals and birds? Removing definitional barriers for comparative research. Major revisions in Biological Reviews

33. Sherman PW, Lacey EA, Reeve HK, Keller L. 1995 Forum: The eusociality continuum. Behavioral Ecology 6, 102–108. (doi:10.1093/beheco/6.1.102)

34. Warrington MH, Rollins LA, Russell AF, Griffith SC. 2015 Sequential polyandry through divorce and re-pairing in a cooperatively breeding bird reduces helper-offspring relatedness. Behav Ecol Sociobiol 69, 1311–1321. (doi:10.1007/s00265-015-1944-7)

35. Digby LJ, Barreto CE. 1993 Social organization in a wild population of *Callithrix jacchus*. I. Group composition and dynamics. Folia Primatologica 61, 123–134.

36. Townsend AK, Clark AB, McGowan KJ, Lovette IJ. 2009 Reproductive partitioning and the assumptions of reproductive skew models in the cooperatively breeding American crow. Anim Behav 77, 503–512. (doi:10.1016/j.anbehav.2008.10.030)

37. Temple HJ, Hoffman JI, Amos W. 2009 Group structure, mating system and extra-group paternity in the co-operatively breeding White-breasted Thrasher *Ramphocinclus brachyurus*. Ibis 151, 99–112. (doi:10.1111/j.1474-919X.2008.00867.x)

38. Ben Mocha Y, Mundry R, Pika S. 2018 Why hide? Concealed sex in dominant Arabian babblers (*Turdoides squamiceps*) in the wild. Evolution and Human Behavior 39, 575–582. (doi:10.1016/j.evolhumbehav.2018.05.009)

39. Cant MA. 2000 Social control of reproduction in banded mongooses. Anim Behav 59, 147–158. (doi:10.1006/anbe.1999.1279)

40. Bezerra BM, Souto ADS, Schiel N. 2007 Infanticide and Cannibalism in a Free-Ranging Plurally Breeding Group of Common Marmosets (*Callithrix Jacchus*). Am J Primatol 69, 945–952. (doi:10.1002/ajp)

41. Williams DA. 2004 Female control of reproductive skew in cooperatively breeding brown jays (*Cyanocorax morio*). Behav Ecol Sociobiol 55, 370–380. (doi:10.1007/s00265-003-0728-7)

42. Jamieson IG. 1997 Testing reproductive skew models in a communally breeding bird, the pukeko, Porhyrio porphyrio. Proceedings of the Royal Society B: Biological Sciences 264, 335–340. (doi:10.1098/rspb.1997.0048)

43. Heinsohn R, Dunn P, Legge S, Double M. 2000 Coalitions of relatives and reproductive skew in cooperatively breeding white-winged choughs. Proceedings of the Royal Society B: Biological Sciences 267, 243–249. (doi:10.1098/rspb.2000.0993)

44. Dugdale HL, Macdonald DW, Pope LC, Johnson PJ, Burke T. 2008 Reproductive skew and relatedness in social groups of European badgers, *Meles meles*. Mol Ecol 17, 1815–1827. (doi:10.1111/j.1365-294X.2008.03708.x)

45. Creel SR, Waser PM. 1991 Failures of reproductive suppression in dwarf mongooses (*Helogale parvula*): Accident or adaptation? Behavioral Ecology 2, 7–15. (doi:10.1093/beheco/2.1.7)

46. Baker AJ, Dietz JM, Kleiman D. 1993 Behavioural evidence for monopolization of paternity in multi-male groups of golden lion tamarins. Anim Behav 46, 1091–1103.

47. Girman DJ, Mills MGL, Geffen E, Wayne RK. 1997 A molecular genetic analysis of social structure, dispersal, and interpack relationships of the African wild dog (*Lycaon pictus*). Behav Ecol Sociobiol 40, 187–198. (doi:10.1007/s002650050332)

48. Sichilima AM, Faulkes CG, Bennett NC. 2008 Field evidence for aseasonality of reproduction and colony size in the Afrotropical giant mole-rat *Fukomys mechowii* (Rodentia: Bathyergidae). Afr Zool 43, 144–149. (doi:10.1080/15627020.2008.11657231)

49. Yeboah S, Dakwa KB. 2002 Aspects of the feeding habits and reproductive biology of the Ghana mole-rat *Cryptomys zechi* (Rodentia, Bathyergidae). Afr J Ecol 40, 110–119. (doi:10.1046/j.1365-2028.2002.00326.x)

50. Garber PA, Porter LM, Spross J, Di Fiore A. 2016 Tamarins: Insights into monogamous and non-monogamous single female social and breeding systems. Am J Primatol 78, 298–314. (doi:10.1002/ajp.22370)

51. Savage A, Giraldo LH, Soto LH, Snowdon CT. 1996 Demography, Group Composition, and Dispersal in Wild Cotton-Top Tamarin (*Saguinus oedipus*) Groups. Am J Primatol 38, 85–100.

52. Riehl C. 2017 Kinship and incest avoidance drive patterns of reproductive skew in cooperatively breeding birds. American Naturalist 190, 774–785. (doi:10.1086/694411)

53. Cockburn A. 1998 Evolution of helping behavior in cooperatively breeding birds. Annu Rev Ecol Syst 29, 141–177.

54. Brown JL. 1974 Alternate routes to sociality in jays - with a theory for the evolution of altruism and communal breeding. Am Zool 14, 63–80. (doi:10.1093/icb/14.1.63)

55. Kimball RT, Parker PG, Bednarz JC. 2003 Occurrence and evolution of cooperative breeding among the diurnal raptors (Accipitridae and Falconidae). Auk 120, 717–729. (doi:10.2307/4090102)

56. Jetz W, Rubenstein DR. 2011 Environmental Uncertainty and the Global Biogeography of Cooperative Breeding in Birds. Current Biology 21, 72–78. (doi:10.1016/j.cub.2010.11.075)

57. Covas R, Doutrelant C. 2019 Testing the Sexual and Social Benefits of Cooperation in Animals. Trends Ecol Evol 34, 112–120. (doi:10.1016/j.tree.2018.11.006)

58. Griesser M, Drobniak SM, Nakagawa S, Botero CA. 2017 Family living sets the stage for cooperative breeding and ecological resilience in birds. PLoS Biol 15(6), e2000483. (doi:10.1371/journal.pbio.2000483)

59. Burkart JM, Hrdy SB, van Schaik CP. 2009 Cooperative breeding and human cognitive evolution. Evol Anthropol 18, 175–186. (doi:10.1002/evan.20222)

60. Tomasello M, Gonzales-Cabrera I. 2017 The role of ontogeny in the evolution of human cooperation. Human Nature 28, 274–288. (doi:10.1007/s12110-017-9291-1)

61. Griesser M, Suzuki TN. 2016 Occasional cooperative breeding in birds and the robustness of comparative analyses concerning the evolution of cooperative breeding. Zoological Lett 2, 1–11. (doi:10.1186/s40851-016-0041-8)

