## Supplementary material for "Evidence for a reproductive sharing continuum in cooperatively breeding mammals and birds: consequences for comparative research": Table S1

| **Table 1. Within-group female reproductive sharing in the 41 mammal species classified as exhibiting alloparental care and an extreme female reproductive skew in Raihani & Clutton-Brock** (2010) **and/or Lukas & Clutton-Brock** (2012)**, and/or Federico *et al*.** (2020)**.** | | | | | | | |
| --- | --- | --- | --- | --- | --- | --- | --- |
| **Species** | **Sample:**  Only social groups with multiple sexually mature females /  A mixture of single-female and multi-female groups | **Percentage of offspring produced by the most dominant female in the social group**  +  (Type of evidence:  Genetic/ Behavioural/ Physiological) | **Percentage of social groups with a single breeding female**  +  (Type of evidence:  Genetic/ Behavioural/ Physiological) | **Female reproductive sharing:**  Extreme reproductive skew /  Not extreme reproductive skew | **Combined sample size:**  *Anecdotal* (≤5 groups or ≤10 group-years);  *Limited* (6–15 groups or 11–30 group-years);  *Substantial* (≥16 groups or ≥31 group-years) | **Remarks** | **Data sets that classified the species as exhibiting alloparental care and an extreme female reproductive skew** |
| **Oldfield mouse**  (*Peromyscus polionotus*) | **Not relevant**  We were unable to find evidence for alloparental care in this species (see also Margulis *et al.*, 2005). In addition, only 3.6% of burrows are occupied by more than two adults (n = 520 borrows: Foltz, 1981, p. 667). Namely, there are virtually no social groups with multiple adults in this species. | | | | | | - Lukas & Clutton-Brock (2012) |
| **California mouse**  (*Peromyscus californicus*) | **Not relevant**  California mice live in pairs and exhibit biparental care (Ribble, 2003). In addition, most offspring disperse before the next litter is whelped (Ribble, 1992, 2003). Accordingly, there are virtually no social groups with multiple adults in this species. | | | | | | - Lukas & Clutton-Brock (2012) |
| **Red** **wolf**  (*Canis rufus*) | A mixture of single-female and multi-female groups | 100%  (N = 174 litters produced by 90 pairs)  (Sparkman *et al.*, 2012, p. 1191) ^Genetic evidence^ | 100%  (N = 174 litters produced by 90 pairs)  (Sparkman *et al.*, 2012, p. 1191) ^Genetic evidence^ | **Not relevant ^1^** | **Substantial** | **^1^** No direct evidence of alloparental care in this species (Sparkman *et al.*, 2011). | - Federico *et al*. (2020) |
| **Eurasian beaver**  (*Castor fiber*) | **We are not aware of systematic data on reproductive sharing.**  Most social units consist of pairs only. A few social units consist of a pair and the offspring from the previous litter (most of them are not sexually mature). We are not aware of a study examining groups with more than one sexually mature female (see also Campbell *et al.*, 2005; Syrůčková *et al.*, 2015; Shavadze, 2016). Yet, see Sun (2003, p. 142) for evidence of plural breeding of multiple females within social groups of this species. | | | | | | - Lukas & Clutton-Brock (2012) |
| **Ochre mole-rat**  (*Fukomys ochraceocinereus*) | **We are not aware of systematic data on reproductive sharing.** | | | | | | - Lukas & Clutton-Brock (2012) |
| **African brush-tailed porcupine**  (*Atherurus africanus*) | **We are not aware of systematic data on reproductive sharing.** | | | | | | - Lukas & Clutton-Brock (2012) |
| **Side-striped jackal**  (*Canis adustus*) **^1^** | Not available **^2^** | Not available **^2^** | Not available **^2^** | Not available **^2^** | Not available **^2^** | **^1^** The taxonomic classification of the population studied by Moehlman (1983) has been changed from *Canis aureus* to *Canis adustus* since her study. Yet, see also evidnce for plural breeding in *Canis aureus* in (Pecorella *et al.*, 2023, pp. 42–43).  **^2^** Moehlman (1983, p. 430) observed only two families (N = 3 group-years) that breed while having a female helper (i.e. multi-female groups). Most helpers in this species and all female helpers in Moehlman’s (1983) study were offspring from the previous year (but see Loveridge & Macdonald (2001) for a report on a group with a few 1–2 year-old helpers). Although these offspring from the previous year may be physically sexually mature, we did not consider them as having breeding opportunities for the following reason: in Moehlman’s research site (Serengeti, Tanzania), the whelping season lasts from December to March (Moehlman, 1983). Since the gestation period of side-striped jackals lasts two months (Loveridge & Macdonald, 2001), the breeding season begins in October. Side-striped jackals reach sexual maturity at the age of 11–12 months old (Moehlman, 1983; Bingham & Purchase, 2003). Hence, even the pups born in the earliest part of the breeding season (i.e. December) would reach sexual maturity after the beginning of the next breeding season (i.e. November), when many partners and breeding territories would already be taken. See Bingham & Purchase (2002) for a similar timeline in Zimbabwe. | - Lukas & Clutton-Brock (2012) - Federico *et al*. (2020) |
| **Ethiopian dwarf mongoose**  (*Helogale hirtula*) | **We are not aware of systematic data on reproductive sharing.**  But see Kingdon (1988) for anecdotal observations of packs with multiple breeding females. | | | | | | - Lukas & Clutton-Brock (2012) - Federico *et al*. (2020) |
| **Black lion tamarin** (*Leontopithecus chrysopygus*) | **We are not aware of systematic data on reproductive sharing.**  Yet, the majority of social groups include a single sexually mature female (see group compositions in Valladares-Padua, 1993; Passos, 1994). | | | | | | - Lukas & Clutton-Brock (2012) |
| **Pied tamarin**  (*Saguinus bicolor*) | **We are not aware of systematic data on reproductive sharing.** | | | | | | - Lukas & Clutton-Brock (2012) |
| **Red-handed tamarin**  (*Saguinus midas*) | **We are not aware of systematic data on reproductive sharing.** | | | | | | - Lukas & Clutton-Brock (2012) |
| **Black-tufted marmoset**  (*Callithrix penicillata*) | Only groups with multiple sexually mature females | Not available | 0%  (N = 1 group)  (Decanini & Macedo, 2008, pp. 636–637) ^Behavioural evidence^ | **Not extreme reproductive skew** | **Anecdotal** |  | - Lukas & Clutton-Brock (2012) |
| **Coyote**  (*Canis latrans*) | Only groups with multiple sexually mature females | 40–60% **^1^**^,^ **^2^**  (N = 5 offspring in 1 group)  (Hennessy, 2007, pp. 66–76; Hennessy, Dubach & Gehrt, 2012, p. 736) ^Genetic evidence^ | 0% **^1^**^,^ **^2^**  (N = 1 group)  (Hennessy, 2007, pp. 66–76; Hennessy *et al.*, 2012, p. 736) ^Genetic evidence^  67%  (N = 3 groups)  (Hatier, 1995, pp. 25–29) ^Behavioural evidence^ | **Not extreme reproductive skew** | **Anecdotal** | **^1^** Group composition is given in Hennessy (2007). There were two additional social groups with two adult females in each group, but the only adult male in these groups was the father of one of these females. In total, the study genotyped 18 litters from 17 group-years; 16 litters were born to a single pair. Yet, estimations of the social composition of these groups may miss the presence of additional sexually mature females and, thus, underestimate the prevalence of female breeding monopolisation (i.e. some of the groups with a single female may include additional females that were not observed and did not breed).  **^2^** Two litters, one litter of two pups and one litter of three pups, born to different mothers sharing a den. Since the dominance rank of the mothers was unknown, we present the range of possibilities (i.e. a scenario in which the dominant female produced the litter of two pups and a scenario in which the dominant female produced the litter of three pups). | - Lukas & Clutton-Brock (2012) - Federico *et al*. (2020) |
| **Meerkat**  (*Suricata suricatta*) | Only groups with multiple sexually mature females | Not available | 79%  (N = 14 group-years)  (Clutton-Brock *et al.*, 1999) ^Behavioural evidence^  0% **^2^**  (N = 4 groups)  (O’Riain *et al.*, 2000, p. 474) ^Behavioural evidence^ | **Not extreme reproductive skew ^5^** | **Substantial** | **^1^** Most probably, the same groups were examined across multiple years and the number of group-years is, therefore, probably higher than 23.  **^2^** In groups with multiple sexually mature females and a male unrelated to all females.  **^3^** The total sample size is 260 litters with at least one pup that survived to emergence from the den (61 litters of subordinant females + 199 litters of dominant females). Assuming that each of the 61 litters of subordinant females was from a group in which a dominant female also produced a litter with at least of surviving pup, the percentage of groups in which only the dominant female produced a surviving litter is 69% (61/199). Namely, in 61 groups there were litters by two females, and in 138 groups there was only a single litter by the dominant female.  **^4^** The parentage proxy can also be calculated from this sample under the following assumptions: (i) the total number of offspring by subordinant females that survived to independence is (63% of emergent pup reach independence * (3.4 emergent pups per litter * 61 litters with at least one surviving pup) = 130.  (ii) the total number of offspring by dominant females that survived to independence is (77.7% of emergent pup reach independence * (3.97 emergent pups per litter * 199 litters with at least one surviving pup) = 614. (iii) 614 surviving offspring of dominant females is 83% of total surviving offspring (614+130=744). All data is from (Clutton-Brock, Hodge & Flower, 2008, p. 694) ^Behavioural and physiological evidence^  **^5^** See also Doolan & MacDonald (1997, p. 841) ^Behavioural evidence^, Clutton-Brock *et al*. (2001), and Clutton-Brock *et al.* (2008).  We did not consider Griffin *et al*. (2003) because the study of Spong *et al*. (2008) is based on a larger sample that also includes the sample studied by Griffin *et al*. (2003). | - Raihani & Clutton-Brock (2010) - Lukas & Clutton-Brock (2012) - Federico et al. (2020) |
|  | A mixture of single-female and multi-female groups | 90% **^1^**  (N = 333 offspring in 23 groups)  (Spong *et al.*, 2008) ^Genetic evidence^ | 69% **^3^**  (N = 260 litters from which at least one pup emerged, assumed from 199 group-years) **^4^**  (Clutton-Brock *et al.*, 2008, p. 694) ^Behavioural and physiological evidence^ |  |  |  |  |
| **American beaver**  (*Castor canadensis*) | Only groups with multiple sexually mature females | Not available | 12%  (N = 8 group-years, 4 groups)  (Busher, Warner & Jenkins, 1983, pp. 315; 317) ^Physiological evidence^ | **Not extreme reproductive skew ^3^** | **Substantial** | **^1^** Most colonies of the species consist of a single pair (Svendsen, 1989; Crawford *et al.*, 2008). In addition, most of the colonies in this study did not have multiple sexually mature females (age distribution of colonies is given in Crawford, 2007); so 80% is probably an overestimation.  **^2^** The percentage of multi-female groups with a single breeding female can be estimated under the following assumptions.  ~16% of colonies in this population had multiple sexually mature females (Muller-Schwarzei & Schulte, 1999, p. 167). Since 16% of 35 groups is 5.6 groups, the three groups observed with multiple breeding females can be estimated to be 54% of the groups with multiple sexually mature females in the population (i.e., 46% of groups in the population had a single breeding female).  **^3^** See also Bergerud & Miller (1977) ^Physiological evidence^, Wheatley (1993) ^Physiological evidence^, and Fischer *et al.* (2010) ^Physiological evidence^ for several cases of multiple lactating females in a common den. See Sun (2003, p. 142) ^Behavioural evidence^ for a review of a long-term study population in New York with evidence of plural breeding by females within the same group. | - Lukas & Clutton-Brock (2012) |
|  | A mixture of single-female and multi-female groups |  | 80% **^1^**  (N = 15 groups)  (Crawford *et al.*, 2008, pp. 577, 579) ^Physiological and genetic evidence^  91% **^2^**  (N = 35 groups)  (Sun, 2003, p. 142) ^Behavioural evidence^ |  |  |  |  |
| **Red fox**  (*Vulpes vulpes*) | Only groups with multiple sexually mature females | 38% **^1^**^,^ **^2^**  (N = 42 surviving offspring in 5 groups)  (Baker *et al.*, 2004, pp. 770–772) ^Genetic evidence^ | 17%  (N = 23 group-years, 22 groups)  (Converse, 2012, pp. 36–44) ^Genetic evidence^  25% **^2^**  (N = 8 group-years, 5 groups)  (Baker *et al.*, 2004, pp. 770–772) ^Genetic evidence^  29%  (N = 7 group-years)  (Zabel & Taggart, 1989, pp. 833–834) ^Behavioural evidence^ | **Not extreme reproductive skew** | **Substantial** | **^1^** We excluded group S1992 because the subordinate female was not genetically sampled, although she did breed.  **^2^** No difference between the survival rate of dominant and subordinate females’ offspring. | - Federico *et al*. (2020) |
| **Striped mouse**  (*Rhabdomys pumilio*) | Only groups with multiple sexually mature females | Not available | 20%  (N = 10 group-years, 8 groups)  (Schradin & Pillay, 2004, pp. 41–42) ^Behavioural and physiological evidence^ | **Not extreme reproductive skew** | **Limited ^1^** | **^1^** But polygyny is common in this species (Schubert, Pillay & Schradin, 2009; Schradin, Schneider & Lindholm, 2010; Schradin, Pillay & Bertelsmeier, 2019). | - Lukas & Clutton-Brock (2012) |
| **Arctic fox**  (*Vulpes lagopus*) | Only groups with multiple sexually mature females | Not available | 20% **^1^**  (N = 15 groups)  (Norén *et al.*, 2012, pp. 1106–1112) ^Genetic evidence^  <28% **^2^**  (N = 39 group-years)  (Goltsman *et al.*, 2005, p. 409) ^Behavioural evidence^  50% **^3^**  (N = 2 groups)  (Strand *et al.*, 2000, pp. 226–227) ^Physiological evidence^  100%  (N = 2 groups)  (Cameron, Berteaux & Dufresne, 2011, pp. 1366; 1369) ^Genetic evidence^ | **Not extreme reproductive skew ^4^** | **Substantial** | **^1^** We could infer maternity in only 15 groups. The calculated percentage (20%) is conservative and may overestimate reproductive monopolisation because singular breeding in some groups was inferred based on limited evidence (e.g. in den “Juvdalskamper 2000”).  **^2^** The sample size of 39 group-years consists of groups with more than two adults, and in most or all of them, there were multiple sexually mature females.  **^3^** Three families consisting of a pair and an additional adult reproduced. At least two of these families included multiple sexually mature females (den A in 1988 and den A in 1990). The sex of the additional adult in the third family (potentially Dan B in 1988) is not indicated. And this third family was therefore excluded from our anaylsis. We considered den A 1988 to have shared maternity and den A 1990 to have a single breeding female.  **^4^** Despite the results from Cameron *et al.* (2011), the species was classified as exhibiting no extreme reproductive skew. This is due to studies with greater sample sizes that are based on genetic evidence.  See also Carmichael *et al*. (2007, p. 341) for further evidence of plural breeding in one social group, but without indication of the social composition of the other groups studied (N = 8).  See White (1992, p. 36) for evidence of at least three cases of communal denning by multiple females (>11%), but the exact social composition of each social group in the sample is not indicated (N = 28 den-years with more than two adults).  See Tannerfeldt *et al*. (2003, p. 175) and Norén *et al*. (Norén *et al.*, 2017, p. 9) for reviews of plural breeding in several locations, and Kruchenkova & Formozov (1995, p. 19) for the usual pattern of plural breeding in Mednyi (Copper) Island. | - Lukas & Clutton-Brock (2012) |
| **Prairie vole**  (*Microtus ochrogaster*) | Only groups with multiple sexually mature females | Not available | 21%  (N = 123 groups)  (Getz & McGuire, 1997, p. 530) ^Physiological evidence^  54% **^1^**  (N = 39 groups)  (Getz & McGuire, 1997, p. 530) ^Physiological evidence^ | **Not extreme reproductive skew ^2^** | **Substantial** | **^1^** Groups with multiple sexually mature females and unrelated individuals.  **^2^** See also McGuire, Getz & Oli (2002) and Solomon & Keane (2018, p. 198) ^Genetic evidence^. | - Lukas & Clutton-Brock (2012) |
| **African wild dog**  (*Lycaon pictus*) | Only groups with multiple sexually mature females | 66.7%  (N = 18 surviving offspring, 7 group-years, 6 groups)  (Spiering *et al.*, 2010, p. 586) ^Genetic evidence^ | 40% ^1^  (N = 10 group-years, 6 groups)  (Spiering *et al.*, 2010, p. 586) ^Genetic evidence^  25% **^2^**  (N = 4 group-years, 3 groups)  (Malcolm & Marten, 1982, pp. 2–3) ^Behavioural evidence^ | **Not extreme reproductive skew ^3^** | **Limited** | **^1^** Group-year Thanda 2007 was excluded as no data on pregnancy and maternity was available.  **^2^** In one of the groups with two reproducing females, the dominant female’s litter died shortly after the birth, while the subordinate female’s litter survived. The next year, the dominant female’s litter survived, while the subordinate female’s litter failed soon after the birth.  **^3^** See also Creel *et al.* (1997, pp. 303–304) ^Behavioural evidence^ for evidence that dominant females reproduce in 81.5% ± 7.0% pack-years (N = 22 litters in 27 pack-years; 11 packs), while sexually mature subordinate females reproduced in 10.4% ± 3.7% pack-years (N = 7 litters by 7 subordinate females during 67 individual-years).  See Burrows (1995, p. 405) for evidence of beta females producing 25% of litters (N = 57 litters and at least 14 beta females). Both studies are probably based on mixed samples. | - Raihani & Clutton-Brock (2010) - Lukas & Clutton-Brock (2012) - Federico *et al*. (2020) |
| **Wolf**  (*Canis lupus*) | Only groups with multiple sexually mature females | Not available | 33%  (N = 3 groups)  (Peterson *et al.*, 2002, p. 1407) ^Behavioural evidence^  44% **^1^**  (N = 9 group-years, 2 groups)  (Haber, 1977, pp. 199–203; 231; 254–259) ^Behavioural evidence^  50%  (N = 2 group-years, 1 group)  (Murie, 1944, pp. 24–40) ^Behavioural evidence^ | **Not extreme reproductive skew ^2^** | **Limited** | **^1^** Females in their third year of life were considered sexually mature.  **^2^** See also Harrington *et al.* (1982). See (Mech & Nelson, 1989, p. 676) ^physiological evidence^ for a case report of polygeny in one group.  Most social groups of the species include a breeding pair and its offspring, which usually disperse upon sexual maturity in the third year of life (Mech, 1999; Mech, Wolf & Packard, 1999). | - Lukas & Clutton-Brock (2012) - Federico *et al*. (2020) |
| **Common marmoset**  (*Callithrix jacchus*) | Only groups with multiple sexually mature females | 33%  (N = 3 surviving offspring in 1 group)  (Bezerra, Souto & Schiel, 2007, pp. 948, 950) ^Behavioural evidence^  67–83% **^1^**  (N = 12 surviving offspring in 3 groups)  (Digby & Barreto, 1993, pp. 126–127; Nievergelt *et al.*, 2000, p. 12) ^Behavioural and genetic evidence^ | 33% **^2^**  (N = 3 groups)  (Digby & Barreto, 1993, pp. 126–127; Nievergelt *et al.*, 2000, p. 12) ^Behavioural and genetic evidence^  50%  (N = 2 groups with at least one lactating female)  (Scanlon, Chalmers & Monteiro da Cruz, 1988, pp. 298–300) ^Behavioural and physiological evidence^  66.7% **^3^**  (N = 3 groups)  (Lazaro-Perea *et al.*, 2000, pp. 139–141) ^Behavioural evidence^ | **Not extreme reproductive skew ^4^** | **Limited** | **^1^** The mother of two surviving offspring in one of the groups (group A) is unknown. Hence, the range of percentages represents the two extreme possibilities; namely, that both offspring were born to the subordinate female (67%) or that they were both born to the dominant female (83%).  **^2^** In each of the three groups, two females gave birth, but in only two groups, both females had at least one surviving offspring.  **^3^** This was a big group that went through three social changes during which the number of females and dominance order changed. It is, thus, not certain who was the dominant female at the time of birth.  **^4^** See also Digby & Ferrari (1994, p. 394) ^Behavioural evidence^ for a review of shared maternity in several study sites.  See Arruda *et al*. (2005) for evidence of six unsuccessful breeding attempts of six subordinate females in four groups and an extended account of an additional group in which successful polygyny existed for at least four years (reported by Digby & Barreto, 1993 as group ‘A’ / ‘Plantacao’). Yet, Arruda *et al*. (2005) deliberately focus on groups with unsuccessful polygyny, and it is not clear whether all successful breeding attempts by dominant females occurred in the presence of multiple sexually mature females. | - Lukas & Clutton-Brock (2012) |
| **Mongolian gerbil**  (*Meriones unguiculatus*) | A mixture of single-female and multi-female groups | Not available | >50% **^1^**  (N = 17 groups)  (Wang *et al.*, 2011, p. 557) ^Genetic evidence^ | **Not extreme reproductive skew ^2^** | **Substantial** | **^1^** This estimation is based on the report of 2.8 ± 2.2 (mean ± SD) breeding pairs in a group.  **^2^** See also Liu *et al.* (2009). See Gromov (2021, p. 18) for a review showing the frequent occurrence of polygyny (<33% of groups) in several populations. | - Lukas & Clutton-Brock (2012) |
| **Golden lion tamarin** (*Leontopithecus rosalia*) | Only groups with multiple sexually mature females | 67% **^1^**  (N = 38 offspring, 19 groups)  (Dietz & Baker, 1993, pp. 1071–1072) ^Behavioural evidence^  73% **^2^**  (N = 15 births in 5 groups)  (Henry, 2011, p. 172; Henry *et al.*, 2013, pp. 677; 679) ^Physiological evidence^ | 60% **^2^**  (N = 5 groups with infants born alive)  (Henry, 2011, p. 172; Henry *et al.*, 2013, pp. 677; 679) ^Physiological evidence^  73.8% **^3^**  (N = 61 group-years with living offspring)  (Baker, Bales & Dietz, 2002, p. 202) ^Behavioural evidence^  77%  (N = 90 group-years)  (Dietz & Baker, 1993, p. 1070) ^Behavioural evidence^ | **Not extreme reproductive skew ^4^** | **Substantial** | **^1^** No significant difference between the number of surviving offspring (per sample period, per female) of subordinate (0.333 ± 0.617) versus dominant (0.684 ± 1.108) females in polygynous groups. 67% is a conservative estimation, as we assumed there is only one subordinate female in each group. If there were multiple subordinate females in a group, the reproductive skew would change in favour of subordinates: (0.684) / ([0.333 X # of subordinate females] + 0.684).  The sample size of groups was based on the statement that there were 19 dominant females in the sample.  In addition, 21% of natal females that reached sexual maturity and had no unrelated male in the group reproduced while their mother was present in the group (N = 24). 50% of natal females that reached sexual maturity in a group with an unrelated male reproduced while their mother was still present in the group (N = 4) (Dietz & Baker, 1993, p. 1071) ^Behavioural evidence^  **^2^** We excluded group “POR” as it is indicated as “No” for polygyny (Henry *et al.*, 2013, p. 677)  **^3^** This sample may partly overlap with Dietz & Baker’s (1993) sample.  **^4^** See also French *et al*. (2003, p. 1284) for evidence of multiple breeding females in the wild. | - Lukas & Clutton-Brock (2012) |
| **European badger**  (*Meles meles*) | Only groups with multiple sexually mature females | Not available | 50%  (N = 2 group-years, 1 group)  (Woodroffe, 1993, pp. 413–414) ^Behavioural evidence^ | **Not extreme reproductive skew ^1-3^** | **Substantial** | **^1^** The percentage is calculated from a sample of groups with offspring that survived to independence.  **^2^** Mean female reproductive skew was measured as *B* 0.039 (N = 24 groups), which is low, despite being significantly different from a random distribution of maternity (*B* ranges from -1 to 2, when 0 means that reproductive skew is randomly distributed Dugdale *et al.*, 2008). See also Woodroffe & Macdonald (2000) and Newman *et al*. (2011).  **^3^** Only a few populations seem to exhibit alloparental care (Woodroffe, 1993). In addition, evidence for alloparental care is limited (Woodroffe, 1993; Dugdale, 2007; Dugdale, Ellwood & Macdonald, 2010) and seems to have limited positive effect (Woodroffe & Macdonald, 2000; Dugdale *et al.*, 2010). | - Federico *et al*. (2020) |
|  | A mixture of single-female and multi-female groups |  | 63% **^1^**  (N = 167 group-years)  (Dugdale *et al.*, 2007, p. 5300) ^Genetic evidence^  51% **^2^**  (N = 79 group-years)  (Carpenter *et al.*, 2005, p. 278) ^Genetic evidence^ |  |  |  |  |
| **Dwarf mongoose**  (*Helogale parvula*) | A mixture of single-female and multi-female groups | 85%  (N = 39 offspring, 21 litters, 9 groups)  (Keane *et al.*, 1994) ^Genetic evidence^ | ≤67% **^1^**  (N = 9 groups)  (Keane *et al.*, 1994) ^Genetic evidence^  ≤67% **^2^**  (N = 30 litters, ~23 group-years)  (Rood, 1980, p. 145) ^Behavioural evidence^ | **Not extreme reproductive skew** | **Substantial** | **^1^** This may be an overestimation of reproductive monopolisation because the paper only indicates that at least three out of the nine groups had more than one pregnant female at the same time.  **^2^** This is a conservative estimation as no exact values are given, but at least 10 litters were born during simultaneous pregnancies. The number of group years is only an estimation because it is not indicated how many of the eight simultaneous litters occurred in the same group-year. Thus, the number of group-years is between 22 and 25 (i.e. 20 group-years with one litter + 1 group-year with two simultaneous litters + between 1 to 4 group-years with simultaneous litters). | - Lukas & Clutton-Brock (2012) - Federico *et al*. (2020) |
| **Mechow’s mole-rat**  (*Fukomys mechowii*) | A mixture of single-female and multi-female groups | Not available | 71%  (N = 14 groups, each with at least one breeding female)  (Sichilima, Faulkes & Bennett, 2008, p. 146) ^Physiological evidence^ | **Not extreme reproductive skew ^1^** | **Limited** | **^1^** See also Scharff *et al.* (2001, p. 1007) for inconclusive evidence of female plural breeding. | - Lukas & Clutton-Brock (2012) |
| **Golden-headed lion tamarin**  (*Leontopithecus chrysomelas*) | A mixture of single-female and multi-female groups **^1^** | Not available | 74.7% **^2^**  (N = 79 births during 51 group-years in 8 groups)  (Heslin Piper, 2015, pp. 19; 27; Heslin Piper, Dietz & Raboy, 2017, p. 175) ^Behavioural evidence^ | **Not extreme reproductive skew ^3^** | **Substantial** | **^1^** In a subset of this data, the percentage of groups with multiple sexually mature females was 53.6% (Raboy, 2002, p. 76). If this rate is used to estimate the prevalence of groups with multiple sexually mature females in the larger sample, then the percentage of multi-female groups with a single breeding female decreases to ~48%.  **^2^** Percentage is calculated based on the statement that “on 10 occasions, two females were observed concurrently reproducing in the same group” (Heslin Piper, 2015, p. 19). Namely, 20 births occurred within a few months of each other. In all 10 cases, one or both litters failed. In two cases, the younger female’s litter survived (the number of older females’ litter that survived is not indicated).  **^3^** See also de Vleeschouwer, van Elsacker & Leusa (2001) for evidence of multiple breeding females in a captive group. | - Lukas & Clutton-Brock (2012) |
| **Saddleback tamarin**  (*Saguinus fuscicollis*) | Only groups with multiple sexually mature females | 81–85% **^1^**  (N = 27 surviving offspring, 16 group-years, 7 groups)  (Goldizen *et al.*, 1996, pp. 62–64, 82–83) ^Behavioural evidence^  100%  (N = 2 surviving offspring, 1 group)  (Erb & Porter, 2020, pp. 3–4) ^Behavioural evidence^ | 75%  (N = 16 group-years with at least one breeding attempt with surviving offspring, 7 groups)  (Goldizen *et al.*, 1996, pp. 62–64, 82–83) ^Behavioural evidence^  100%  (N = 1 group with surviving offspring)  (Erb & Porter, 2020, pp. 3–4) ^Behavioural evidence^ | **Not extreme reproductive skew ^2^** | **Limited** | **^1^** It is not clear which female in the group “N 1982” had two surviving offspring. Hence, the range of percentages represents the two extreme possibilities. Namely, both offspring were born to the subordinate female (81%) or the dominant female (85%).  **^2^** Despite the data from Erb & Porter (2020), the species is classified as exhibiting no extreme reproductive skew in the light of other studies with greater sample sizes and that are based on the same type of evidence.  See also Calegaro-Marques, Bicca-Marques & Azevedo (1995) ^Behavioural evidence^ for anecdotal evidence of multiple breeding females in one group. | - Raihani & Clutton-Brock (2010) |
| **Woodland vole**  )*Microtus pinetorum*) | A mixture of single-female and multi-female groups | Not available | 83.5% **^1^**  (N = ~23 groups over 3 seasons)  (Solomon, Vandenbergh & Sullivan, 1998, p. 2133) ^Behavioural and physiological evidence^ | **Not extreme reproductive skew ^2^** | **Substantial** | **^1^** Sample size of groups with at least one reproductive female is calculated according to Table 2 (Solomon *et al.*, 1998, p. 2134).  **^2^** Most social units in this species are family units, with a single sexually mature female. See also FitzGerald & Madison (1983). | - Lukas & Clutton-Brock (2012) |
| **Cotton-top tamarin**  (*Saguinus Oedipus*) | Only groups with multiple sexually mature females | Not available | 92% **^2^**  (N = 52 litters in 50 group-years)  (Savage *et al.*, 2021, pp. 5–6) ^Behavioural evidence^ | **Extreme reproductive skew ^3^** | **Substantial** | **^1^** Although multiple females did not give birth simultaneously (simultaneous pregnancies resulted in only one female producing offspring or in the failure of both pregnancies), different females in the same group produced offspring in different years.  **^2^** Offspring that were born simultaneously to two females in the same group did not survive.  **^3^** Despite data from Savage *et al.* (Savage *et al.*, 1996), the species was classified as exhibiting extreme reproductive skew due to data from studies with greater sample sizes and samples of only multi-female groups.  See also Wheaton *et al*. (2022, p. 3) for evidence of subordinate females’ pregnancies. | - Lukas & Clutton-Brock (2012) |
|  | A mixture of single-female and multi-female groups | 89% **^1^**  (N = 19 surviving offspring, 5 group-years)  (Savage *et al.*, 1996, pp. 90–91) ^Behavioural evidence^ |  |  |  |  |  |
| **Naked mole-rat** (*Heterocephalus glaber*) | Only groups with multiple sexually mature females | Not available | 92%  (N = 25 group-years, 18 groups with a queen)  (Braude, 1991, pp. 75–76; 142–161) ^Physiological evidence^ | **Extreme reproductive skew ^1^** | **Substantial** | **^1^** See also Lacey & Sherman (1997, p. 276) ^Physiological evidence^ and Buffenstein *et al*. (2022) for reviews. | - Lukas & Clutton-Brock (2012) |
| **Damaraland mole-rat**  (*Cryptomys damarensis*) | A mixture of single-female and multi-female groups **^1^** | >94% **^2^**  (N = 376 offspring in 18 colonies, each with a breeding female in two sites)  (Burland *et al.*, 2004, p. 2374) ^Genetic evidence^ | 100%  (N = 18 colonies with a breeding female in two sites)  (Burland *et al.*, 2004, p. 2374) ^Physiological and genetic evidence^  100%  (N = 55 colonies)  (Young & Bennett, 2010, p. 3192) ^Physiological evidence^ | **Extreme reproductive skew ^3^** | **Substantial** | **^1^** All colonies had multiple females. However, since their age is not indicated, it is not clear whether all females were sexually mature.  **^2^** The identities of the mothers of the 6% of offspring that were not assigned to the dominant female are unclear.  **^3^** See also Jarvis & Bennett (1993) and Faulkes & Bennett (2009) for evidence of single female breeding as a rule in this species. | - Raihani & Clutton-Brock (2010) - Lukas & Clutton-Brock (2012) |
| **Ethiopian wolf**  (*Canis simensis*) | Only groups with multiple sexually mature females | 95%  (N = 21 offspring in 5 groups)  (Randall *et al.*, 2007, p. 584) ^Genetic evidence^ | 60%  (N = 5 groups)  (Randall *et al.*, 2007, p. 584) ^Genetic evidence^  94% **^1^**  (N = 17 group-years)  (Sillero-Zubiri, Gottelli & Macdonald, 1996, p. 334) ^Behavioural evidence^ | **Extreme reproductive skew ^2^** | **Limited** | **^1^** Sillero-Zubiri *et al.* (1996) suggest that the observed monogamy may be the result of a human-caused fragmented habitat.  **^2^** Despite the high percentage of groups with multiple breeding females (Randall *et al.*, 2007, p. 584), the species was classified as exhibiting extreme female reproductive skew in the light of the maternity data in (Randall *et al.*, 2007, p. 584) and the greater sample size of (Sillero-Zubiri *et al.*, 1996, p. 334). | - Raihani & Clutton-Brock (2010) - Lukas & Clutton-Brock (2012) - Federico *et al*. (2020) |
| **Alpine marmot**  (*Marmota marmota*) | Only groups with multiple sexually mature females | Not available | 100% **^1^**  (N ~ 29 group-years)  (Hackländer, Möstl & Arnold, 2003, pp. 1135–1137) ^Genetic evidence^ | **Extreme reproductive skew ^2^** | **Limited** | **^1^** Dominant females produced weaned offspring while living in groups with multiple sexually mature females in ~29 years (figure 1 in Hackländer *et al.*, 2003). But the exact number of these individual females is not indicated.  In addition, none of the 60 subordinate females observed during this long-term study produced weaned offspring. But the exact number of groups in which these females lived is not indicated.  **^2^** In 14 years of research, only once did a subordinate female produce surviving offspring. | - Raihani & Clutton-Brock (2010) |
| **Dhole**  (*Cuon alpinus*) | Only groups with multiple sexually mature females | Not available | 100% **^1^**  (N = 2 group-years, 1 group)  (Johnsingh, 1982, p. 446) ^Behavioural evidence^  50%  (N = 2 groups)  (Davidar, 1974, pp. 184–187) ^Physiological evidence^ | **Extreme reproductive skew ^2^** | **Anecdotal** | **^1^** It is not clear whether the same or different females bred during each of these two years.  **^2^** But see Cohen (1978) for further reports on multiple breeding females. | - Federico *et al*. (2020) |
|  | A mixture of single-female and multi-female groups |  | 100%  (N = 7 group-years, 2 groups)  (Venkataraman, 1998, pp. 679, 682) ^Behavioural evidence^ |  |  |  |  |
| **Black-backed jackal**  (*Canis mesomelas*) | Only groups with multiple sexually mature females | 100% **^1^**  (N = 1 surviving offspring)  (Moehlman, 1979, 1983, p. 428) ^Behavioural evidence^ | 100% **^1^**  (N = 1 group)  (Moehlman, 1979, 1983, p. 428) ^Behavioural evidence^ | **Extreme reproductive skew ^2^** | **Anecdotal** | **^1^** The sexually mature female helper was the daughter of the breeding pair and had no genetically unrelated male in the group.  **^2^** See Ferguson, Nel & Wet (1983, p. 499) and Nattrass *et al.* (2017, p. 12) for evidence of groups with multiple breeding females.  In addition, there is no evidence that black-backed jackals can breed before the age of one year (Ferguson *et al.*, 1983; Loveridge & Macdonald, 2001). Thus, offspring from the previous year, which are the typical helpers in this species (but see Loveridge & Macdonald, 2001), are not considered sexually mature helpers. | - Lukas & Clutton-Brock (2012) - Federico *et al*. (2020) |
| **Moustached tamarin**  (*Saguinus mystax*) | Only groups with multiple sexually mature females | 100%  (N = 5 surviving offspring, 1 group)  (Huck *et al.*, 2005, pp. 450, 455) ^Genetic evidence^ | 100%  (N = 1 group)  (Huck *et al.*, 2005, pp. 450, 455) ^Genetic evidence^ | **Extreme reproductive skew ^1^** | **Anecdotal** | **^1^** See Ramirez (1984) ^Behavioural evidence^, Smith *et al.* (2001) ^Behavioural evidence^, Löttker, Huck & Heymann (2004) ^Behavioural evidence^, and Culot *et al.* (2011) ^Behavioural evidence^ for further evidence of unsuccessful simultaneous breeding in multi-female groups. But see Ramirez (1989) ^Behavioural evidence^ for anecdotal evidence of multiple breeding females in one group. | - Lukas & Clutton-Brock (2012) |
| **Pygmy marmoset**  (*Cebuella pygmaea*) | Only groups with multiple sexually mature females | Not available | 100% **^1^**  (N = 2 groups)  (Soini, 1988, pp. 90; 103; 107) ^Behavioural evidence^ | **Extreme reproductive skew** | **Anecdotal** | **^1^** Based on the social composition of the groups observed (Table 3 in Soini, 1988) and the age of the first conception (Table 4 in Soini, 1988). | - Raihani & Clutton-Brock (2010) |
| **Common mole-rat**  (*Cryptomys hottentotus*) | A mixture of single-female and multi-female groups | 99–100%  (N < 176 surviving offspring from 13 colonies in 2 sites)  (Bishop *et al.*, 2004, p. 1222) ^Physiological and genetic evidence^ | 100%  (N = 13 colonies in 2 sites)  (Bishop *et al.*, 2004, p. 1224) ^Physiological and genetic evidence^ | **Extreme reproductive skew ^1^** | **Substantial** | **^1^** See also Faulkes & Bennett (2009), which provides evidence of single-female breeding as a rule in this species, with rare exceptions. | - Lukas & Clutton-Brock (2012) |
| **Ghana mole-rat**  (*Fukomys zechi*) | Only groups with multiple sexually mature females | Not available | 100%  (N = 3 colonies)  (Yeboah & Dakwa, 2002a, p. 89) ^Physiological evidence^ | **Extreme reproductive skew ^1^** | **Anecdotal** | **^1^** See also Yeboah & Dakwa (2002b) ^Physiological evidence^ for a larger sample, but which may include colonies with a single sexually mature female. | - Lukas & Clutton-Brock (2012) |

**References**

Arruda, M.F., Araújo, A., Sousa, M.B.C., Albuquerque, F.S., Albuquerque, A.C.S.R. & Yamamoto, M.E. (2005) Two breeding females within free-living groups may not always indicate polygyny: Alternative subordinate female strategies in common marmosets (*Callithrix jacchus*). *Folia Primatologica* **76**, 10–20.

Baker, A.J., Bales, K. & Dietz, J.M. (2002) Mating system and group dynamics in lion tamarins. In *Lion tamarins: biology and conservation* (eds D. Kleiman & A. Rylands), pp. 188–212. Smithsonian Institution Press, Washington DC.

Baker, P.J., Funk, S.M., Bruford, M.W. & Harris, S. (2004) Polygynandry in a red fox population: Implications for the evolution of group living in canids? *Behavioral Ecology* **15**, 766–778.

Bergerud, A.T. & Miller, D.R. (1977) Population dynamics of Newfoundland beaver. *Canadian Journal of Zoology* **55**, 1480–1492.

Bezerra, B.M., Souto, A.D.S. & Schiel, N. (2007) Infanticide and Cannibalism in a Free-Ranging Plurally Breeding Group of Common Marmosets (*Callithrix Jacchus*). *American Journal of Primatology* **69**, 945–952.

Bingham, J. & Purchase, G. k. (2002) Reproduction in the jackals *Canis adustus* Sundevall, 1846, and *Canis mesomelas* Schreber, 1778 (Carnivora: Canidae), in Zimbabwe. *African Zoology* **37**, 21–26.

Bingham, J. & Purchase, G.K. (2003) Age determination in jackals (*Canis adustus* Sundevall, 1846, and *Canis mesomelas* Schreber, 1778; Carnivora: Canidae) with reference to the age structure and breeding patterns of jackal populations in Zimbabwe. *African Zoology* **38**, 153–160.

Bishop, J.M., Jarvis, J.U.M., Spinks, A.C., Bennett, N.C. & O’Ryan, C. (2004) Molecular insight into patterns of colony composition and paternity in the common mole-rat *Cryptomys hottentotus hottentotus*. *Molecular Ecology* **13**, 1217–1229.

Braude, S. (1991) The behavior and demographics of the naked mole-rat, *Heterocephalus glaber*. The University of Michigan.

Buffenstein, R., Amoroso, V., Andziak, B., Avdieiev, S., Azpurua, J., Barker, A.J., Bennett, N.C., Brieño-Enríquez, M.A., Bronner, G.N., Coen, C., Delaney, M.A., Dengler-Crish, C.M., Edrey, Y.H., Faulkes, C.G., Frankel, D., et al. (2022) The naked truth: a comprehensive clarification and classification of current ‘myths’ in naked mole-rat biology. *Biological Reviews* **97**, 115–140.

Burland, T.M., Bennett, N.C., Jarvis, J.U.M. & Faulkes, C.G. (2004) Colony structure and parentage in wild colonies of co-operatively breeding Damaraland mole-rats suggest incest avoidance alone may not maintain reproductive skew. *Molecular Ecology* **13**, 2371–2379.

Burrows, R. (1995) Demographic changes and social consequences in wild dogs, 1964–1992. In *Serengeti II: dynamics, management and conservation of an ecosystem* (eds A.R.E. Sinclair, C. Packer, S.A.R. Mduma & J.M. Fryxell), pp. 400–420. University of Chicago Press, Chicago, Illinois.

Busher, P.E., Warner, R.J. & Jenkins, S. (1983) Population Density, Colony Composition, and Local Movements in Two Sierra Nevadan Beaver Populations. *Journal of Mammalogy* **64**, 314–318.

Calegaro-Marques, C., Bicca-Marques, J. & Azevedo, M. (1995) Two breeding females in a Saguinus fuscicollis weddelli group. *Neotropical Primates* **3**, 183.

Cameron, C., Berteaux, D. & Dufresne, F. (2011) Spatial variation in food availability predicts extrapair paternity in the arctic fox. *Behavioral Ecology* **22**, 1364–1373.

Campbell, R.D., Rosell, F., Nolet, B.A. & Dijkstra, V.A.A. (2005) Territory and group sizes in Eurasian beavers (*Castor fiber*): echoes of settlement and reproduction? *Behavioral Ecology and Sociobiology* **58**, 597–607.

Carmichael, L.E., Szor, G., Berteaux, D., Giroux, M.A., Cameron, C. & Strobeck, C. (2007) Free love in the far north: Plural breeding and polyandry of arctic foxes (*Alopex lagopus*) on Bylot Island, Nunavut. *Canadian Journal of Zoology* **85**, 338–343.

Carpenter, P.J., Pope, L.C., Greig, C., Dawson, D.A., Rogers, L.M., Erven, K., Wilson, G.J., Delahay, R.J., Cheeseman, C.L. & Burke, T. (2005) Mating system of the Eurasian badger, *Meles meles*, in a high density population. *Molecular Ecology* **14**, 273–284.

Clutton-Brock, T.H., Brotherton, P.N.M., Russell, A.F., O’Riain, M.J., Gaynor, D., Kansky, R., Griffin, A., Manser, M., Sharpe, L., Mcilrath, G.M., Small, T., Moss, A. & Monfort, S. (2001) Cooperation, Control, and Concession in Meerkat Groups. *Science* **291**, 478–481.

Clutton-Brock, T.H., Hodge, S.J. & Flower, T.P. (2008) Group size and the suppression of subordinate reproduction in Kalahari meerkats. *Animal Behaviour* **76**, 689–700.

Clutton-Brock, T.H., Maccoll, A., Chadwick, P., Gaynor, D., Kansky, R. & Skinner, J.D. (1999) Reproduction and survival of suricates (*Suricata suricatta*) in the southern Kalahari. *African Journal of Ecology* **37**, 69–80.

Cohen, J.A. (1978) *Cuon alpinus*. *Mammalian Species* **100**, 1–3.

Converse, K.E. (2012) Genetic mating system and territory inheritance in the Sacramento Valley red fox. California State University, Sacramento.

Crawford, J.C. (2007) Mating, kinship, and population structure in Illinois beaver populations. Eastren Illinois University.

Crawford, J.C., Liu, Z., Nelson, T.A., Nielsen, C.K. & Bloomquist, C.K. (2008) Microsatellite analysis of mating and kinship in beavers (*Castor canadensis*). *Journal of Mammalogy* **89**, 575–581.

Creel, S., Creel, N.M., Mills, M.G.L. & Monfort, S.L. (1997) Rank and reproduction in cooperatively breeding African wild dogs: Behavioral and endocrine correlates. *Behavioral Ecology* **8**, 298–306.

Culot, L., Lledo-Ferrer, Y., Hoelscher, O., Mun˜oz Lazo, F.J.J., Huynen, M.-C. & Heymann, E.W. (2011) Reproductive failure, possible maternal infanticide, and cannibalism in wild moustached tamarins, *Saguinus mystax*. *Primates* **52**, 179–186.

Davidar, E.R.C. (1974) Observations at the dens of the Dhole or Indian wild dog (*Cuon alpinus*). *The journal of the Bombay Natural History Society* **71**, 183–187.

Decanini, D.P. & Macedo, R.H. (2008) Sociality in *Callithrix penicillata*: II. Individual strategies during intergroup encounters. *International Journal of Primatology* **29**, 627–639.

Dietz, J.M. & Baker, A.J. (1993) Polygyny and female reproductive success in golden lion tamarins, *Leontopithecus rosalia*. *Animal Behaviour* **46**, 1067–1078.

Digby, L.J. & Barreto, C.E. (1993) Social organization in a wild population of *Callithrix jacchus*. I. Group composition and dynamics. *Folia Primatologica* **61**, 123–134.

Digby, L.J. & Ferrari, S.F. (1994) Multiple Breeding Females in Free-Ranging Groups of *Callithrix jacchus*. *International Journal of Primatology* **15**, 389–397.

Doolan, S.P. & MacDonald, D.W. (1997) Band Structure and Failures of Reproductive Suppression in a Cooperatively Breeding Carnivore, the Slender-Tailed Meerkat (*Suricata suricatta*). *Behaviour* **134**, 827–848.

Dugdale, H.L. (2007) The evolution of social behaviour: the effect of mating system and social structure in the European badger *Meles meles*. Linacre College, University of Oxford.

Dugdale, H.L., Ellwood, S.A. & Macdonald, D.W. (2010) Alloparental behaviour and long-term costs of mothers tolerating other members of the group in a plurally breeding mammal. *Animal Behaviour* **80**, 721–735.

Dugdale, H.L., Macdonald, D.W., Pope, L.C. & Burke, T. (2007) Polygynandry, extra-group paternity and multiple-paternity litters in European badger (Meles meles) social groups. *Molecular Ecology* **16**, 5294–5306.

Dugdale, H.L., Macdonald, D.W., Pope, L.C., Johnson, P.J. & Burke, T. (2008) Reproductive skew and relatedness in social groups of European badgers, *Meles meles*. *Molecular Ecology* **17**, 1815–1827.

Erb, W.M. & Porter, L.M. (2020) Variable infant care contributions in cooperatively breeding groups of wild saddleback tamarins. *American Journal of Primatology* **82**, 1–14.

Faulkes, C.G. & Bennett, N.C. (2009) Reproductive skew in African mole-rats: behavioral and physiological mechanisms to maintain high skew. In *Reproductive Skew in Vertebrates: Proximate and Ultimate Causes* (eds R. Hager & C.B. Jones), pp. 369–396. Cambridge University Press, Cambridge, UK.

Federico, V., Allainé, D., Gaillard, J. & Cohas, A. (2020) Evolutionary Pathways to Communal and Cooperative Breeding in Carnivores. *The American Naturalist* **195**, 1037–1055.

Ferguson, J.W.H., Nel, J.A.J. & Wet, M.J. (1983) Social organization and movement patterns of Black-backed jackals *Canis mesomelas* in South Africa. *Journal of Zoology* **199**, 487–502.

Fischer, J.W., Joos, R.E., Neubaum, M.A., Taylor, J.D., Bergman, D.L., Nolte, D.L. & Piaggio, A.J. (2010) Lactating North American beavers (*Castor canadensis*) sharing dens in the southwestern United States. *The Southwestern Naturalist* **55**, 273–277.

FitzGerald, R.W. & Madison, D.M. (1983) Social organization of a free-ranging population of pine voles, *Microtus pinetorum*. *Behavioral Ecology and Sociobiology* **13**, 183–187.

Foltz, D.W. (1981) Genetic Evidence for Long-Term Monogamy in a Small Rodent, *Peromyscus polionotus*. *The American naturalist* **117**, 665–675.

French, J.A., Bales, K.L., Baker, A.J. & Dietz, J.M. (2003) Endocrine Monitoring of Wild Dominant and Subordinate Female *Leontopithecus rosalia*. *International Journal of Primatology* **24**, 1281–1300.

Getz, L.L. & McGuire, B. (1997) Communal nesting in prairie voles (*Microtus ochrogaster*): Formation, composition, and persistence of communal groups. *Canadian Journal of Zoology* **75**, 525–534.

Goldizen, A., Mendelson, J., van Vlaardingen, M. & Terborgh, J. (1996) Saddle-Back Tamarin (*Saguinus fuscicollis*) Reproductive Strategies: Evidence From a Thirteen-Year Study of a Marked Population. *American Journal of Primatology* **38**, 57–83.

Goltsman, M., Kruchenkova, E.P., Sergeev, S., Volodin, I. & Macdonald, D.W. (2005) ‘Island syndrome’ in a population of Arctic foxes (*Alopex lagopus*) from Mednyi Island. *Journal of Zoology* **267**, 405–418.

Griffin, A.S., Pemberton, J.M., Brotherton, P.N.M., Mcilrath, G., Gaynor, D., Kansky, R., Riain, J.O. & Clutton-Brock, T.H. (2003) A genetic analysis of breeding success in the cooperative meerkat (*Suricata suricatta*). *Behavioral Ecology* **14**, 472–480.

Gromov, V.S. (2021) Ecology and social behaviour of the Mongolian gerbil: A generalised review. *Behaviour* **1**, 1–39.

Haber, G.C. (1977) Socio-ecological dynamics of wolves and prey in a subarctic ecosystem. University of British Columbia.

Hackländer, K., Möstl, E. & Arnold, W. (2003) Reproductive suppression in female Alpine marmots, Marmota marmota. *Animal Behaviour* **65**, 1133–1140.

Harrington, F., Paquet, P., Ryon, J. & Fentress, J. (1982) Monogamy in wolves: a review of the evidence. In *Wolves of the world: perspectives of behavior, ecology, and conservation* (eds F.H. Harrington & P.C. Paquet), pp. 209–222. Noyes Publications, Park Ridge, New Jersey.

Hatier, G.K. (1995) Effects of helping behaviors on coyote packs in Yellowstone National Park, Wyoming. Montana State University-Bozeman.

Hennessy, A.C., Dubach, J. & Gehrt, S.D. (2012) Long-term pair bonding and genetic evidence for monogamy among urban coyotes (*Canis latrans*). *Journal of Mammalogy* **93**, 732–742.

Hennessy, C.A. (2007) Mating Strategies And Pack Structure of Coyotes in an Uban Landscape: A Genetic Investigation. Ohio State University.

Henry, M.D. (2011) Proximate mechanisms and ultimate causes of female reproductive skew in cooperatively breeding golden lion tamarins, *Leontopithecus rosalia*. University of Maryland, College Park.

Henry, M.L.D., Hankerson, S.J., Siani, J.M., French, J.A. & Dietz, J.M. (2013) High rates of pregnancy loss by subordinates leads to high reproductive skew in wild golden lion tamarins (*Leontopithecus rosalia*). *Hormones and Behavior* **63**, 675–683. Elsevier Inc.

Heslin Piper, L.A. (2015) Social and Environmental Factors Influencing Reproductive Success in a Cooperatively Breeding Primate by Success in a Cooperatively Breeding Primate. University of Toront.

Heslin Piper, L.A., Dietz, J.M. & Raboy, B.E. (2017) Multi-male groups positively linked to infant survival and growth in a cooperatively breeding primate. *Behavioral Ecology and Sociobiology* **71**, 1–12. Behavioral Ecology and Sociobiology.

Huck, M., Löttker, P., Böhle, U. & Heymann, E.W. (2005) Paternity and Kinship Patterns in Polyandrous Moustached Tamarins (*Saguinus mystax*). *American Journal of Physical Anthropology* **127**, 449–464.

Jarvis, J.U.M. & Bennett, N.C. (1993) Eusociality has evolved independently in two genera of bathyergid mole-rats - but occurs in no other subterranean mammal. *Behavioral Ecology and Sociobiology* **33**, 253–260.

Johnsingh, A.J.T. (1982) Reproduction and social behaviour of the dhole, *Cuon alpinus* (Canidae). *Journal of Zoology* **198**, 443–463.

Keane, B., Waser, P.M., Creel, S.R., Creel, N.M., Elliott, L.F. & Minchella, D.J. (1994) Subordinate reproduction in dwarf mongooses. *Animal Behaviour* **47**, 65–75.

Kingdon, J. (1988) *East African mammals: an atlas of evolution in Africa, volume 3, Part A: Carnivores*. University of Chicago Press.

Kruchenkova, E.P. & Formozov, N. (1995) The Arctic foxes of Mednyi (Copper) Island. *Russian Conservation News* **2**, 19–20.

Lacey, E. & Sherman, P.W. (1997) Cooperative Breeding in Naked Mole-Rats: Implications for Vertebrate and Invertebrate Sociality. In *Cooperative breeding in mammals* (eds N. Solomon & J. French), pp. 267–301. Cambridge University Press, New York.

Lazaro-Perea, C., Castro, C.S.S., Harrison, R., Araujo, A., Arruda, M.F. & Snowdon, C.T. (2000) Behavioral and demographic changes following the loss of the breeding female in cooperatively breeding marmosets. *Behavioral Ecology and Sociobiology* **48**, 137–146.

Liu, W., Wang, G., Wang, T., Zhong, W. & Wan, X. (2009) Population ecology of wild Mongolian gerbils Meriones unguiculatus. *Journal of Mammalogy* **90**, 832–840.

Löttker, P., Huck, M. & Heymann, E.W. (2004) Demographic Parameters and Events in Wild Moustached Tamarins (*Saguinus mystax*). *American Journal of Primatology* **64**, 425–449.

Loveridge, A.J. & Macdonald, D.W. (2001) Seasonality in spatial organization and dispersal of sympatric jackals (*Canis mesomelas* and *C. adustus*): Implications for rabies management. *Journal of Zoology* **253**, 101–111.

Lukas, D. & Clutton-Brock, T. (2012) Cooperative breeding and monogamy in mammalian societies. *Proceedings of the Royal Society B: Biological Sciences* **279**, 2151–2156.

Malcolm, J.R. & Marten, K. (1982) Natural selection and the communal rearing of pups in African wild dogs (*Lycaon pictus*). *Behavioral Ecology and Sociobiology* **10**, 1–13.

Margulis, S.W., Nabong, M., Alaks, G., Walsh, A. & Lacy, R.C. (2005) Effects of early experience on subsequent parental behaviour and reproductive success in oldfield mice, *Peromyscus polionotus*. *Animal Behaviour* **69**, 627–634.

McGuire, B., Getz, L.L. & Oli, M.K. (2002) Fitness consequences of sociality in prairie voles, *Microtus ochrogaster*: Influence of group size and composition. *Animal Behaviour* **64**, 645–654.

Mech, L.D. (1999) Alpha status, dominance, and division of labor in wolf packs. *Canadian Journal of Zoology* **77**, 1196–1203.

Mech, L.D. & Nelson, M.E. (1989) Polygyny in a wild wolf pack. *Journal of Mammalogy* **70**, 675–676.

Mech, L.D., Wolf, P.C. & Packard, J.M. (1999) Regurgitative food transfer among wild wolves. *Canadian Journal of Zoology* **77**, 1192–1195.

Moehlman, P.D. (1979) Jackal helpers and pup survival. *Nature* **277**, 382–383.

Moehlman, P.D. (1983) Socioecology of silverbacked and golden jackals (Canis mesomelas and Canis aureus). In *Advances in the study of mammalian behavior 7* (eds J.F. Eisenberg & D.G. Kleiman), pp. 423–453. The American Society of Mammalogists, Kansas.

Muller-Schwarzei, D. & Schulte, B.A. (1999) Behavioral and ecological characteristics of a ‘climax’ population of beaver (Castor canadensis). In *Beaver Protection, Management, and Utilization in Europe and North America* (eds P.E. Busher & R.M. Dziciaowski), pp. 161–178. Kluwer Academic/Plenum Publishers.

Murie, A. (1944) *The Wolves of Mount McKinley No. 5*. U.S. National Park Service, Washington, DC.

Nattrass, N., Conradie, B., Drouilly, M. & O’Riain, M.J. (2017) Understanding the black-backed jackal. *CSSR Working Paper No. 399*.

Newman, C., Zhou, Y.-B., Buesching, C.D., Kaneko, Y. & Macdonald, D.W. (2011) Contrasting Sociality in Two Widespread, Generalist, Mustelid Genera, *Meles* and *Martes*. *Mammal Study* **36**, 169–188.

Nievergelt, C.M., Digby, L.J., Ramakrishnan, U. & Woodruff, D.S. (2000) Genetic analysis of group composition and breeding system in a wild common marmoset (*Callithrix jacchus*) population. *International Journal of Primatology* **21**, 1–20.

Norén, K., Dalén, L., Flagstad, Ø., Berteaux, D., Wallén, J. & Angerbjörn, A. (2017) Evolution, ecology and conservation—revisiting three decades of Arctic fox population genetic research. *Polar Research* **36**, 1–14. Routledge.

Norén, K., Hersteinsson, P., Samelius, G., Eide, N.E., Fuglei, E., Elmhagen, B., Dalén, L., Meijer, T. & Angerbjörn, A. (2012) From monogamy to complexity: Social organization of arctic foxes (*Vulpes lagopus*) in contrasting ecosystems. *Canadian Journal of Zoology* **90**, 1102–1116.

O’Riain, M.J., Bennett, N.C., Brotherton, P.N.M., McIlrath, G. & Clutton-Brock, T.H. (2000) Reproductive suppression and inbreeding avoidance in wild populations of co-operatively breeding meerkats (*Suricata suricatta*). *Behavioral Ecology and Sociobiology* **48**, 471–477.

Passos, F.C. (1994) Behavior of the black lion tamarin, Leontopithecus chrysopygus, in different forest levels in the Caetetus Ecological Station, São Paulo, Brazil. *Neotropical Primates* **2 (suppl.)**, 40–41.

Pecorella, S., De Luca, M., Fonda, F., Viviano, A., Candelotto, M., Candotto, S., Mori, E. & Banea, O. (2023) First record of allonursing in golden jackal (*Canis aureus*, L. 1758): a case of double breeding and communal denning within the same social unit. *European Journal of Wildlife Research* **69**, 43.

Peterson, R.O., Jacobs, A.K., Drummer, T.D., Mech, L.D. & Smith, D.W. (2002) Leadership behavior in relation to dominance and reproductive status in gray wolves, *Canis lupus*. *Canadian Journal of Zoology* **80**, 1405–1412.

Raboy, B.E. (2002) The ecology and behavior of wild golden-headed lion tamarins (Leontopithecus chrysomelas). PhD, University of Maryland, College Park.

Raihani, N.J. & Clutton-Brock, T.H. (2010) Higher reproductive skew among birds than mammals in cooperatively breeding species. *Biology letters* **6**, 630–632.

Ramirez, M. (1984) Population Recovery in the Moustached Tamarin (Saguinus mystax): Management Strategies and Mechanisms of Recovery. *American Journal of Primatology* **7**, 245–259.

Ramirez, M.M. (1989) Feeding ecology and demography of the moustached tamarin *Saguinus mystax* in northeastern Peru. City University of New York.

Randall, D.A., Pollinger, J.P., Wayne, R.K., Tallents, L.A., Johnson, P.J. & Macdonald, D.W. (2007) Inbreeding is reduced by female-biased dispersal and mating behavior in Ethiopian wolves. *Behavioral Ecology* **18**, 579–589.

Ribble, D.O. (1992) Dispersal in a Monogamous Rodent, *Peromyscus Californicus*. *Ecology* **73**, 859–866.

Ribble, D.O. (2003) The evolution of social and reproductive monogamy in Peromyscus: Evidence from Peromyscus californicus (the California mouse). In *Monogamy: Mating Strategies and Partnerships in Birds, Humans and Other Mammals* (eds U. Reichard & C. Boesch), pp. 81–92. Cambridge University Press.

Rood, J.P. (1980) Mating relationships and breeding suppression in the dwarf mongoose. *Animal Behaviour* **85**, 143–150.

Savage, A., Giraldo, L.H., Soto, L.H. & Snowdon, C.T. (1996) Demography, Group Composition, and Dispersal in Wild Cotton-Top Tamarin (*Saguinus oedipus*) Groups. *American Journal of Primatology* **38**, 85–100.

Savage, A., Snowdon, C.T., Soto, L., Medina, F., Emeris, G. & Guillen, R. (2021) Factors influencing the survival of wild cotton-top tamarin (Saguinus oedipus) infants. *American Journal of Primatology* **83**, 1–10.

Scanlon, C.E., Chalmers, N.R. & Monteiro da Cruz, M.A.O. (1988) Changes in the size, composition, and reproductive condition of wild marmoset groups (*Callithrix jacchus jacchus*) in north east Brazil. *Primates* **29**, 295–305.

Scharff, A., Locker-Grütjen, O., Kawalika, M. & Burda, H. (2001) Natural history of the giant mole-rat, Cryptomys mechowi (rodentia: Bathyergidae), from Zambia. *Journal of Mammalogy* **82**, 1003–1015.

Schradin, C. & Pillay, N. (2004) The Striped Mouse (*Rhabdomys pumilio*) From the Succulent Karoo, South Africa: A Territorial Group-Living Solitary Forager With Communal Breeding and Helpers at the Nest. *Journal of Comparative Psychology* **118**, 37–47.

Schradin, C., Pillay, N. & Bertelsmeier, C. (2019) Social flexibility and environmental unpredictability in African striped mice. *Behavioral Ecology and Sociobiology* **73**, 1–12. Behavioral Ecology and Sociobiology.

Schradin, C., Schneider, C. & Lindholm, A.K. (2010) The nasty neighbour in the striped mouse (*Rhabdomys pumilio*) steals paternity and elicits aggression. *Frontiers in Zoology* **7**, 1–8.

Schubert, M., Pillay, N. & Schradin, C. (2009) Parental and alloparental care in a polygynous mammal. *Journal of Mammalogy* **90**, 724–731.

Shavadze, I. (2016) Extra pair copulation in the Eurasian beaver (Castor fiber)? University College of Southeast Norway.

Sichilima, A.M., Faulkes, C.G. & Bennett, N.C. (2008) Field evidence for aseasonality of reproduction and colony size in the Afrotropical giant mole-rat *Fukomys mechowii* (Rodentia: Bathyergidae). *African Zoology* **43**, 144–149.

Sillero-Zubiri, C., Gottelli, D. & Macdonald, D.W. (1996) Male philopatry, extra-pack copulations and inbreeding avoidance in Ethiopian wolves (*Canis simensis*). *Behavioral Ecology and Sociobiology* **38**, 331–340.

Smith, A.C. & Tirado, Herrera E. R. Buchanan-Smith, H. M. Heymann, E.W. (2001) Multiple breeding females and allo-nursing in a wild group of moustached tamarins (*Saguinus mystax*). *Neotropical Primates* **9**, 67–69.

Soini, P. (1988) The Pygmy Marmoset, Genus Cebuella. In *The Ecology and Behavior of Neotropical Primates, Vol. II* (eds R.A. Mittermeier, A. Rylands, A.F. Coimbra-Filho & G.A.B. Fonseca), pp. 79–129. World Wildlife Fund, Washington, DC.

Solomon, N.G. & Keane, B. (2018) Dispatches from the field: sociality and reproductive success in prairie voles. *Animal Behaviour* **143**, 193–203.

Solomon, N.G., Vandenbergh, J.G. & Sullivan, W.T. (1998) Social influences on intergroup transfer by pine voles (*Microtus pinetorum*). *Canadian Journal of Zoology* **76**, 2131–2136.

Sparkman, A.M., Adams, J.R., Steury, T.D., Waits, L.P. & Murray, D.L. (2011) Direct fitness benefits of delayed dispersal in the cooperatively breeding red wolf (*Canis rufus*). *Behavioral Ecology* **22**, 199–205.

Sparkman, A.M., Adams, J.R., Steury, T.D., Waits, L.P. & Murray, D.L. (2012) Pack social dynamics and inbreeding avoidance in the cooperatively breeding red wolf. *Behavioral Ecology* **23**, 1186–1194.

Spiering, P.A., Somers, M.J., Maldonado, J.E., Wildt, D.E. & Gunther, M.S. (2010) Reproductive sharing and proximate factors mediating cooperative breeding in the African wild dog (*Lycaon pictus*). *Behavioral Ecology and Sociobiology* **64**, 583–592.

Spong, G.F., Hodge, S.J., Young, A.J. & Clutton-Brock, T.H. (2008) Factors affecting the reproductive success of dominant male meerkats. *Molecular Ecology* **17**, 2287–2299.

Strand, O., Landa, A., Linnell, J.D.C., Zimmermann, B. & Skogland, T. (2000) Social Organization and Parental Behavior in the Arctic Fox. *Journal of Mammalogy* **81**, 223–233.

Sun, L. (2003) Monogamy correlates, socioecological factors, and mating systems in beavers. In *Monogamy: Mating Strategies and Partnerships in Birds, Humans and Other Mammals* (eds R. Ulrich & C. Boesch), pp. 138–146. Cambridge University Press.

Svendsen, G.E. (1989) Pair formation, duration of pair-bonds, and mate replacement in a population of beavers (*Castor canadensis*). *Canadian Journal of Zoology* **67**, 336–340.

Syrůčková, A., Saveljev, A.P., Frosch, C., Durka, W. & Savelyev, A. A. Munclinger, P. (2015) Genetic relationships within colonies suggest genetic monogamy in the Eurasian beaver (*Castor fiber*). *Mammal Research* **60**, 139–147.

Tannerfeldt, M., Moehrenschlager, A. & Angerbjörn, A. (2003) Den ecology of swift, kit and arctic foxes: a review. In *Ecology and conservation of swift foxes in a changing world* (eds M. Sovada & L. Carbyn), pp. 167–181. Canadian Plains Research Center, Regina, Sask.

Valladares-Padua, C. (1993) The ecology, behavior and conservation of the black lion tamarins (*Leontopithecus chrysopygus* Mikan, 1823). University of Florida.

Venkataraman, A.B. (1998) Male-biased adult sex ratios and their significance for cooperative breeding in dhole, *Cuon alpinus*, packs. *Ethology* **104**, 671–684.

de Vleeschouwer, K., van Elsacker, L. & Leusa, K. (2001) Multiple Breeding Females in Captive Groups of Golden-Headed Lion Tamarins (*Leontopithecus chrysomelas*): Causes and Consequences. *Folia Primatologica* **72**, 1–10.

Wang, Y., Liu, W., Wang, G.M., Zhong, W. & Wan, X. (2011) Genetic consequences of group living in Mongolian gerbils. *Journal of Heredity* **102**, 554–561.

Wheatley, M. (1993) Report of two pregnant beavers, Castor canadensis, at one beaver lodge. *Canadian Field-Naturalist* **107**, 103.

Wheaton, C.J., Feilen, K.L., Soto, L.H., Medina, F., Emeris, G., Guillen, R. & Savage, A. (2022) Seasonality of reproduction in wild cotton-top tamarins (Saguinus oedipus) in Colombia. *American Journal of Primatology* **84**, 1–14.

White, P.A. (1992) Social organization and activity patterns of the arctic fox (*Alopex lagopus pribilofensis*) on St. Paul Island, Alaska. University of California at Berkeley.

Woodroffe, R. (1993) Alloparental behaviour in the European badger. *Animal Behaviour* **46**, 413–415.

Woodroffe, R. & Macdonald, D.W. (2000) Helpers provide no detectable benefits in the European badger (*Meles meles*). *Journal of Zoology* **250**, 113–119.

Yeboah, S. & Dakwa, K.B. (2002a) Colony and social structure of the ghana mole-rat (Cryptomys zechi, Matchie) (Rodentia: Bathyergidae). *Journal of Zoology* **256**, 85–91.

Yeboah, S. & Dakwa, K.B. (2002b) Aspects of the feeding habits and reproductive biology of the Ghana mole-rat *Cryptomys zechi* (Rodentia, Bathyergidae). *African Journal of Ecology* **40**, 110–119.

Young, A.J. & Bennett, N.C. (2010) Morphological divergence of breeders and helpers in wild damaraland mole-rat societies. *Evolution* **64**, 3190–3197.

Zabel, C.J. & Taggart, S.J. (1989) Shift in red fox, *Vulpes vulpes*, mating system associated with El Niño in the Bering Sea. *Animal Behaviour* **38**, 830–838.
