## Supplementary material for "Evidence for a reproductive sharing continuum in cooperatively breeding mammals and birds: consequences for comparative research": Table S2

| **Table S2.** Within-group distribution of maternity and paternity in bird species that were classified by (i) Raihani & Clutton-Brock (2010) as exhibiting alloparental care and extreme female reproductive skew and/or (ii) species classified by Cornwallis *et al.* (2010) as engaging in alloparental care that is provided by “helpers [that do] not breed or [have] zero-to-limited opportunities to breed” (i.e., extreme reproductive skew in males and/or females). Results are from studies of wild populations, after excluding extra-group parentage (when possible), considering only multi-female or multi-male groups (when possible) and considering only clutches with >1 sampled offspring when possible (i.e., where shared maternity/paternity can be detected). | | | | | | | | |
| --- | --- | --- | --- | --- | --- | --- | --- | --- |
| **Species** | | **Sample:**  Only social groups with multiple sexually mature females/males  or  A mixture of single-female/male and multi-female/male groups | **Percentage of offspring produced by the most dominant female/male in the social group**  +  (Type of evidence:  Genetic/ Behavioural/ Physiological) | **Percentage of social groups with a single breeding female/male**  +  (Type of evidence:  Genetic/ Behavioural/ Physiological) | **Reproductive sharing:**  Extreme reproductive skew  /  Not extreme reproductive skew | **Combined sample size:**  Anecdotal (≤5 groups or ≤10 group-years);  Limited (6–15 groups or 11–30 group-years);  Substantial (≥16 groups or ≥31 group-years) | **Remarks** | **Data sets that classified the species as exhibiting alloparental care and extreme reproductive skew:**  Raihani & Clutton-Brock (2010) (for females only)  Cornwallis *et al.* (2010) (for males and/or females) |
| **Merlin**  (*Falco columbarius*) | Females | **Not relevant** ^1^ | | | | | ^1^ We were unable to find systematic evidence for multi-female groups that engage in alloparental care (James & Oliphant, 1986, p. 533; Sodhi, 1989, pp. 506–507, 1991, p. 436; Warkentin *et al.*, 1994, p. 233, 2013, p. 70). | - Cornwallis *et al.* (2010) |
|  | Males | **Not relevant** ^1^ | | | | | ^1^ We were unable to find systematic evidence for multi-male groups that engage in alloparental care (James & Oliphant, 1986, p. 533; Sodhi, 1989, pp. 506–507, 1991, p. 436; Warkentin *et al.*, 1994, p. 233, 2013, p. 70). |  |
| **Common eider**  (*Somateria mollissima*) | Females | **Not relevant** ^1-2^ | | | | | ^1^ There are no cohesive social groups in this species. Each female has her own nest where she lays eggs. About 6% of eggs are brood parasitism by other females and 20%-22% of nests are parasitized, but this is done by females that may have their own nest and may not provide care later (Waldeck *et al.*, 2004). During the first week of ducklings’ life (Öst, 1999, p. 599) about 56% - 71% of broods are joined into temporal associations (Öst, 1999, p. 602). Hence, there is no monopolisation of breeding in this species, but rather an amalgamation of broods.  ^2^ The alleged care may not be qualified as alloparental care as it only includes vigilance by associated females (Öst *et al.*, 2007, p. 74). Vigilance may be conducted for self-defence against predators or to guard one’s offspring. Anti-predator defence for other’s offspring which are nearby is a by-product of sociality, rather than alloparental care (Ben Mocha, Scemama de Gialluly & Markman, n.d.). | - Raihani & Clutton-Brock (2010) |
|  | Males | **Not relevant** ^1-2^ | | | | | ^1^ There are no social groups in this species and brood coalitions (i.e., ‘‘creche’’) involve mothers only (Öst *et al.*, 2003, pp. 311–312).  ^2^ In addition, common eiders were classified as a cooperative breeding species only by Raihani & Clutton-Brock (2010) who did not consider reproductive skew in males. |  |
| **Western bluebird**  (*Sialia Mexicana*) | Females | **Not relevant** ^1^ | | | | | ^1^ No adult female helpers in the population of this species in Hastings Reservation, California USA (Dickinson, Koenig & Pitelka, 1996, p. 169; Dickinson & Akre, 1998, p. 96).  In an experimental population in northwestern Oregon, USA, 1.3% of helpers were females, but it is not clear whether these female helpers were sexually mature (Charmantier, Keyser & Promislow, 2007, p. 1758) | - Raihani & Clutton-Brock (2010) - Cornwallis *et al.* (2010) |
|  | Males | Only groups with multiple sexually mature males | <89% ^1^  )N = 27 nestlings, 6 nests of two brothers) ^2-3^  (Dickinson & Akre, 1998, p. 101) ^Genetic evidence^ | <83% ^1^  )N = 6 nests of two brothers) ^2-3^  (Dickinson & Akre, 1998, p. 101) ^Genetic evidence^ | **Not extreme reproductive skew** | **Limited** | ^1^ The study did not assess within-group dominance. Thus, the observed offspring could belong to any of these categories of dominance.  ^2^ Since both males were brothers, there was no incest limitation on either of the males. This is probably a sample of groups. But it could also be that the same pair of brothers produced more than one nest (i.e., group-years).  ^3^ This sample does not exclude paternity by extra-group males, who may have sired some of the offspring in this sample (Dickinson & Akre, 1998, p. 101) ^Genetic evidence^. Hence, this is a maximum estimation of reproductive output by dominant (as it assumed all offspring not sired by helpers are of the group’s breeder male). |  |
| **Superb fairy-wren**  (*Malurus cyaneus*) | Females |  |  |  | **Not relevant** ^1^ |  | ^1^ No female helpers in this species  (Rowley, 1965, pp. 277–278, 288; Dunn, Cockburn & Mulder, 1995, p. 340; Dunn & Cockburn, 1999, p. 393). | - Raihani & Clutton-Brock (2010) - Cornwallis *et al.* (2010) |
|  | Males | Only groups with multiple sexually mature males | 68%  (N = 146 broods) ^1^  (Hajduk *et al.*, 2021, p. 390) ^Genetic evidence^  81%  (N = 80 offspring, 51 group-years) ^2^  (Dunn *et al.*, 1995, p. 342)  ^Genetic evidence^  50%  (N = 2 offspring, 1 group) ^3^  (Colombelli-Négrel, Schlotfeldt & Kleindorfer, 2009, pp. 300–302) ^Genetic evidence^ | 0%  (N = 1 group) ^3^  (Colombelli-Négrel *et al.*, 2009, pp. 300–302) ^Genetic evidence^ | **Not extreme reproductive skew** | **Substantial** | ^1^ The number of offspring in this sample is not indicated. The percentage is based on the last row in Table 2 after excluding extra-group paternity. This sample consists of multi-male groups with at least one unrelated male helper to the female. It is assumed to be a group-years sample.  **Population:** Australian National Botanic Gardens, Canberra, Australia.  ^2^ This is a sample of groups with male helpers. But it does not exclude extra-group paternity and not groups in which helpers were related to the breeding female.  Population: Australian National Botanic Gardens, Canberra, Australia.  ^3^ One offspring was excluded as extra-group paternity. **Population:** Newland Head Conservation Park, Fleurieu Peninsula, Australia |  |
| **Noisy miner** (*Manorina melanocephala*) | Females | Only groups with multiple sexually mature females |  | 100%  (N ≤ 7 broods) ^1^  (Barati *et al.*, 2018a, p. 1382) ^Genetic evidence^ | **Extreme reproductive skew** ^2^ | **Limited** | ^1^ There were only 7 female helpers in this study. So, if each helper attended one nest, 7 is the maximum number of nests with female helpers. Though, it may be that the same groups with helpers produced more than 1 brood, hence the number of group-years may be higher. All female helpers were unrelated to the breeding pair (Barati *et al.*, 2018a Figure 1). Population: Two colonies at Armidale, NSW, Australia.  ^2^ Social groups of noisy miners have very few female helpers.  No female helpers in the population of Lake Wivenhoe, Australia studied by these authors (Dow, 1979, pp. 71, 73; Põldmaa, Montgomerie & Boag, 1995, pp. 138, 140).  In two colonies at Armidale, NSW, Australia, only 7% (N = 103) of helpers were females (Barati *et al.*, 2018a, p. 1385). | - Raihani & Clutton-Brock (2010) - Cornwallis *et al.* (2010) |
|  | Males | A mixture of single-male and multi-male groups | 98%  (N = 55 nestlings, 25 broods) ^1^  (Barati *et al.*, 2018b, pp. 246–248) ^Genetic evidence^ | 100% ^2^  (N = 30 brood-years) ^3^  (Põldmaa *et al.*, 1995, pp. 140–141) ^Genetic evidence^  96%  (N = 25 broods) ^4^  (Barati *et al.*, 2018b, pp. 247–248) ^Genetic evidence^ | **Extreme reproductive skew** ^5^ | **Substantial** | ^1^ Only in one brood did two fathers feed the nestlings (Barati *et al.*, 2018b, pp. 246–248). Two nestlings were excluded as extra-group paternity (Barati *et al.*, 2018b, pp. 246–248). It is assumed to be a group-year sample. Population: Two colonies at Armidale, NSW, Australia.  ^2^ Only males that fed the nestling were considered members of the group. But as dominance was not assessed in this study, it is not clear whether the most dominant male always sired all offspring. Hence, also the maternity proxy was not calculated from this study’s data. Population: Lake Wivenhoe, Australia.  ^3^ We considered only broods with more than one sampled nestling. In addition, one brood with multiple nestlings (nest “24”) was excluded because 3 out of the 4 nestlings were sired by a male that never fed them (i.e., probably a case of extra-group paternity).  ^4^ Only in one out of eight broods with multiple paternity both sires fed the nestlings. Hence, only this brood was considered as including extra-pair rather than extra-group paternity. Four broods were excluded since the authors could not identify the sire of any of the nestlings. It is assumed to be a group-year sample. **Population:** Two colonies at Armidale, NSW, Australia.  ^5^ According to Barati et al. (Barati *et al.*, 2018b, p. 249) it may be that most groups in this study had helpers related to the breeding female. |  |
| **White-browed scrubwren** (*Sericornis frontalis*) | Females | **Not relevant** ^1^ | | | | | ^1^ No adult female helpers in this species (Magrath & Whittingham, 1997, pp. 187–188; Whittingham, Dunn & Magrath, 1997, p. 262). | - Raihani & Clutton-Brock (2010) - Cornwallis *et al.* (2010) |
|  | Males | Only groups with multiple sexually mature males | 64%  (N = 45 offspring, 19 broods, 10 groups) ^1-2^  (Whittingham *et al.*, 1997, pp. 264–267) ^Genetic evidence^ | 47%  (N = 19 broods, 10 groups) ^1-3^  (Whittingham *et al.*, 1997, pp. 264–267) ^Genetic evidence^ | **Not extreme reproductive skew** | **Limited** | ^1^ Only groups with multiple males in which both males were unrelated to the breeding female were considered (see Table 2 in Whittingham *et al.*, 1997).  ^2^ Five nestlings that were sired by extra-group males were excluded.  ^3^ Percentages are calculated from the number of broods. |  |
| **American bushtit** (*Psaltriparus minimus*) | Females | A mixture of single-female and multi-female groups |  | ≤75%  (N = 12 nests) ^1^  (Sloane, 1996, pp. 760–761) ^Physiological evidence^ | **Not extreme reproductive skew** ^2^ | **Limited** | ^1^ The percentage is calculated from the 12 nests in which there were more than two opposite-sex individuals before or during the egg-laying period. Though, it may be that the same groups with helpers produced more than 1 nest. Hence the number of groups may be smaller. Additionally, some of these extra group members could have been male. 25% of maternity sharing is calculated from the fact that in three of these 12 nests, there were more eggs than is expected by a single female.  ^2^ No group had a female helper in Bruce *et al.* (Bruce *et al.*, 1996, p. 512) parentage study. | - Raihani & Clutton-Brock (2010) - Cornwallis *et al.* (2010) |
|  | Males | **No data** ^1-2^ | | | | | ^1^ Bruce *et al.* (Bruce *et al.*, 1996, pp. 512–513) ^Genetic evidence^ sampled three families with a female and two males. Yet, two of these families (nests 204 and 207) started as pairs and the second male joined only after the first male disappeared during the incubation stage of the clutch. The third family (family P2) started as a pair and the third male joined during the chick-rearing stage. Since these groups were not multi-male groups during the fertilisation stage we did not consider them relevant to our study.  ^2^ But see Sloane (Sloane, 1996, pp. 760–761) for behavioural evidence of mating and reproductive sharing in multi-male groups. |  |
| **Green woodhoopoe** (*Phoeniculus purpureus*) | Females | A mixture of single-female and multi-female groups |  | <96% ^1^  (N = 73 clutches)  (Ligon & Ligon, 1990, pp. 41–42) ^Behavioural and physiological evidence^ | **Extreme reproductive skew** | **Substantial** | ^1^ This estimation is based on the number of clutches that were larger than the maximum clutch size for a single female. It thus should be seen as the maximum percentage of groups with a singular breeding female. | - Raihani & Clutton-Brock (2010) - Cornwallis *et al.* (2010) |
|  | Males | **No data** ^1^ | | | | | ^1^ We were not able to find systematic data about paternity in this species. But see (Ligon & Ligon, 1990, pp. 41–42) for behavioural observations that suggest monopolisation of paternity. |  |
| **Red-cockaded woodpecker** (*Picoides borealis*) | Females | **No data** ^1^ | | | | | ^1^ Only ~2% of groups in this species have a female helper (Walters, Doerr & Carter, 1988, p. 286). In addition, the genetic studies by Haig, Belthoff & Allen (Haig, Belthoff & Allen, 1993, pp. 190–191) and Haig *et al.* (1994, p. 299) did not include groups with female helpers. | - Raihani & Clutton-Brock (2010) - Cornwallis *et al.* (2010) |
|  | Males | Only groups with multiple sexually mature males | 100%  (N = 31 offspring, 16 nest-years) ^1^  (Haig *et al.*, 1994, p. 299) ^Genetic evidence^  100%  (N = 2 offspring, 1 nest) ^2^  (Haig *et al.*, 1993, pp. 190–191) ^Genetic evidence^ | 100%  (N = 16 nest-years) ^1^  (Haig *et al.*, 1994, p. 299) ^Genetic evidence^  100%  (N = 1 nest) ^2^  (Haig *et al.*, 1993, pp. 190–191) ^Genetic evidence^ | **Extreme reproductive skew ^3^** | **Limited** | ^1^ One offspring was excluded from the sample as extra-group paternity. In addition, many of the helpers were the offspring of the male, and probably also of the female, breeders in the group (Table 3). Hence, incest limitation may have been present in most groups. Population: Sandhills region of south-central North Carolina, USA.  **^2^** Only case C in Table 2 was considered as it is the only case in which the male helper was not the offspring of the breeding female. Population: Savannah River Site,  South Carolina, USA.  **^3^** For further behavioural evidence of monogamy see (Lennartz, Hooper & Harlow, 1987, p. 78). |  |
| **White-breasted thrasher** (*Ramphocinclus brachyurus*) | Females | A mixture of single-female and multi-female groups ^1^ | > 56% ^2^  (N = 16 offspring, 7 multi-female groups) ^3^  (Temple, Hoffman & Amos, 2009, pp. 103, 105) ^Genetic evidence^ |  | **No data** ^2-3^ **/ not relevant** ^1^ | **Limited** | ^1^ In all seven groups with a female helper, the helper was the daughter of the breeding pair (Temple *et al.*, 2009, p. 104). In addition, all female helpers were yearlings that may be too young to be sexually mature (Temple *et al.*, 2009, p. 108). Hence, this sample, and the species in general, may not have sexually mature female helpers.  ^2^ 56% of offspring were assigned to the older female in the group (assumed to be the mother of the yearling female helper). The remaining offspring could be assigned to either the older or the yearling female in each group. Because females were mother-daughter, maternity is not clear.  But, see Temple *et al* (Temple *et al.*, 2009, pp. 105–106) for physiological evidence that only the older female in the group breeds.  ^3^ Only 5 of the 7 groups examined had more than one offspring, thereby enabling shared maternity to be detected (Temple *et al.*, 2009, pp. 103, 105). We considered all seven groups because maternity assignments were not indicated for each group. | - Clutton-Brock (2010) - Cornwallis *et al.* (2010) |
|  | Males | A mixture of single-male and multi-male groups ^1^ | > 64% **^2^**  (N = 22 offspring, 10 multi-male groups) ^3^  (Temple *et al.*, 2009, pp. 103, 105) ^Genetic evidence^ |  | **No data** ^2-3^ **/ Not relevant** ^1^ | **Limited** | ^1^ All male helpers were the offspring of the breeding pair (Temple *et al.*, 2009, p. 104). In addition, all helpers, except one male, were yearlings that may be too young to be sexually mature (Temple *et al.*, 2009, p. 108). Hence, the species may not have sexually mature male helpers.  ^2^ 64% of offspring were assigned to the older male in the group. The remaining offspring could be assigned to either the older or the yearling male in each group. Since males were father-son, paternity is not clear.  ^3^ Only 8 of the 10 examined groups had more than one offspring, thereby enabling shared maternity to be detected (Temple *et al.*, 2009). In addition, one nestling was excluded as extra-group paternity. |  |
| **White-winged chough**  (*Corcorax melanorhamphos*) | Females | Only groups with multiple sexually mature females | Not available | 75%  (n = 13 offspring, 4 multi-female groups) ^1^  (Heinsohn *et al.*, 2000, pp. 244–246) ^Genetic evidence^  ≤ 77% ^2^  (n = 52 broods from <30 newly formed and established groups) ^3^  (Beck, 2006, pp. 106–107, 109–111) ^Genetic evidence^ | **Not extreme reproductive skew ^4^** | **Substantial** | **^1^** In newly formed groups after a severe drought (each group had multiple females that were unrelated to at least one male: groups “O’Connor 9”, “Veterans”, “Bullies” and “Village People”).  The mean reproductive skew in these four newly formed groups was 0.895 (1 means complete monopolisation of breeding by a single male). There were only two socially established groups with multiple sexually mature females (“Red” and “Tidbinbilla”). But there was no breeding in these groups. **Population:** Campbell Park, Black Mountain and Tidbinbilla Nature Reserves, ACT, (around Canberra), Australia.  **^2^** This is a maximum estimation due to incomplete sampling of all offspring within a brood. Hence, it may be that shared maternity within some broods was missed. In addition, a single female reproduced in 23% of new groups, but in 85% of established groups. Since the number of broods from each group category is not indicated, we did not indicate these estimations but used the overall percentage. Population: Canberra Nature Park and neighbouring suburban areas of Canberra, Australia.  **^3^** This sample is from a combined group of broods, including those from newly formed and socially established groups. Both types of groups included multiple sexually mature females and more than one sampled offspring.  In addition, 3 broods were excluded from our analysis as cases of extra-group maternity and parasitism. Since it is not clear from which groups these excluded broods come, the sample size of groups is 27-30 groups. The percentage is calculated from the number of groups. In established groups, most helpers were the offspring of the breeding pair and, thus, reproductive skew may occur due to incest avoidance (Beck, 2006, p. 118). For the relatively high fitness value of offspring hatched in bad years see (Shen *et al.*, 2017).  **^4^** See also Rowley (1978, pp. 180–181) for evidence that 15% of broods involved polygyny (multiple breeding females in a group), based on larger-than-average clutch size in 203 broods. This is a minimum estimation since some of these broods may have come from groups with a single female. For similar evidence, though more anecdotical, see also North (North, 1901, pp. 21–22). | - Raihani & Clutton-Brock (2010) - Cornwallis *et al.* (2010) |
|  | Males | Only groups with multiple sexually mature males | Not available | 80%  (n = 22 offspring, 10 multi-male group-years, 5 groups) **^1^**  (Heinsohn *et al.*, 2000, pp. 244–246) ^Genetic evidence^  38%  (n = 26 offspring, 8 multi-male groups) **^2^**  (Heinsohn *et al.*, 2000, pp. 244–246) ^Genetic evidence^  ≤ 87% **^3^**  (n = 54 broods from <30 newly formed and established groups) **^4^**  (Beck, 2006, pp. 106–107, 112–114) ^Genetic evidence^ | **Equivocal ^3;^ ^5^** | **Substantial** | **^1^** In socially established groups (with multiple males that were unrelated to at least one female). The sample size is the number of groups. We only considered groups with multiple sexually mature males that had offspring. Namely, groups “Farmers”, “Yellow”, Campbell Park Blue”, and “Green”. Note that group “Green” was considered twice, before and after a new unrelated male joined the group and sired an offspring which was the only case of shared paternity. The sample size of offspring is 20-22 offspring since 3 group members in the “Yellow” group were not sampled for DNA and 2 out of 4 adults were sampled (meaning, a minimum of one offspring was not sampled for DNA, a maximum of 3 offspring were not sampled). Population: Campbell Park, Black Mountain and Tidbinbilla Nature Reserves, ACT, (around Canberra), Australia.  **^2^** In newly formed groups after a severe drought (each group had multiple males that were unrelated to at least one female). The sample size is the number of groups. The mean reproductive skew in these groups was 0.64 (1 means complete monopolisation of breeding by a single male). Group “Haig Park” was excluded since there was only one offspring in this group so paternity could not be shared. Population: Campbell Park, Black Mountain and Tidbinbilla Nature Reserves, ACT, (around Canberra), Australia.  **^3^** This is a maximum estimation due to incomplete sampling of all offspring within a brood. Therefore, shared paternity could be missed in some broods. In addition, a single male reproduced in 57% of new groups (Table 2, p. 114), but in 100% of established groups (Beck, 2006). However, since the number of broods from each group category is not indicated, we did not indicate these estimations. But, rather present the overall percentage (i.e., 87%). Population: Canberra Nature Park and neighbouring suburban areas of Canberra, Australia.  **^4^** This sample is from a combined sample of broods from newly formed and established groups with multiple sexually mature males and more than one sampled offspring. In addition, 1 brood was excluded from our analysis as a case of extra-group paternity. Since it is not clear from which groups these excluded broods come, the sample size of groups is 29-30 groups. In addition, in established groups, most helpers were the offspring of the breeding pair and, thus, reproductive skew may occur due to incest avoidance (Beck, 2006, p. 118).  **^5^** It seems like there is high male reproductive skew in socially established groups, but not in newly established groups. Yet, for the relatively high fitness value of those offspring that hatch in “bad” years see (Shen *et al.*, 2017). |  |
| **Superb starling** (*Lamprotornis superbus*) | Females | A mixture of single-female and multi-female groups **^1^** |  |  | **Not extreme reproductive skew ^2^** | **Substantial** **^2^** | **^1^** >90% of groups had at least one helper and most helpers were males. In addition, most social groups include more than one breeding pair (see comment number 2). Yet the reported results do not necessarily include only groups with multiple sexually mature females (Rubenstein, 2007, p. 1896).  **^2^** Only nesting females lay eggs in their nest (Rubenstein, 2016, p. 187), but there are up to 6 monogamous breeding pairs (mean = 3.5) in the long rain season and up to 4 monogamous breeding pairs (mean of 1.7) in the short rain season.  (N = 247 offspring, 100 nests, 9 groups) (Rubenstein, 2007, p. 1896, 184 2016). See also Rubenstein (Rubenstein, 2016, p. 189) where it is indicated that 25% of group members breed. Population: Mpala Research Centre, Laikipia, Kenya. | - Cornwallis *et al.* (2010) |
|  | Males | A mixture of single-male and multi-male groups **^1^** | 92%  (N = 231 offspring, 100 nests, 9 groups) **^2^**  (Rubenstein, 2007, pp. 1897–1898) ^Genetic evidence^ | 84%  (N = 100 nests, 9 groups)  (Rubenstein, 2007, p. 1898) ^Genetic evidence^ | **Not extreme reproductive skew** **^3^** | **Substantial** | **^1^** >90% of groups had at least one helper and most helpers were males. In addition, see comment 3 for evidence that groups include multiple monogamous pairs. But the reported results do not necessarily include only groups with multiple sexually mature males (Rubenstein, 2007, p. 1896).  **^2^** 16 offspring were excluded as extra-group paternity (Rubenstein, 2007, p. 1898).  **^3^** Superb starlings are plural breeders living in large groups (10-35 birds) that include up to 6 breeding pairs (mean = 3.5) in the long rain season and up to 4 monogamous breeding pairs (mean of 1.7) in the short rain season (N = 247 offspring, 100 nests, 9 groups) (Rubenstein, 2007, p. 1896, 184 2016). Hence, besides the minimum threshold of 8% reproduction by within-group helpers, many other group members are breeders (Rubenstein, 2007, 2016, p. 184). See also Rubenstein (Rubenstein, 2016, p. 189) where it is indicated that 25% of group members breed. Population: Mpala Research Centre, Laikipia, Kenya.  See also (Weinman, Solomon & Rubenstein, 2015, p. 504) for genetic evidence of a high percentage of extra-pair paternity in the same population, but from a greater larger sample. Yet, this sample does not differentiate between extra-pair paternity by males within and outside the group. |  |
| **Australian magpie** (*Gymnorhina tibicen*) | Females | Only groups with multiple sexually mature females |  | 0%  (N = 14 group-years, 6 multi-female groups) **^1, 2^**  (Hughes *et al.*, 2003, pp. 3445–3446) ^Genetic evidence^  67%  (N = 15 multi-female groups with more than 1 offspring that could be assigned to within-group females) ^3^  (Durrant & Hughes, 2005, pp. 541–542) ^Genetic evidence^ | **Not extreme reproductive skew ^5^** | **Substantial** | **^1^** Maternity sharing was assessed across years within the same group. The sample size is, therefore, groups. Yet, even when considering only group-years with multiple females and more than one offspring, there is shared maternity in 100% of groups (N = 10 group years, 5 groups).  Maternity is estimated across years within the same group because dominance was not assessed in this study. Population: Guildford on the Swan River at the northeastern edge of Perth, Western Australia (lat. 31°54′south, long. 115°59′ east).  **^2^** Only groups with multiple females and with more than 1 offspring that could be assigned to within-group females were considered (i.e., groups FM, HC, HR, MM, NH, PG1).  **^3^** Only groups with multiple females, more than one offspring that was assigned within-group female according to genetics and which had the same social composition over multiple years were included. Namely, the following groups were included: “AS”, “BAC”, “BPP”, “FW”, “GK01”, “HB”, “JD”, “LB”, “PC”, “RR”, “SD”, “SS”, “SW”, “TR”, “VE”. Population: Rowsley, 50 km west of Melbourne, Victoria, Australia (37°43°’S, 144°24°’E).  **^4^** Population: Seymour (37°2’S, 145°9’E), Australia. Although all groups had more than 2 adults, it is not clear whether all groups had multiple females.  ^5^ See also Carrick (1972, pp. 45, 66–67) for behavioural evidence that only 19% of 1,117 adult females did not breed. Population: Canberra (35°15' North of Canberra), Australia.  See also (Fulton, 2006, pp. 198–201) for a case of plural breeding even within the same tree in a population in Dryandra Woodland, south-east of Perth, Western Australia (32°46′S, 116°58′E). | - Cornwallis *et al.* (2010) |
|  |  | A mixture of single-female and multi-female groups **^4^** |  | 25%  (N = 36 groups)  (Hughes *et al.*, 1996, pp. 67–68) ^Behavioural evidence^ |  |  |  |  |
|  | Males | Only groups with multiple sexually mature males | 50%  (N = 2 offspring, 2 group-years, 1 multi-male group) **^1^**  (Hughes *et al.*, 2003, pp. 3445–3446) ^Genetic evidence^ | 0%  (N = 1 multi-male group with more than 1 offspring that could be assigned to within-group males) **^1^**  (Hughes *et al.*, 2003, pp. 3445–3446) ^Genetic evidence^  50%  (N = 2 multi-male groups with more than 1 offspring that could be assigned to within-group males) ^2^  (Durrant & Hughes, 2005, pp. 541–542) ^Genetic evidence^ | **Not extreme reproductive skew** | **Anecdotical ^3^** | **^1^** Only in one group (HR) there were multiple males and more than one offspring that could be assigned to within-group males (Hughes *et al.*, 2003, p. 3445). Namely, only in one group shared paternity could be detected. Population: Guildford on the Swan River at the northeastern edge of Perth, Western Australia (lat. 31°54′south, long. 115°59′ east).  ^2^ Only groups “CT” and “HHH” had more than one offspring that could be assigned paternity and multiple males. The parentage proxy was not calculated for this study because the dominance categories of the males were not assessed. Population: Rowsley, 50 km west of Melbourne, Victoria, Australia (37°43°’S, 144°24°’E).  ^3^ In both studies paternity sharing was assessed across years of the same group. The sample size is therefore groups. |  |
| **Splendid** **fairy-wren**  (*Malurus splendens*) | Females | Only groups with multiple sexually mature females |  | 69% **^1^**  (N = 106 brood-years)  (Rowley *et al.*, 1989, pp. 232–233) ^Behavioural evidence^  0%  (N = 1 brood) **^1, 2^**  (Brooked *et al.*, 1990, pp. 196–199) ^Genetic evidence^ | **Equivocal**  (population dependent) **^6^** | **Substantial** | **^1^** Population: Gooseberry hill, east of Perth, Australia.  **^2^** Group HH was the only one with two adult females and both bred.  **^3^** 89.7% of groups studied over 6 years (N = 319 group-years) had a single sexually mature female over the breeding season. Hence, there are very few groups with multiple females in this species (Webster *et al.*, 2004, p. 908). See also (Van Bael & Pruett-Jones, 2000, p. 98) for group composition over 132 group-years. Population: Brookfield Conservation Park, South Australia.  **^4^** Most probably the 3 cases of maternity mismatch with the dominant female in the group resulted from brood parasitism from outside the social group (Webster *et al.*, 2004, p. 911). See table 3 for the number of broods.  **^5^** Assuming the three cases of maternity mismatch with the alpha female came from three different broods. See also Van Bael & Pruett-Jones (2000, p. 105) for evidence of the rarity of plural breeding among females in the population of Brookfield Conservation Park, South Australia.  **^6^** In the Gooseberry hill population there was no extreme female reproductive skew. But in the Brookfield Conservation Park, South Australia population, there is extreme female reproductive skew. In both populations, though there were only a few groups with multiple females. | - Raihani & Clutton-Brock (2010) - Cornwallis *et al.* (2010) |
|  |  | A mixture of single-female and multi-female groups **^3^** | 99% **^4^**  (N = 450 nestlings, 159 brood-years)  (Webster *et al.*, 2004, pp. 908, 911) ^Genetic evidence^ | 98% **^4-5^**  (N = 159 brood-years)  (Webster *et al.*, 2004, pp. 908, 911) ^Genetic evidence^ |  |  |  |  |
|  | Males | Only groups with multiple sexually mature males | 70% **^1^**  (N = 23 nestlings, 9 brood-years) ^2^  (Brooked *et al.*, 1990, pp. 193, 196–199) ^Genetic evidence^ | 50%  (N = 9 brood-years, 8 groups) **^2^**  (Brooked *et al.*, 1990, pp. 196–199) ^Genetic evidence^  76%  (N = 41 brood-year) **^3^**  (Webster *et al.*, 2004, p. 912) ^Genetic evidence^  60%  (N = 20 brood-year) **^4^**  (Webster *et al.*, 2004, p. 912) ^Genetic evidence^ | **Not extreme reproductive skew ^5^** | **Substantial** | ^1^ Yet, since the probability to detect paternity by male helpers is 0.65, the estimated paternity of within-group helpers is 47%.  ^2^ Only eight multi-male groups in which at least 1 offspring was assigned to a within-group male were considered. Namely, only groups “CC” (2 years), “H”, “HH”, “N”, “Q”, “R”, “S”, and “T”. The percentage of groups with a singular breeder was calculated from the number of groups.  Population: Gooseberry hill, east of Perth, Australia.  ^3^ Out of 41 broods in which it could be inferred whether the auxiliary male was the son of the breeding female or not. In 20 and 21 of these brood-years, the male helper was not / was the son of the breeding female, respectively.  Male helpers were significantly more likely to breed if they were unrelated to the female in the group.  Population: Brookfield Conservation Park, South Australia.  **^4^** This is a sub-sample of the sample presented in comment 3. This sample includes only the 20 broods in which the auxiliary male was not the son of the breeding female and thus there was no incest limitation. **Population:** Brookfield Conservation Park, South Australia.  **^5^** See also Figure 1 in (Webster *et al.*, 2007) for further genetic evidence for “auxiliaries” breeding in the same sample of Webster *et al.* (2004, pp. 908, 911). |  |
| **Acorn woodpecker** (*Melanerpes formicivorus*) | Females | Only groups with multiple sexually mature females | 58 ± 2% **^1^**  (mean ± SE)  (N = 11 groups) **^2^**  56% ± 6% **^1^**  (mean ± SE)  (N = 2 groups) **^3^**  (Haydock, Koenig & Stanback, 2001, pp. 1520–1521 Table 6) ^Genetic evidence^ | 8%  (N = 13 sets of co-breeder females) **^4^**  (Haydock *et al.*, 2001, pp. 1520–1521) ^Genetic evidence^  19%  (N = 27 nests) **^5^**  (Haydock *et al.*, 2001, pp. 1520–1521) ^Genetic evidence^ | **Not extreme reproductive skew ^6^** | **Substantial ^6^** | **^1^** These percentages (mean and SE) of offspring produced by the more successful female in the group may underestimate reproduction by the most dominant females as they do not include groups in which reproduction was monopolised by a single female. Nonetheless, according to table 4 (Haydock *et al.*, 2001, p. 1519) only 3 of 54 groups were like this and these groups produced only 14 out of the 400 offspring examined. Population: Hastings Natural History Reservation, central coastal California, USA.  **^2^** In groups with two females.  **^3^** In groups with more than two females.  **^4^** This sample includes 13 sets of co-breeding females that reproduced together multiple times. In 12 of these sets, maternity was shared between the females. Population: Hastings Natural History Reservation, central coastal California, USA.  **^5^** In nests with more than one offspring for which maternity was assigned. Population: Hastings Natural History Reservation, central coastal California, USA.  **^6^** See also Haydock & Koenig (2002) and Haydock & Koenig (2003, p. 278)for similar results from an extended sample which partly overlaps with the results from (Haydock *et al.*, 2001). | - Cornwallis *et al.* (2010) |
|  | Males | Only groups with multiple sexually mature males | 77 ± 4% ^1^  (N = 25 groups) **^2^**  (mean ± SE)  69% ± 7% **^1^**  (mean ± SE)  (N = 15 groups) **^3^**  (Haydock *et al.*, 2001, p. 1520 Table 5) ^Genetic evidence^ | 47%  (N = 40 sets of co-breeder males) **^4^**  (Haydock *et al.*, 2001, p. 1520) ^Genetic evidence^  66%  (N = 91 nests) **^5^**  (Haydock *et al.*, 2001, p. 1520) ^Genetic evidence^  0  (N = 1 group) **^6^**  (Joste, Ligon & Stacey, 1985, pp. 39–40) ^Genetic evidence^ | **Not extreme reproductive skew ^7^** | **Substantial** | **^1^** These percentages (mean and SE) of offspring produced by the more successful male in the group may underestimate reproduction by the most dominant males as they do not include groups in which reproduction was monopolised by a single male. Nonetheless, according to table 4 (Haydock *et al.*, 2001, p. 1519) only 3 of 54 groups were like this and these groups produced only 14 out of the 400 offspring examined. Population: Hastings Natural History Reservation, central coastal California, USA.  **^2^** In groups with two males  **^3^** In groups with more than two males.  **^4^** This sample includes 40 sets of co-breeding males that reproduced together multiple times. In 25 of these sets, paternity was shared between the males. Population: Hastings Natural History Reservation, central coastal California, USA.  **^5^** In nests with more than one offspring for which paternity was assigned.  **^6^** Two other multi-male groups were tested in this study but were not sampled completely and one male remained unsampled. The results from both of these groups suggest multiple paternity as well. Population: Water Canyon in the Magdalena Mountains, Socorro County, New Mexico, USA.  **^7^** See also (Haydock & Koenig, 2002) and (Haydock & Koenig, 2003, p. 278)for similar results from an extended sample which partly overlaps with the results from (Haydock *et al.*, 2001). |  |
| **Common moorhen** (*Gallinula chloropus*) | Females | Only groups with multiple sexually mature females | 58%  (N = 90 offspring, 10 broods, 6 groups)  (McRae, 1996, p. 231; Table IV) ^Genetic evidence^ | <30%  (N = 46 broods) **^1^**  (McRae, 1996, p. 231; Table IV) ^Genetic and physiological evidence^ **^2^** | **Not extreme reproductive skew ^3^** | **Substantial** | **^1^** Overall, 46 broods with multiple females were observed during the same 3 years (McRae, 1996, p. 229). Thereof broods, 14 females helped and did not reproduce (some of these 14 females reproduced later in the same group) (McRae, 1996, p. 238). As it is not clear from how many different groups the 14 non-reproducing female helpers were, the maximum percentage of broods with a non-reproducing female helper is presented (i.e., assuming that the 14 non-reproducing females were from 14 different groups). Population: Peakirk Waterfowl Gardens, Cambridge-shire, U.K.  **^2^** In some cases maternity was inferred by eggshell patterns, and in other cases by DNA fingerprints.  **^3^** Mean female “reproductive skew” was 0.59 (N = 418 eggs, 19 year-groups, 14 groups). Reproductive skew in this study was calculated as the proportion of each female’s eggs in the nest. Namely, the ”skew” represents a divergence from equal sharing regardless of the female’s dominance (i.e., equal sharing means 0.5) (McRae, 1996, pp. 236–237, Table VI).  See also (Loyau & Schmeller, 2012, p. 677) for some evidence of plural breeding in France. | - Cornwallis *et al.* (2010) |
|  | Males | Only groups with multiple sexually mature males | 65% **^1^**  (N = 20 offspring, 3 broods, 3 multi-male groups) **^2^**  (McRae, 1996, pp. 231–232, Table IV, 238) ^Genetic evidence^ | 0% **^1^**  (N = 3 broods, 3 multi-male groups) **^2^**  (McRae, 1996, pp. 231–232, Table IV, 238) ^Genetic evidence^ | **Not extreme reproductive skew** | **Anecdotical** | **^1^** The younger male of the 2 sexually mature males in the group was considered the “subordinate” male (McRae, 1996, p. 230). The sample size of three groups consists of: one group from 1992 (Table IV on pages 231-232), and another two groups in which the “auxiliary” male was the son of the breeding female (McRae, 1996, p. 238).  **Population:** Peakirk Waterfowl Gardens, Cambridge-shire, U.K.  **^2^** In two of the three groups the younger male was the son of the breeding female (McRae, 1996, p. 238). |  |
| **Seychelles warbler** (*Acrocephalus sechellensis*) | Females | Only groups with multiple sexually mature females | 48%  )N = 29 offspring, 15 multi-female groups) **^1^**  (Groenewoud *et al.*, 2018, pp. 1257–1258, Table 2)  ^Genetic evidence^  56%  )N = 18 offspring, 12 multi-female groups)  (Richardson *et al.*, 2001, p. 2267) ^Genetic evidence^  66%  )N = 314 offspring, >30 groups) **^2^**  (Raj Pant *et al.*, 2019, p. 1257) ^Genetic evidence^ | 40%  )N = 5 multi-female groups) **^3^**  (Richardson *et al.*, 2001, pp. 2267–2268) ^Genetic evidence^  42% - 75% **^4^**  )N = 12 multi-female groups)  (Richardson *et al.*, 2001, pp. 2267–2268) ^Genetic evidence^  47% **^5^**  )N = 15 multi-female groups) **^1^**  (Groenewoud *et al.*, 2018, pp. 1257–1258, Table 2)  ^Genetic evidence^ | **Not extreme reproductive skew ^6^** | **Substantial** | **^1^** This sample includes only groups with multiple sexually mature females in which the subordinate female immigrated to the group (i.e., no incest limitation). 6 individuals that were only present in their new non-natal group for a short time (“stagers”) were excluded. We assume that each female was present in a different group and, therefore, estimated the number of groups as 15. This sample is from 2002-2015 and thus partly overlaps with (Raj Pant *et al.*, 2019, p. 1257). Population: Cousin Island (29 ha, 04°20′S, 55°40′E).  **^2^** This sample probably also includes the smaller sample from Richardson *et al.* (2001, p. 2267). We estimated the total number of offspring to be between 312 and 316 because it is indicated that 106 offspring were produced by subordinate females and that this number was equal to 34% of all offspring produced in multi-females groups. The number of groups is not indicated but is more than 30 groups. Because no nest contained more than 3 eggs and usually each female produced only one egg. This sample is from 1997-2014. Population: Cousin Island (29 ha, 04°20′S, 55°40′E).  **^3^** In groups in which mixed maternity could be detected (i.e., >1 sampled offspring). Population: Cousin Island (04°20′S, 55°40′E).  **^4^** In these 12 multi-female groups there were 16 female helpers. 7 of these female helpers gained maternity, but it is not indicated from how many groups they were. Hence, the range presents the minimum and maximum possibilities. Namely, that the 7 breeding helpers were from 7 different groups (i.e., 42% = minimum percentage of groups with a singular breeder) or that these helpers were from 3 different groups (i.e., 75% = maximum percentage of groups with a singular breeder).  Three is the minimum number of groups with helpers as each of the 12 groups must have had at least one helper and then one group could have all the rest helpers (i.e., 4). Then, it may be that all these 5 helpers in one group gain maternity in addition to female helpers from another 2 groups. Population: Cousin Island (29 ha, 04°20′S, 55°40′E).  **^5^** This is a maximum estimation as there may have been no breeding in some of these groups. For example, in the groups in which “stagers” were present, there was no breeding during their presence (Table 2).  **^6^** See also (Komdeur, 2005) for physiological and behavioural evidence for joint laying in multi-female groups. See (Richardson, Komdeur & Burke, 2003, p. 580) for genetic evidence that 8 out of 21 subordinate females produced offspring in their group. | - Cornwallis *et al.* (2010) |
|  | Males | Only groups with multiple sexually mature males | 83%  )N = 6 offspring, 6 multi-male groups)  (Richardson *et al.*, 2001, pp. 2267–2268) ^Genetic evidence^ | 83%  )N = 6 multi-male groups)  (Richardson *et al.*, 2001, pp. 2267–2268) ^Genetic evidence^  70% - 90% **^1^**  (N = 10 - 20 breeding events, each breeding event is a group) **^2^**  (Richardson, Burke & Komdeur, 2002, p. 2315) ^Genetic evidence^ | **Not extreme reproductive skew** | **Substantial** | **^1^** In total, there were 68 breeding events with a subordinate male and/or female. Also, these 68 breeding events involved a total of 20 subordinate males.  Of these breeding events, 37 events involved 1 subordinate, 14 events involved 2 subordinates and 1 event involved 3 subordinates. Since the exact number of breeding events with subordinate males is not indicated, we estimated the minimum and maximum possible numbers of breeding events with subordinate males. A minimum number of 10 breeding events with subordinate males assumes that 1 breeding event involved 3 subordinate males, 8 breeding events involved 2 subordinate males each and 1 breeding event involved 1 subordinate male. On the contrary, a minimum number of 20 breeding events with subordinate males assumes that 20 breeding events involved 1 subordinate male.  In addition, the percentages are probably an overestimation of breeding by dominant males since the sample does not exclude extra-group paternity, which is about 40% of nestlings in this species.  **^2^** This sample is from the years 1997-1999 and thus partly overlaps with the sample of (Richardson *et al.*, 2001, pp. 2267–2268) which is from 1999. In addition, this sample includes only breeding events with at least one nestling, in this species most of the clutches include a single egg.  Population: Cousin Island (29 ha, 04°20′S, 55°40′E). |  |
| **Carrion crow** (*Corvus corone*) | Females | Only groups with multiple sexually mature females | 40% - 60% **^1^**  (N = 5 offspring, 1 multi-female group with an unrelated male) **^2^**  (Baglione *et al.*, 2002, pp. 889, 891) ^Genetic evidence^ | 0%  (N = 1 multi-female group with an unrelated male) **^2^**  (Baglione *et al.*, 2002, pp. 889, 891) ^Genetic evidence^ | **Not extreme reproductive skew ^3^** | **Anecdotical** | **^1^** Two offspring were produced by one female and three offspring were produced by another female. Since the dominance ranks of the mothers are unknown, we present the range of possibilities. Namely, the minimum possibility that the dominant female produced only 2 out of the 5 offspring (i.e., 40% of offspring produced by the alpha female), and the maximum possibility that the dominant female produced 3 out of the 5 offspring (i.e., 60% of offspring produced by the alpha female).  **^2^** Group 3 is the only relevant group as it had multiple females with an unrelated male and more than one offspring was sampled, so multiple maternity could be detected. Population: La Sobarriba, northern Spain (42 N, 5 W).  **^3^** See also (Canestrari, Marcos & Baglione, 2005) for genetic evidence of substantial shared parentage. | - Raihani & Clutton-Brock (2010) - Cornwallis *et al.* (2010) |
|  | Males | Only groups with multiple sexually mature males | 73% - 87% **^1^**  (N = 15 offspring, 5 multi-male groups with an unrelated female) **^2^**  (Baglione *et al.*, 2002, pp. 889, 891) ^Genetic evidence^ | 60%  (N = 5 multi-male groups with an unrelated female) **^2^**  (Baglione *et al.*, 2002, pp. 889, 891) ^Genetic evidence^ | **Not extreme reproductive skew ^3^** | **Anecdotical** | **^1^** In each of the 2 groups with multiple paternity (67, 70) there were 3 offspring. Since the dominance ranks of the fathers were unknown, we present the range of minimum and maximum possibilities. Namely, the minimum possibility is that the dominant male produced only 11 out of the 15 offspring (i.e., 73% of offspring produced by the alpha male; the subordinate male sired 2 offspring in each of these groups). And the maximum possibility is that the dominant male produced 13 out of the 15 offspring (i.e., 87% of offspring produced by the alpha male; the subordinate male sired only 1 offspring in each of these groups).  **^2^** Two groups with evidence for shared paternity were excluded, as it could not be determined whether paternity was shared within the group or with extra-group males (groups 14 and 27). Groups 3 and 85 were excluded as paternity could not be assigned to a specific male.  Only groups with multiple males that were unrelated to a mature female were included in the analysis (i.e., groups 70, 82, 68, 87, 67). Population: La Sobarriba, northern Spain (42 N, 5 W).  **^3^** See also (Canestrari *et al.*, 2005) for genetic evidence of substantial shared parentage. |  |
| **Mexican jay** (*Aphelocoma ultramarine*) | Females | Only groups with multiple sexually mature females | **Not available** | 51%  (N = 47 group-years, 10 groups sampled over 6 years) **^1-2^**  (Li, 1997 Appendix C) ^Genetic evidence^ | **Not extreme reproductive skew ^3^** | **Substantial** | **^1^** We only considered the following groups-years:  Group 11 (years: 1991-1995),  Group 12 (years: 1991, 1993-1995),  Group 13 (years: 1991-1996),  Group 16 (years: 1991-1994),  Group 17 (years: 1991-1996),  Group 18 (years: 1991-1994, 1996),  Group 19 (years: 1991-1996),  Group 20 (years: 1991- 1995),  Group 21 (years: 1991- 1992, 1994 - 1995),  Group 22 (years: 1991, 1994).  Birds older than 2 years were considered sexually mature. Group years with only one nestling were excluded (e.g., group 16 in 1995). The percentage was calculated from group-years, as group composition changed from year to year. Population: Portal, Arizona at the Southwestern Research Station of the American Museum of Natural History and the surrounding Coronado National Forest; USA (Latitude 31.883 N, longitude 109.203 W).  **^2^** See also (Li & Brown, 2000, pp. 868, 872) Genetic evidence for a publication that is based on this dataset.  **^3^** Usually, groups have multiple females and several females breed simultaneously in the same group. Hence, it is probably the case that most adult females breed in their group but not necessarily simultaneously (Li & Brown, 2000, pp. 867–868).  **^4^** This is the percentage of group-years with more than one nesting pair. This is a maximum value since in this species there are frequently several nesting pairs and not all nests within each group-year were examined.  This sample partly overlaps with the sample of (Li & Brown, 2000, pp. 868, 872). Namely, the 11 nests from years 1993-1995 were probably considered in both studies. Population: Portal, Arizona at the Southwestern Research Station of the American Museum of Natural History and the surrounding Coronado National Forest; USA (Latitude 31.883 N, longitude 109.203 W). | - Cornwallis *et al.* (2010) |
|  |  | A mixture of single-female and multi-female groups **^3^** |  | ≤81% **^4^**  (N = 31 group-years, 10 groups)  (Eimes, 2004, p. 36; Table 1; see also Eimes *et al.*, 2005) ^Genetic evidence^ |  |  |  |  |
|  | Males | Only groups with multiple sexually mature males | **Not available** | 35%  (N = 48 group-years, 10 groups sampled over 6 years) **^1^**  (Li, 1997 Appendix C) ^Genetic evidence^ | **Not extreme reproductive skew ^6^** | **Substantial** | **^1^** We only considered the following groups-years:  Group 11 (years: 1991-1995),  Group 12 (years: 1991, 1993-1995),  Group 13 (years: 1991-1993, 1995-1996),  Group 16 (years: 1991-1994),  Group 17 (years: 1991-1994, 1996),  Group 18 (years: 1991-1996),  Group 19 (years: 1991-1996),  Group 20 (years: 1991-1995),  Group 21 (years: 1991-1995),  Group 22 (years: 1991, 1993, 1995). Birds older than 2 years were considered sexually mature. Group years with only one nestling were excluded (e.g., group 16 in 1995). The percentage was calculated from group-years, as group composition changed from year to year. See also (Li & Brown, 2000, pp. 870, 874) for a publication that is based on this dataset.  **^2^** Most groups have multiple males that were unrelated to at least one female (Li & Brown, 2000, pp. 867, 874).  **^3^** Three offspring were excluded because it was not clear whether the father was the dominant male or one of the helpers in the group. Additional two offspring were excluded as extra-group paternity (Li & Brown, 2000, pp. 870, 874).  **^4^** This is the maximum value, as there are often multiple nesting pairs in this species and not all nests within each group-year were examined. We excluded six offspring sired by a male other than the female's social mate from our calculation because it was impossible to determine whether the father was a male within or outside of the group.  **^5^** This is the percentage of groups with more than one nesting pair. This is a maximum value since in this species there are frequently several nesting pairs and not all nests within each group-year were examined. In addition, in at least 29% of the nests there was at least one offspring sired by another male from within the social group (Eimes, 2004, p. 26; 11 out of 38 nests for which EPF could be assigned to a father).  This sample partly overlaps with the sample of (Li & Brown, 2000, pp. 868, 872). Namely, the 11 nests from the years 1993-1995 were probably considered in both studies**. Population:** Portal, Arizona at the Southwestern Research Station of the American Museum of Natural History and the surrounding Coronado National Forest; USA (Latitude 31.883 N, longitude 109.203 W).  **^6^** See also (Bowen, Koford & Brown, 1995) for genetic data in the same population (Portal, Arizona at the Southwestern Research Station of the American Museum of Natural History and the surrounding Coronado National Forest; USA (Latitude 31.883 N, longitude 109.203 W), but from an earlier period. |  |
|  |  | A mixture of single-male and multi-male groups **^2^** | 61%  (N =137 offspring, 52 broods, 10 groups sampled over 5 years) **^3^**  (Li & Brown, 2000, pp. 870, 874) ^Genetic evidence^  ≤86% **^4^**  (N = 110 offspring, 31 group-years, 10 groups)  (Eimes, 2004, p. 26; see also Eimes *et al.*, 2005) ^Genetic evidence^ | ≤81% **^5^**  (N = 31 group-years, 10 groups)  (Eimes, 2004, p. 36; Table 1; see also Eimes *et al.*, 2005) ^Genetic evidence^ |  |  |  |  |
| **Subdesert mesite**  (*Monias benschi*) | Females | Only groups with multiple sexually mature females | 79% ^1 -2^  (N = 14 offspring, 7 group-years, 6 multi-female groups)  (Seddon *et al.*, 2005, pp. 3577, 3582) ^Genetic evidence^ | 57% ^1 -2^  (N = 7 group-years, 6 multi-female groups) ^3^  (Seddon *et al.*, 2005, pp. 3577, 3582) ^Genetic evidence^ | **Not extreme reproductive skew** | **Limited** | ^1^ No distinction between dominant and subordinate individuals, so the observed offspring could belong to any of these dominance categories.  ^2^ We included in our calculation only multi-female groups with more than 1 offspring, for which maternity was assigned to within group females (i.e., groups P4, P6, P8a and b, P10, P12, M8). Group composition was inferred from appendix 1, and paternity was inferred from Table 1).  ^3^ Percentages are calculated from the group-years. | - Cornwallis *et al.* (2010) |
|  | Males | Only groups with multiple sexually mature males | 90% ^1 -2^  (N = 10 offspring, 5 multi-male groups)  (Seddon *et al.*, 2005, pp. 3577, 3582) ^Genetic evidence^ | 80% ^1 -2^  (N = 5 multi-male groups)  (Seddon *et al.*, 2005, pp. 3577, 3582) ^Genetic evidence^ | **Not extreme reproductive skew** | **Anecdotal** | ^1^ No distinction between dominant and subordinate individuals, so observed offspring could belong to any of these dominance categories.  ^2^ Only multi-male groups with more than one offspring for which paternity was assigned to within group male/s were included (i.e., groups P4, P8b, P10, P12, M9). Group composition was inferred from appendix 1, and paternity was inferred from Table 1). |  |
| **Florida scrub jay**  )*Aphelocoma coerulescens*( | Females | A mixture of single-female and multi-female groups | 100%  (N = 771 offspring, 279 group-years, 165 groups, three sites) **^1^**  (Townsend *et al.*, 2011a, p. 468) ^Genetic evidence^ | 96.2%  (N = 314 groups) **^2^**  (Windsor *et al.*, 2021, p. 487) ^Genetic and behavioural evidence^ | **Extreme reproductive skew** | **Substantial** | **^1^** Only 17-29% of groups in these populations had an unrelated male or female helper (Townsend *et al.*, 2011a, p. 467) ^Genetic evidence^. **Populations:** (i) Archbold Biological Station (27.10°N, 81.21°W); (ii)  Avon Park Air Force Range (27.41°N, 81.17°W); (iii) Placid Lakes Estates (27.16°N, 81.24°W). **Period:** 2004-2007.  **^2^** The total sample size was calculated as 302 breeding pairs with helpers + 12 polygynous triads with and without helpers (see supplementary materials in (Windsor *et al.*, 2021) for evidence that co-breeders provide alloparental care). This is a mixed sample as it is not indicated whether the helpers were males or females.  **Populations and periods:** (i) Archbold Biological Station (27.10°N, 81.21°W) (1969–2019); (ii)  Avon Park Air Force Range (27.41°N, 81.17°W) (1993–2019); (iii) Placid Lakes Estates (27.16°N, 81.24°W) (1992–2010).  (Townsend *et al.*, 2011a, p. 468) ^Behavioural evidence^ was not considered as the sample from (Windsor *et al.*, 2021) includes also the former sample while distinguishing between pairs and groups. | - Raihani & Clutton-Brock (2010) - Cornwallis *et al.* (2010) |
|  | Males | A mixture of single-male and multi-male groups **^1^** | ≥99.87  (N = 770 offspring, 279 group-years, 165 groups, three sites) **^2^**  (Townsend *et al.*, 2011a, p. 468) ^Genetic evidence^ | Not available | **Extreme reproductive skew** | **Substantial** | ^1^ Only 17-29% of groups in these populations had an unrelated male or female helper (Townsend *et al.*, 2011a, p. 467) ^Genetic evidence^.  ^2^ One offspring that was sired by an extra-group male was excluded.  **Populations:** (i) Archbold Biological Station (27.10°N, 81.21°W); (ii)  Avon Park Air Force Range (27.41°N, 81.17°W); (iii) Placid Lakes Estates (27.16°N, 81.24°W). **Period:** 2004-2007. |  |
| **Bicolored wren** (*Campylorhynchus griseus*( | Females | Only groups with multiple sexually mature females | 100%  (N = 27 offspring, 11 multi-female group-years with an unrelated male) **^1^**  (Haydock, Parker & Rabenold, 1996, pp. 6–7) ^Genetic evidence^ | 100%  (N = 11 multi-female group-years with an unrelated male) **^1^**  (Haydock *et al.*, 1996, pp. 6–7) ^Genetic evidence^ | **Extreme reproductive skew** | **Limited** | **^1^ Population:** Hato Masaguaral research reserve, Venezuela (8°30’N, 67°36’W). **Period:** 1989-1991. | - Raihani & Clutton-Brock (2010) - Cornwallis *et al.* (2010) |
|  | Males | Only groups with multiple sexually mature males | 95.3% ^1^  (N = 43 offspring, 22 multi-male group-years with an unrelated female) ^2^  (Haydock *et al.*, 1996, pp. 6–7) ^Genetic evidence^ | 90.9 – 95.5% ^1, 3^  (N = 22 multi-male group-years with an unrelated female)  (Haydock *et al.*, 1996, pp. 6–7) ^Genetic evidence^ | **Extreme reproductive skew ^4^** | **Limited** | ^1^ In one experimental group (i.e., experimental replacement of the original female breeder) the dominant male sired one offspring and the subordinate male sired 3 offspring (Haydock *et al.*, 1996; Figure 2). This brood was excluded from our analysis. **Population:** Hato Masaguaral research reserve, Venezuela (8°30’N, 67°36’W). **Period:** 1989-1991.  ^2^ Five offspring that were sired by extra-group males were excluded.  ^3^ It is not clear whether the 2 offspring sired by male helpers were from 1 or 2 other groups. Hence, the minimum (95.5%; i.e., 21 groups) and maximum (90.9%; i.e., 20 groups) percentages of groups with a single male breeder are presented.  **^4^** It may be that some of the groups in this sample were experimental groups (Haydock *et al.*, 1996; Figure 2). |  |
| **Stripe-backed wren** (*Campylorhynchus nuchalis*) | Females | Only groups with multiple sexually mature females | 100%  (N = 36 offspring, 19 multi-female group-years) **^1^**  (Rabenold *et al.*, 1990, pp. 538–539) ^Genetic evidence^ | 100%  (N =19 multi-female group-years) **^1^**  (Rabenold *et al.*, 1990, pp. 538–539) ^Genetic evidence^ | **Extreme reproductive skew** | **Limited** | **^1^ Population:** Hato Masaguaral, Venezuela, Estado Guárico (8°30′ N, 67°36′ W).  **Period:** 1988-1989 | - Raihani & Clutton-Brock (2010) - Cornwallis *et al.* (2010) |
|  | Males | Only groups with multiple sexually mature males | 89%  (N = 57 offspring, 28 multi-male group-years) **^1^**  (Rabenold *et al.*, 1990, pp. 538–539) ^Genetic evidence^  85.1%  (N = 47 offspring, 24 group-years, 19 multi-male groups) **^2^**  (Piper & Slater, 1993, p. 237) ^Genetic evidence^ | 69%  (N = 13 multi-male group-years with an unrelated female) **^1^**  (Rabenold *et al.*, 1990, pp. 538–539) ^Genetic evidence^ | **Not extreme reproductive skew ^3^** | **Substantial** | **^1^ Population:** Hato Masaguaral, Venezuela, Estado Guárico (8°30′ N, 67°36′ W).  **Period:** 1988-1989  **^2^** In groups with multiple males of which at least two males were unrelated to the dominant female. Subordinate males sired offspring in 4-5 clutches. See also (Piper, 1994, p. 662) evidence for shared paternity. **Population:** Hato Masaguaral, Venezuela, Estado Guárico (8°30′ N, 67°36′ W). **Period:** 1990-1991.  **^3^** In groups with multiple males, but with only one male unrelated to the dominant female, all offspring were sired by the dominant (unrelated) male (Piper & Slater, 1993, p. 237) ^Genetic evidence^ |  |
| **American crow** (*Corvus brachyrhynchos*) | Females | A mixture of single-female and multi-female groups | 100%  (N = 202 offspring, 60 broods, 21 groups)  (Townsend *et al.*, 2009, p. 507) ^Genetic evidence^ | 100%  (N < 50 groups) **^1^**  (Townsend *et al.*, 2009, p. 507) ^Genetic evidence^ | **Extreme reproductive skew** | **Substantial** | **^1^** The number of groups with multiple adult females is not indicated. Hence, only 10 groups consisting of a single dyad could be excluded from the overall sample of 60 groups. Population: Ithaca, New  York, USA. Period: 2004-2007. | - Raihani & Clutton-Brock (2010) - Cornwallis *et al.* (2010) |
|  | Males | Only groups with multiple sexually mature males | **^1-2^** | 54%  (N = 13 groups) **^2^**  (Townsend *et al.*, 2009, p. 507) ^Genetic evidence^ | **Not extreme reproductive skew** | **Substantial** | **^1^** The mean value for the binomial skew index for males ranged between  *B* = 0.18 - 0.21, which is low, despite being significantly (P < 0.001) different from a random distribution of paternity (*B* ranges from -1 to 2; 0 means that reproductive skew is randomly distributed) (Townsend *et al.*, 2009, p. 507) ^Genetic evidence^.  **^2^** **Population:** Ithaca, New  York, USA. **Period:** 2004-2007.  **^3^** 21 offspring that were sired by extra group males were excluded. We did not use similar data from (Townsend *et al.*, 2009, p. 507) ^Genetic evidence^ since  (Townsend, Clark & McGowan, 2011b, p. 418) ^Genetic evidence^ used the same sample and expended it with further data. **Population**: Ithaca, New  York, USA. **Period:** 2004-2009. |  |
|  |  | A mixture of single-male and multi-male groups | 88%  (N = 269 offspring, 87 broods) **^3^**  (Townsend *et al.*, 2011b, p. 418) ^Genetic evidence^ | Not available |  |  |  |  |
| **Laughing kookaburra**  (*Dacelo novaeguineae*) | Females | Only groups with multiple sexually mature females | 100%  (N = 4 offspring, 2 broods, 2 groups) ^1^  (Legge & Cockburn, 2000, p. 226) ^Genetic evidence^ | 100%  (N = 2 groups) ^1^  (Legge & Cockburn, 2000, p. 226) ^Genetic evidence^ | **Extreme reproductive skew** | **Anecdotal** | ^1^ This sample includes multi-female groups with at least one unrelated male. | - Raihani & Clutton-Brock (2010) - Cornwallis *et al.* (2010) |
|  | Males | Only groups with multiple sexually mature males | 97%  (N = 33 offspring, 12 broods, 10 groups) ^1^  (Legge & Cockburn, 2000, p. 226) ^Genetic evidence^ | 90%  (N = 10 groups) ^1^  (Legge & Cockburn, 2000, p. 226) ^Genetic evidence^ | **Extreme reproductive skew** | **Limited** | ^1^ Multi-male groups with at least one unrelated female |  |
| **Purple-crowned fairy-wren**  (*Malurus coronatus*) | Females | Only groups with multiple sexually mature females |  | 100% **^1^**  (N = 57 broods)  (Kingma, Hall & Peters, 2011; Table 3) ^Genetic evidence^ | **Extreme reproductive skew** | **Substantial** | **^1^** **Population:** Annie Creek  and the Adcock River, Mornington Wildlife Sanctuary, Australia (17°31′S, 126°6′E). **Period:** 2005-2009.  **^2^** One potential scoring error was excluded from the analysis. **Population**: Annie Creek  and the Adcock River, Mornington Wildlife Sanctuary, Australia (17°31′S, 126°6′E). **Period:** 2005-2010. | - Cornwallis *et al.* (2010) |
|  |  | A mixture of single-female and multi-female groups | 100%  (N = 508 offspring, 217 broods) **^2^**  (Kingma, Hall & Peters, 2013, p. 1391) ^Genetic evidence^ |  |  |  |  |  |
|  | Males | Only groups with multiple sexually mature males |  | 98.4% **^1^**  (N = 123 broods)  (Kingma *et al.*, 2011; Table 3) ^Genetic evidence^ | **Extreme reproductive skew ^4^** | **Substantial** | **^1^** **Population:** Annie Creek  and the Adcock River, Mornington Wildlife Sanctuary, Australia (17°31′S, 126°6′E). **Period:** 2005-2009.  **^2^** 34 out of 104 nests did not have helpers (Kingma *et al.*, 2009, p. 5). Assuming that all nests had the same number of offspring, and all helpers were sexually mature males, helpers sired 3% of offspring.  **^3^** 6 offspring from 4 broods were excluded as extra-group paternity. **Population:** Annie Creek  and the Adcock River, Mornington Wildlife Sanctuary, Australia (17°31′S, 126°6′E). **Period:** 2005-2008.  **^4^** See also Kingma et al. (2013, p. 1392). |  |
|  |  | A mixture of single-male and multi-male groups | 98% **^2^**  (N = 221 offspring, 100 broods) **^3^**  (Kingma *et al.*, 2009, p. 3) ^Genetic evidence^ |  |  |  |  |  |
| **Bell miner** (*Manorina melanophrys*) | Females | A mixture of single-female and multi-female groups | Not available | 100%  (N = 8 broods) **^1^**  (Conrad *et al.*, 1998, p. 346) ^Genetic evidence^ | **Not extreme reproductive skew ^2^** | **Limited** | **^1^** This sample includes only broods with helpers, but it is not indicated whether the helpers are male or female. In addition, 59% of (male or female) helpers were first or second-order relatives of the breeding male (Conrad *et al.*, 1998, p. 346). Only broods in which multiple maternity could be detected were considered in our analysis (i.e., clutch size >1). Note, that dominance rank was not assessed in this study and, therefore, “subordinate” females may have produced the entire observed clutch (The reported percentage indicated the percentage of broods with single maternity). Population: Corande Reserve at Healesville, southeastern Victoria, Australia (37°41'S, 145°31'E). Period: 1993-1994  **^2^** The species lives in multi-level societies (Painter *et al.*, 2000; Camerlenghi *et al.*, 2022). The society is built from colonies that are divided into breeding units with contiguous home ranges. Each breeding unit consists of a breeding pair, their offspring, and may also include non-breeding sexually mature members that help. Breeding units that share the same helpers are grouped into “coteries” (Conrad *et al.*, 1998, p. 344). Breeders in one nest may also help in another nest simultaneity (Conrad *et al.*, 1998, p. 346). Painter et al. (2000, p. 1344) suggested that the basic social unit of bell miners is the coterie. Since coteries consist of several breeding pairs, the authors classified the species as a plural breeder. In following this interpretation, we classify the species as not exhibiting extreme reproductive skew. | - Raihani & Clutton-Brock (2010) - Cornwallis *et al.* (2010) |
|  | Males | A mixture of single-male and multi-male groups | Not available | 100%  (N = 8 broods) **^1^**  (Conrad *et al.*, 1998, p. 346) ^Genetic evidence^ | **Not extreme reproductive skew ^2^** | **Limited** | **^1^** This sample includes only broods with helpers, but it is not indicated whether the helpers are male or female. In addition, 31% of (male or female) helpers were first or second-order relatives of the breeding female (Conrad *et al.*, 1998, p. 346). Only broods in which multiple paternity could be detected were considered in our analysis (i.e., clutch size >1). Note, that dominance rank was not assessed in this study and, therefore, “subordinate” males may have sired the entire observed clutch. Namely, we report the percentage of broods with a single breeder.  **^2^** The species lives in multi-level societies (Painter *et al.*, 2000; Camerlenghi *et al.*, 2022). The society is built from colonies that are divided into breeding units with contiguous home ranges. Each breeding unit consists of a breeding pair, their offspring and may also include non-breeding sexually mature members that help. Breeding units that share the same helpers are grouped into “coteries” (Conrad *et al.*, 1998, p. 344). Breeders in one nest may also help in another nest simultaneity (Conrad *et al.*, 1998, p. 346). Painter et al. (2000, p. 1344) suggested that the basic social unit of bell miners is the coterie. Since coteries consist of several breeding pairs, the authors classified the species as a plural breeder. In following this interpretation, we classify the species as not exhibiting extreme reproductive skew. |  |
| **White-fronted bee-eater**  (*Merops bullockoides*) | Females | A mixture of single-female and multi-female groups | ≥94.8% **^1^**  (N = 97 offspring, 65 nest-years)  (Wrege & Emlen, 1987, pp. 156–157) ^Genetic evidence^ | ≥95.4% **^1^**  (N = 65 nest-years)  (Wrege & Emlen, 1987, pp. 156–157) ^Genetic evidence^ | **Not extreme reproductive skew ^2^** | **Substantial** | ^1^ It is not clear whether the offspring that were not produced by the resident female were produced by extra-group maternity or by female helpers. **Population:** Lake Nakuru National Park, Kenya. **Period:** 1982-1984.  **^2^** White-fronted bee-eaters live in multi-levels societies. The basic social unit is a “clan” which is an extended family with up to four overlapping generations. In the breeding season, there are usually multiple breeding pairs within each clan (Emlen, 1990, p. 500; Emlen & Wrege, 1991, p. 310; mean = 1.9 breeding pairs in a season). Helpers are birds that have not been paired yet or pairs that failed to breed earlier in the season (p. 506), but in both cases, helpers are birds from within the clan. Across years, the same birds change from breeders to helpers and vice versa (p. 509). All page numbers are from (Emlen, 1990) ^behavioural and genetic evidence^. Since several pairs usually breed simultaneously within the clan, this species is considered a plural breeder, and we classified it as exhibiting no extreme reproductive skew. | - Cornwallis *et al.* (2010) |
|  | Males | A mixture of single-male and multi-male groups | ≥94.8% **^1^**  (N = 97 offspring, 65 nest-years)  (Wrege & Emlen, 1987, pp. 156–157) ^Genetic evidence^ | ≥93.8% **^1^**  (N = 65 nest-years)  (Wrege & Emlen, 1987, pp. 156–157) ^Genetic evidence^ | **Not extreme reproductive skew ^2^** | **Substantial** | **^1^** It is not clear whether the offsprings which were not of the resident male were the offspring of extra-group paternity or were sired by male helpers. Population: Lake Nakuru National Park, Kenya. Period: 1982-1984.  **^2^** White-fronted bee-eaters live in multi-levels societies. The basic social unit is a “clan” which is an extended family with up to four overlapping generations. In the breeding season, there are usually multiple breeding pairs within each clan (Emlen, 1990, p. 500; Emlen & Wrege, 1991, p. 310; mean = 1.9 breeding pairs in a season). Helpers are birds that have not been paired yet or pairs that failed to breed earlier in the season (p. 506), but in both cases, helpers are birds from within the clan. Across years, the same birds change from breeders to helpers and vice versa (p. 509). All page numbers are from (Emlen, 1990)^behavioural and genetic evidence^. Since several pairs usually breed simultaneously within the clan, this species is considered a plural breeder, and we classified it as exhibiting no extreme reproductive skew. |  |
| **Sociable weaver** (*Philetairus socius*) | Females | Only groups with multiple sexually mature females | 92.3% **^1^**  (N = 13 offspring, 5 nest-years)  (Covas *et al.*, 2006, p. 327) ^Genetic evidence^ | 80% **^1^**  (N = 5 nest-years)  (Covas *et al.*, 2006, p. 327) ^Genetic evidence^ | **Extreme reproductive skew ^2-3^** | **Anecdotical** | **^1^** One offspring was assigned to a female helper based on the highest LOD score. Yet, the other female feeding at the nest, who was also the mother of the female helper, could not be excluded from maternity as well.  **^2^** Sociable weaver females older than one year do not usually help and sexual maturity is not reached until three years old. So sexually mature female helpers in this species are very rare (Doutrelant *et al.*, 2004; Covas *et al.*, 2006).  **^3^** The species live in large colonies with many breeding pairs, in which individuals breed in some seasons and help in other seasons (Covas *et al.*, 2006). Hence, reproduction is not necessarily skewed towards the same individuals in every breeding season. | - Cornwallis *et al.* (2010) |
|  | Males | Only groups with multiple sexually mature males | 96% **^1^**  (N = 50 offspring, 18 nest-years)  (Covas *et al.*, 2006, pp. 326–327) ^Genetic evidence^ | 88.8% **^1^**  (N = 18 nest-years)  (Covas *et al.*, 2006, pp. 326–327) ^Genetic evidence^ | **Extreme reproductive skew ^2-4^** | **Limited** | **^1^** Two offspring were assigned to two male helpers based on the highest LOD score. Yet, the other males feeding at the nests could not be excluded from paternity as well.  **^2^** Although in Covas et al. (2006) sample all groups included multiple sexually mature males, in general, most male helpers are not sexually mature (sexual maturity is at three years old: Doutrelant *et al.*, 2004).  **^3^** Birds may help in one nest while simultaneously breeding at another nest (Covas *et al.*, 2006, p. 326).  **^4^** The species live in large colonies with many breeding pairs, in which individuals breed in some seasons and help in other seasons (Covas *et al.*, 2006). Hence, reproduction is not necessarily skewed towards the same individuals in every breeding season. |  |
| **Grey-crowned babbler** (*Pomatostomus temporalis*) | Females | Only groups with multiple sexually mature females |  | 90.9% **^1^**  (N = 22 brood-years)  (Blackmore & Heinsohn, 2008, pp. 66–67) ^Genetic evidence^ | **Extreme reproductive skew** | **Substantial** | **^1^** Some of the broods may consist of only extra-group maternity. Hence, this percentage may overestimate the reproductive share of dominant females  **^2^** Five offspring were excluded from the analysis as extra-group maternity (assumed to be eggs dumped by females outside the group).  **^3^** Population: Coomalie Farm (13°00′S, 131°08′E), 85 km south of Darwin, Northern Territory, Australia. Period: 2002-2008. The exact number of broods is not indicated but the minimum number of brood-years in which a helper and a nestling were sampled is 26 (page 7). It seems like not all offspring and not all helpers in groups with helpers were sampled. One offspring were excluded as extra-group maternity. | - Raihani & Clutton-Brock (2010) - Cornwallis *et al.* (2010) |
|  |  | A mixture of single-female and multi-female groups | 98%  (N = 112 offspring, 60 brood-years) **^2^**  (Blackmore & Heinsohn, 2008, pp. 66–67) ^Genetic evidence^  91.8%  (N = 86 offspring; ≥26 brood-years) **^3^**  (Mikami *et al.*, 2021; Table 2) ^Genetic evidence^ |  |  |  |  |  |
|  | Males | Only groups with multiple sexually mature males |  | 89.5% **^1^**  (N = 38 brood-years)  (Blackmore & Heinsohn, 2008, p. 66) ^Genetic evidence^  60% **^1-3^**  (N = 5 groups)  (Blackmore & Heinsohn, 2008, p. 67) ^Genetic evidence^ | **Not extreme reproductive skew** | **Substantial** | **^1^** Some of the broods may consist of only extra-group paternity. Hence, this percentage may overestimate the reproductive share of dominant males  **^2^** Only groups with a male helper that was unrelated to the breeding female.  **^3^** 13 of 14 groups (93%) with a male helper that was related to the breeding female had a single male breeder (Blackmore & Heinsohn, 2008, p. 67) ^Genetic evidence^ (comment 1 applies here too).    **^4^** 21 offspring were excluded from the analysis as extra-group paternity.  **^5^** Population: Coomalie Farm (13°00′S, 131°08′E), 85 km south of Darwin, Northern Territory, Australia. Period: 2002-2008. The exact number of broods is not given but the minimum number of brood-years in which a helper and a nestling were sampled is 26 (page 7). It seems like not all offspring and not all helpers in groups with helpers were sampled. Seven offspring were excluded as extra-group paternity. |  |
|  |  | A mixture of single-male and multi-male groups | 95.8%  (N = 96 offspring, 60 brood-years) **^4^**  (Blackmore & Heinsohn, 2008, pp. 66–67) ^Genetic evidence^  96.3%  (N = 80 offspring; ≥26 brood-years) **^5^**  (Mikami *et al.*, 2021; Table 2) ^Genetic evidence^ |  |  |  |  |  |
| **Pied babbler** (*Turdoides bicolor*) | Females | Only groups with multiple sexually mature females | 74.3%  (N = 113 offspring, 50 group-years) **^1^**  (Nelson-Flower, Flower & Ridley, 2018, p. 2440) ^Genetic evidence^ | 66%  (N = 50 group-years) **^1^**  (Nelson-Flower *et al.*, 2018, p. 2440) ^Genetic evidence^ | **Not extreme reproductive skew ^2^** | **Substantial** | **^1^** This sample includes only groups with a subordinate female with an unrelated male in her group. **Population:** Kuruman River Reserve, South Africa (26° 58’ S; 21° 49’ E). **Period:** 2003-2014.  **^2^** The sample included in (Nelson-Flower *et al.*, 2018, p. 2440) includes the samples in (Nelson-Flower *et al.*, 2011, 2013) and the latter studies are, therefore, not presented here. | - Raihani & Clutton-Brock (2010) - Cornwallis *et al.* (2010) |
|  | Males | Only groups with multiple sexually mature males | 85.8%  (N = 134 offspring, 61 group-years) **^1^**  (Nelson-Flower *et al.*, 2018, p. 2440) ^Genetic evidence^ | 78.7%  (N = 61 group-years) **^1^**  (Nelson-Flower *et al.*, 2018, p. 2440) ^Genetic evidence^ | **Not extreme reproductive skew ^2^** | **Substantial** | **^1^** This sample includes only groups with a subordinate male with an unrelated female in his group. **Population:** Kuruman River Reserve, South Africa (26° 58’ S; 21° 49’ E). **Period:** 2003-2014.  **^2^** The sample included in (Nelson-Flower *et al.*, 2018, p. 2440) includes the samples in (Nelson-Flower *et al.*, 2011, 2013) and the latter studies are, therefore, not presented here. |  |
| **Tasmanian native hen**  (*Tribonyx mortierii*) | Females | Only groups with multiple sexually mature females | 100% **^1^**  (N = 12 offspring, 2 groups)  (Gibbs *et al.*, 1994, pp. 367–368) ^Genetic evidence^ | 100% **^1^**  (N = 2 groups)  (Gibbs *et al.*, 1994, pp. 367–368) ^Genetic evidence^  42.9% **^2^**  (N = 9 group-years, 7 groups)  (Goldizen *et al.*, 2000; Table 4) ^Behavioural evidence^ | **Not extreme reproductive skew ^3^** | **Limited** | **^1^** The study used a genetic method that may not be accurate enough to assign maternity. See discussion in Gibbs *et al.* (1994) and (Goldizen *et al.*, 2000)*.* **Population:** Maria Island, Tasmania, Australia (42° 30'S, 148° 00'E). **Period:** 1989.  **^2^** This is the percentage of multi-female groups in which a single female monopolised more than 90% of copulations. The percentage is calculated from the number of groups (group 56 in the year 1995 was considered a new group due to a change in female composition). **Population:** Maria Island, Tasmania, Australia (42° 30'S, 148° 00'E). **Period:** 1993-1996.  **^3^** For evidence of substantial mate sharing in this species see Goldizen, Putland, & Goldizen (1998b), (Goldizen *et al.*, 1998a, p. 530) and Goldizen et al. (2000 Table 4). According to behavioural (copulations) and physiological (clutch size, different egg patterns) evidence, Goldizen et al. (2000; Table 4) concluded that there is significant mate-sharing in most groups with multiple sexually mature females. The authors of this paper were also the authors of the genetic analysis (Gibbs *et al.*, 1994, pp. 367–368) in the same population. They argue that the genetic results may not have been accurate enough because of the close relatedness between breeders (Goldizen et al., 2000; p. 45). In light of the data in both papers, we classified the species as exhibiting no extreme reproductive skew. | - Cornwallis *et al.* (2010) |
|  | Males | Only groups with multiple sexually mature males | 88% **^1^**  (N = 25 offspring, 6 groups) **^2^**  (Gibbs *et al.*, 1994, pp. 367–368) ^Genetic evidence^ | 83% **^1^**  (N = 6 groups)  (Gibbs *et al.*, 1994, pp. 367–368) ^Genetic evidence^  10% **^3^**  (N = 19 group-years, 10 groups)  (Goldizen *et al.*, 2000; Table 3) ^Behavioural evidence^ | **Not extreme reproductive skew ^4^** | **Substantial** | **^1^** The study used a genetic method that may not be accurate enough to assign paternity. See discussion in Gibbs *et al.* (1994) and (Goldizen *et al.*, 2000)*.* Population: Population: Maria Island, Tasmania, Australia (42° 30'S, 148° 00'E). Period: 1989.  **^2^** Two chicks that could not be assigned with certainty to either of the males in the group were excluded from the analysis.  **^3^** This is the percentage of multi-male groups in which a single male monopolised more than 90% of copulations. The percentage is calculated from the number of groups. (group 66 in the year 1996 was considered a new group due to a change in male composition). **Population:** Maria Island, Tasmania, Australia (42° 30'S, 148° 00'E). **Period:** 1993-1996.  **^4^** For evidence of substantial mate sharing in this species see Goldizen, Putland, & Goldizen (1998b), (Goldizen *et al.*, 1998a, p. 530) and Goldizen et al. (2000 Table 3). According to behavioural evidence (copulations), Goldizen et al. (2000 Table 3) concluded that there is significant mate-sharing in most groups with multiple sexually mature males. The authors of this paper were also the authors of the genetic analysis (Gibbs *et al.*, 1994, pp. 367–368) in the same population. They argue that their former genetic results may not have been accurate enough because of the close relatedness between breeders and also because, at the time, the authors considered younger males as sexually mature (Goldizen et al., 2000; p. 45). In light of the data in both papers, we classified the species as exhibiting no extreme reproductive skew. |  |
| **Arabian babbler** (*Turdoides squamiceps*) | Females | Only groups with multiple sexually mature females | 92%  (N = 12 offspring, 4 multi-female group-years with an unrelated male) **^1^**  (Lundy, Parker & Zahavi, 1998, pp. 175–176) ^Genetic evidence^ | 75%  (N = 4 multi-female group-years with an unrelated male) **^1^**  (Lundy *et al.*, 1998, pp. 175–176) ^Genetic evidence^ | **Extreme reproductive skew** | **Anecdotal** | **^1^ Population**: 30 km south of the Dead Sea, Hatzeva, Israel. **Period:**  1993-1995. | - Raihani & Clutton-Brock (2010) - Cornwallis *et al.* (2010) |
|  | Males | Only groups with multiple sexually mature males | 86.5%  (N = 74 offspring, 22 multi-male group-years with an unrelated female) **^1^**  (Lundy *et al.*, 1998, pp. 175–176) ^Genetic evidence^ | 72.7%  (N = 22 multi-male group-years with an unrelated female) **^1^**  (Lundy *et al.*, 1998, pp. 175–176) ^Genetic evidence^ | **Not extreme reproductive skew** | **Limited** | **^1^ Population**: 30 km south of the Dead Sea, Hatzeva, Israel. **Period:**  1993-1995. |  |
| **White-throated magpie-jay** (*Calocitta formosa*) | Females | Only groups with multiple sexually mature females | 84.4%  (N = 51 offspring, 8 groups) **^1^**  (Berg, 2005, pp. 379–380) ^Genetic evidence^  78.9%  (N = 38 successful nests) **^2^**  (Langen, 1996a, p. 511) ^Behavioural evidence^ | 80% **^3^**  (N = 15 clutches, 8 groups)  (Berg, 2005, p. 380) ^Genetic evidence^  47%  (N = 17 group-years) **^4^**  (Langen, 1996a, p. 510) ^Behavioural evidence^ | **Not extreme reproductive skew** | **Substantial** | **^1^** It is not indicated whether helpers were sexually mature and whether all females in the group were sampled for maternity.  Results are from all sampled nests and with 95% confidence of maternity. Populatio**n:** Santa Rosa National Park, Guan-acaste Conservation Area. Guanacaste Province, Costa Rica (85”37’W, 10”50‘N). **Period:** 1999-2002.  **^2^** This is the percentage of successful nests of the primarily female out of the total number of nests of primary and secondary females (namely, the sample size is group-years). **Population:** Santa Rosa National Park, Guan-acaste Conservation Area. Guanacaste Province, Costa Rica (85”37’W, 10”50‘N). **Period:** 1990-1993.  **^3^** This result is from nests maternity could be assigned with 95% confidence and where multiple maternity could be detected (i.e. >1 offspring). Out of the 15 clutches, 2 had multiple maternity and an additional clutch was completely produced by a female helper. The percentage is calculated from the number of clutches.  **Population:** Santa Rosa National Park, Guan-acaste Conservation Area. Guanacaste Province, Costa Rica (85”37’W, 10”50‘N). **Period:** 1999-2002.  **^4^ Population:** Santa Rosa National Park, Guan-acaste Conservation Area. Guanacaste Province, Costa Rica (85”37’W, 10”50‘N). **Period:** 1990-1993. | - Cornwallis *et al.* (2010) |
|  | Males | **Not relevant** **^1^** | | | | | **^1^** Males usually do not provide alloparental care in this species (only 7%-12% of helpers are males)  (Berg, 2005, p. 376). Also, it seems that males disperse before sexual maturity and thus male helpers are not sexually mature (Langen, 1996b, p. 506). Finally, the group’s male usually produces about 65% of offspring while the rest are produced by floaters males (Berg, 2005, pp. 379–380) ^Genetic evidence.^ |  |
| **Brown jay**  (*Cyanocorax morio*) | Females | A mixture of single-female and multi-female groups **^1^** | 91%  (N = 222 nestlings, 87 group-years, 15 groups) **^2^**  (Williams, 2004, p. 375) ^Genetic evidence^ | 76% **^3^**  (N = 87 group-years, 15 groups) **^2^**  (Williams, 2004, p. 375) ^Genetic evidence^  71%  (N = 156 group-years) **^4^**  (Williams, 2004, p. 371) ^behavioural and genetic evidence^ | **Not extreme reproductive skew ^5^** | **Substantial** | **^1^** While it is not clearly indicated that all sampled groups included multiple sexually mature females. This is often the case and reproduction by unassisted pair is unknown in this species (Williams, 2004, p. 371).  **^2^** No evidence for extra group maternity (Williams, 2004, p. 375). Sampling period: 1994-1999. **Population:** Monteverde, Cordillera de Tilarμn (10°12’N, 84°42'W), Costa Rica.  **^3^** The percentage is calculated out of the number of group years.  **^4^** **Sampling period:** 1988-1999. This sampling period partly overlaps with the report on page 375 of the same study. We included this estimation as it represents a longer period, yet not entirely based on genetic evidence. **Population:** Monteverde, Cordillera de Tilarμn (10°12’N, 84°42'W), Costa Rica.  **^5^** The average female binomial skew index across group-years was 0.42 ± 0.09, with a range of 0.25 - 0.6 (n = 29 group years). *B* values can range from -1 to 2. *B =* 0 indicates randomly distributed reproduction, negative values suggest more equally shared reproduction than random and positive values indicate a greater skew than expected by chance (Williams, 2004, p. 376). | - Raihani & Clutton-Brock (2010) - Cornwallis *et al.* (2010) |
|  | Males | A mixture of single-male and multi-male groups **^1^** | 26% **^2-3^**  (N = 38 offspring, 18 broods, 11 groups) **^4^**  (Williams, 2004, p. 373) ^Genetic evidence^ | 57% - 69% **^5^**  (N = 35 broods)  (Williams, 2004, pp. 373–374) ^Genetic evidence^ | **Not extreme reproductive skew ^6^** | **Substantial** | **^1^** While it is not clearly indicated that all sampled groups included multiple sexually mature males. This is often the case and reproduction by unassisted pair is unknown in this species (Williams, 2004, p. 371).  **^2^** This is the percentage of offspring sired by the “consorting” male in the group. All other males in the group were considered helpers (Williams, 2004, p. 372). **Population:** Monteverde, Cordillera de Tilarμn (10°12’N, 84°42'W), Costa Rica.  **^3^** This percentage may underestimate reproduction by the group’s consorting male, as 4 consorting males were not sampled. Yet, a sub-sample that included only broods in which consorting males were sampled resulted in 17.5% of offspring produced by consorting males.  **^4^** 12 offspring were excluded as extra group paternity.  **^5^** This is the range of percentages of broods that were sired by a single father (i.e. the minimum percentage of broods with multiple paternity is 31%). Because ~8 broods had extra group paternity, this sample may be biased in one way or another (Williams, 2004, pp. 373–374). This sample is of broods with more than one nestling. **Population:** Monteverde, Cordillera de Tilarμn (10°12’N, 84°42'W), Costa Rica.  **^6^** The average male binomial skew index across group-years was 0.27 ± 0.2, with a range of 0.17 - 0.6 (n = 28 group years). *B* values can range from -1 to 2. *B =* 0 indicates randomly distributed reproduction, negative values suggest reproduction is more equally shared than random and positive values indicate a greater skew than is expected by random chance (Williams, 2004, p. 376). |  |
| **Apostlebird**  (*Struthidea cinerea*) | Females | Only groups with multiple sexually mature females |  | 75%  (N = 38 offspring, 12 brood-years of 4 multi-female groups) **^1-2^**  (Woxvold & Mulder, 2008, p. 52) ^Genetic evidence^ | **Not extreme reproductive skew** | **Substantial** | **^1^** Only completely typed groups with multiple females were considered (i.e., groups “Gready”, “Eron, “Gallery” [excluding the year 1998b], “G-bud” in Table 2). Percentages are calculated from the number of broods. **Population:** Riverina district of central southern New South Wales, Australia (34° 34’S, 145°45’E).  **^2^** No distinction between dominant and subordinate individuals, so observed offspring could be the offspring of any of these dominance categories.  **^3^** This percentage is an underestimation of “subordinate’s” reproductive share for three reasons: (1) this sample includes all nests in which at least one parent could be identified, so the sample size for nests with identified mothers is smaller than 78 nests; (2) in most nests only one fledgling was sampled (although there were frequently >1 egg), so shared parentage could rarely be detected; (3) in some of these groups there was only a single sexually mature female (Warrington *et al.*, 2013, p. 4675). Hence, if even only 1 out of the 78 nests was in a group with a single female, the percentage of nests with a single breeding female among the groups with multiple sexually mature females would have been 90%. **Population:** Fowlers Gap Arid Zone Research Station (142°E31°S), New South Wales, Australia. | - Raihani & Clutton-Brock (2010) - Cornwallis *et al.* (2010) |
|  |  | A mixture of single-female and multi-female groups |  | 92% **^2-3^**  (N = 78 nest-years)  (Warrington *et al.*, 2013, pp. 4676–4677) ^Genetic evidence^ |  |  |  |  |
|  | Males | Only groups with multiple sexually mature males |  | 60%  (N = 31 offspring, 10 brood-years of 5 multi-male groups) **^1-2^**  (Woxvold & Mulder, 2008, p. 52) ^Genetic evidence^ | **Not extreme reproductive skew ^4^** | **Substantial** | **^1^** Only completely typed groups with multiple males were considered (i.e., groups “Gready 200ab”, “Eron 1998ab”, “Gallery 1998-1999”, “G-bud” and “Sturt 1999” in Table 2).  Percentages are calculated from the number of groups (i.e., 5) because multiple paternity occurred within brood and also within the same group across different roods (i.e. group Gallery). **Population:** Riverina district of central southern New South Wales, Australia (34° 34’S, 145°45’E).  **^2^** No distinction between dominant and subordinate individuals, so observed offspring could be the offspring of any of these dominance categories.  **^3^** This percentage is an underestimation of “subordinate’s” reproductive share for four reasons: (1) this sample includes all nests in which at least one parent could be identified, so the sample size of nests with identified fathers is smaller than 78 nests; (2) in most nests only one fledgling was sampled (although there were frequently >1 eggs), so shared parentage could rarely be detected; (3) in some of these groups there was only a single sexually mature male (Warrington *et al.*, 2013, p. 4675). Hence, if even only 1 out of the 78 nests was in a group with a single male, the percentage of nests with a single breeding male among the groups with multiple sexually mature males would be 90%; (4) in this sample sequential polyandry was prevalent within and between years (Warrington et al., 2015, p. 1315). **Population:** Fowlers Gap Arid Zone Research Station (142°E31°S), New South Wales, Australia.  **^4^** See also (Warrington *et al.*, 2015; Table 4) for evidence of a significant prevalence of sequential polyandry also in the population in Fowlers Gap Arid Zone Research Station (142°E31°S), New South Wales, Australia. |  |
|  |  | A mixture of single-male and multi-male groups |  | 92%  (N = 78 nest-years) **^2-3^**  (Warrington *et al.*, 2013, pp. 4676–4677) ^Genetic evidence^ |  |  |  |  |
| **Long-tailed tits**  )*Aegithalos caudatus*) | Females | A mixture of single-female and multi-female groups **^1^** | 100% **^2^**  (N = 365 offspring, 51 brood-years)  (Hatchwell *et al.*, 2002, pp. 57–61) ^Genetic evidence^ | 100% **^2^**  (N = 51 brood-years)  (Hatchwell *et al.*, 2002, pp. 57–61) ^Genetic evidence^ | **Not relevant ^3^** | **Substantial** | **^1^** Only 12.8% of the helpers in this study were female. But it is not clear how many groups had female helpers. In addition, in about 50% of the nests, there was at least one helper (Hatchwell *et al.*, 2002, pp. 61–62).  **^2^** Two cases of “joint nesting” were excluded as the authors describe these cases as non-representative or since the eggs of one of the females did not hatch (Hatchwell *et al.*, 2002, pp. 57–61). **Population:** Rivelin Valley, Sheffield, U.K. **Period:** 1994-2000.  **^3^** Long-tailed tits live in relatively stable flocks during the non-breeding season. Flocks consist of one or more nuclear families and unrelated individuals. At the beginning of the breeding season, flocks dissolve into monogamous pairs (Glen & Perrins, 1988; Hatchwell *et al.*, 2001). All individuals attempt to breed. Failed breeders may try to re-nest during the season with the same or another partner (Hatchwell et al., 2002, p. 58). Alternatively, failed breeders may turn to help other breeding pairs (Hatchwell *et al.*, 2002, p. 56). Most helpers join successfully breeding pairs after the clutch has been completed and thus have no opportunities to share reproduction. At the end of the breeding season, the breeding pair and their offspring join other families to establish new winter flocks (Hatchwell *et al.*, 2000; Napper & Hatchwell, 2016, p. 27).  We do not consider this species to be a clear case of a species with extreme reproductive skew for the following reasons: (i) all individuals in a flock attempt to breed; (ii) at the time in which parentage is determined (i.e., fertilisation of eggs) there are no social groups present (i.e., only pairs exist); (iii) breeding pairings are not necessarily stable even within a breeding season (i.e., failed breeders may divorce); (iv) after the breeding season, birds do not re-establish the same winter flocks they had before the breeding season (i.e., winter flock have no stable composition). | - Raihani & Clutton-Brock (2010) - Cornwallis *et al.* (2010) |
|  | Males | Only groups with multiple sexually mature males | 94%  (N = 284 offspring, 38 brood-years, ≥14 groups) **^1^**  (Hatchwell *et al.*, 2002, p. 58) ^Genetic evidence^ | 76.3%  (N = 38 brood-years, ≥14 groups)  (Hatchwell *et al.*, 2002, p. 58) ^Genetic evidence^ | **Not relevant ^2^** | **Substantial** | **^1^** Four sampled offspring could not be assigned to any sampled male and were thus excluded from our analysis. **Population:** Rivelin Valley, Sheffield, U.K. **Period:** 1994-2000.  **^2^** Long-tailed tits live in relatively stable flocks during the non-breeding season. Flocks consist of one or more nuclear families and unrelated individuals. At the beginning of the breeding season, flocks dissolve into monogamous pairs (Glen & Perrins, 1988; Hatchwell et al., 201). All individuals attempt to breed. Failed breeders may try to re-nest during the season with the same partner or another partner (Hatchwell et al., 2002, p. 58). Alternatively, failed breeders may turn to help other breeding pairs (Hatchwell *et al.*, 2002, p. 56). Most helpers join successfully breeding pairs after the clutch has been completed and thus have no opportunities to share reproduction. At the end of the breeding season, the breeding pairs and their offspring join other families to establish new winter flocks.  We do not consider this species to be a clear case of a species with extreme reproductive skew for the following reasons: (i) all individuals in a flock attempt to breed; (ii) at the time in which parentage is determined (i.e., fertilisation of eggs) there are no social groups present (i.e., only pairs exist); (iii) breeding pairings are not necessarily stable even within a breeding season (i.e., failed breeders may divorce); (iv) after the breeding season, birds do not re-establish the same winter flocks they had before the breeding season (i.e., winter flock have no stable composition). |  |

**References**

Van Bael, S. & Pruett-Jones, S. (2000) Breeding biology and social behaviour of the eastern race of the Splendid Fairy-wren Malurus splendens melanotus. *Emu* **100**, 95–108.

Baglione, V., Marcos, J.M., Canestrari, D. & Ekman, J. (2002) Direct fitness benefits of group living in a complex cooperative society of carrion crows, *Corvus corone corone*. *Animal Behaviour* **64**, 887–893.

Barati, A., Andrew, R.L., Gorrell, J.C., Etezadifar, F. & McDonald, P.G. (2018a) Genetic relatedness and sex predict helper provisioning effort in the cooperatively breeding noisy miner. *Behavioral Ecology* **29**, 1380–1389.

Barati, A., Andrew, R.L., Gorrell, J.C. & McDonald, P.G. (2018b) Extra-pair paternity is not driven by inbreeding avoidance and does not affect provisioning rates in a cooperatively breeding bird, the noisy miner (*Manorina melanocephala*). *Behavioral Ecology* **29**, 244–252.

Beck, N.R. (2006) Causes and consequences of dispersal in an obligate cooperative breeder, the white-winged chough (*Corcorax melanorhamphos*). Ph.D, The Australian National University.

Berg, E.C. (2005) Parentage and reproductive success in the white-throated magpie-jay, *Calocitta formosa*, a cooperative breeder with female helpers. *Animal Behaviour* **70**, 375–385.

Blackmore, C.J. & Heinsohn, R. (2008) Variable mating strategies and incest avoidance in cooperatively breeding grey-crowned babblers. *Animal Behaviour* **75**, 63–70.

Bowen, B.S., Koford, R.R. & Brown, J.L. (1995) Genetic Evidence for Undetected Alleles and Unexpected Parentage in the Gray-Breasted Jay. *The Condor* **97**, 503–511.

Brooked, M.G., Rowley, I., Adams, M. & Baverstock, P.R. (1990) Promiscuity: an inbreeding avoidance mechanism in a socially monogamous species? *Behavioral Ecology and Sociobiology* **26**, 191–199.

Bruce, J.P., Quinn, J.S., Sloane, S.A. & White, B.N. (1996) DNA fingerprinting reveals monogamy in the bushtit, a cooperatively breeding species. *The Auk* **113**, 511–516.

Camerlenghi, E., McQueen, A., Delhey, K., Cook, C.N., Kingma, S.A., Farine, D.R. & Peters, A. (2022) Cooperative breeding and the emergence of multilevel societies in birds. *Ecology Letters* **25**, 766–777.

Canestrari, D., Marcos, J.M. & Baglione, V. (2005) Effect of parentage and relatedness on the individual contribution to cooperative chick care in carrion crows *Corvus corone corone*. *Behavioral Ecology and Sociobiology* **57**, 422–428.

Carrick, R. (1972) Population ecology of the black-backed magpie, royal penguin and silver gulls. *Population ecology of migratory birds* **2**, 41. Bureau of Sport Fisheries and Wildlife.

Charmantier, A., Keyser, A.J. & Promislow, D.E.L. (2007) First evidence for heritable variation in cooperative breeding behaviour. *Proceedings of the Royal Society B: Biological Sciences* **274**, 1757–1761.

Colombelli-Négrel, D., Schlotfeldt, B.E. & Kleindorfer, S. (2009) High levels of extra-pair paternity in Superb Fairy-wrens in South Australia despite low frequency of auxiliary males. *Emu* **109**, 300–304.

Conrad, K.F., Clarke, M.F., Robertson, R.J. & Boag, P.T.. (1998) Paternity and the relatedness of helpers in the cooperatively breeding bell miner. *The Condor* **100**, 343–349.

Cornwallis, C.K., West, S.A., Davis, K.E. & Griffin, A.S. (2010) Promiscuity and the evolutionary transition to complex societies. *Nature* **466**, 969–972.

Covas, R., Dalecky, A., Caizergues, A. & Doutrelant, C. (2006) Kin associations and direct vs indirect fitness benefits in colonial cooperatively breeding sociable weavers *Philetairus socius*. *Behavioral Ecology and Sociobiology* **60**, 323–331.

Dickinson, J.L. & Akre, J.J. (1998) Extrapair paternity, inclusive fitness, and within-group benefits of helping in western bluebirds. *Molecular Ecology* **7**, 95–105.

Dickinson, J.L., Koenig, W.D. & Pitelka, F.A. (1996) Fitness consequences of helping behavior in the western bluebird. *Behavioral Ecology* **7**, 168–177.

Doutrelant, C., Covas, R., Caizergues, A. & Du Plessis, M. (2004) Unexpected sex ratio adjustment in a colonial cooperative bird: pairs with helpers produce more of the helping sex whereas pairs without helpers do not. *Behavioral Ecology and Sociobiology* **56**, 149–154.

Dow, D.D. (1979) The influence of nests on the social behaviour of males in manorina melanocephala, a communally breeding honeyeater. *Emu* **79**, 71–83.

Dunn, P.O. & Cockburn, A. (1999) Extrapair mate choice and honest signaling in cooperatively breeding superb fairy-wrens. *Evolution* **53**, 938–946.

Dunn, P.O., Cockburn, A. & Mulder, R.A. (1995) Fairy-wren helpers often care for young to which they are unrelated. *Proceedings of the Royal Society B: Biological Sciences* **259**, 339–343.

Durrant, K.L. & Hughes, J.M. (2005) Differing rates of extra-group paternity between two populations of the Australian magpie (*Gymnorhina tibicen*). *Behavioral Ecology and Sociobiology* **57**, 536–545.

Eimes, J.A. (2004) Extra-pair fertilization, mate choice and genetic similarity in the Mexican jay (*Aphelocoma ultramarina*). M.Sc, University of Missouri-St. Louis.

Eimes, J.A., Parker, P.G., Brown, J.L. & Brown, E.R. (2005) Extrapair fertilization and genetic similarity of social mates in the Mexican jay. *Behavioral Ecology* **16**, 456–460.

Emlen, S.T. (1990) White-fronted Bee-eaters: helping in a colonially nesting species. In *Cooperative breeding in birds: Long-term studies of ecology and behavior* (eds P. Stacey & W. Koenig), pp. 487–526. Cambridge University Press, Cambridge.

Emlen, S.T. & Wrege, P.H. (1991) Breeding Biology of White-Fronted Bee-Eaters at Nakuru: The Influence of Helpers on Breeder Fitness. *Journal of Animal Ecology* **60**, 309–326.

Fulton, G.R. (2006) Plural-breeding Australian Magpies *Gymnorhina tibicen dorsalis* Nesting annually in the same tree. *Australian Field Ornitholog* **23**, 198–201.

Gibbs, L.H., Goldizen, A.W., Bullough, C. & Goldizen, A.R. (1994) Parentage analysis of multi-male social groups of tasmanian native hens (*Tribonyx mortierii*): genetic evidence for monogamy and polyandry. *Behavioral Ecology and Sociobiology* **35**, 363–371.

Glen, N.W. & Perrins, C.M. (1988) Co-operative breeding by Long-tailed Tits. *British Birds* **81**, 630–641.

Goldizen, A.W., Buchan, J.C., Putland, D.A., Goldizen, A.R. & Krebs, E.A. (2000) Patterns of mate-sharing in a population of Tasmanian Native Hens *Gallinula mortierii*. *Ibis* **142**, 40–47.

Goldizen, A.W., Goldizen, A.R., Putland, D.A., Lambert, D.M., Millar, C.D. & Buchan, J.C. (1998a) ‘Wife-Sharing’ in the Tasmanian Native Hen (*Gallinula mortierii*): Is It Caused by a Male-Biased Sex Ratio? *The Auk* **115**, 528–532.

Goldizen, A.W., Putland, D.A. & Goldizen, A.R. (1998b) Variable mating patterns in Tasmanian native hens (*Gallinula mortierii*): Correlates of reproductive success. *Journal of Animal Ecology1* **67**, 307–317.

Groenewoud, F., Kingma, S.A., Hammers, M., Dugdale, H.L., Burke, T., Richardson, D.S. & Komdeur, J. (2018) Subordinate females in the cooperatively breeding Seychelles warbler obtain direct benefits by joining unrelated groups. *Journal of Animal Ecology* **87**, 1251–1263. Blackwell Publishing Ltd.

Haig, S.M., Belthoff, J.R. & Allen, D.H. (1993) Examination of population structure in red-cockaded woodpeckers using DNA profiles. *Evolution* **47**, 185–194.

Haig, S.M., Walters, J.R. & Plissner, J.H. (1994) Genetic evidence for monogamy in the cooperatively breeding red-cockaded woodpecker. *Behavioral Ecology and Sociobiology* **34**, 295–303.

Hajduk, G.K., Cockburn, A., Osmond, H.L. & Kruuk, L.E.B. (2021) Complex effects of helper relatedness on female extrapair reproduction in a cooperative breeder. *Behavioral Ecology* **32**, 386–394.

Hatchwell, B.J., Anderson, C., Ross, D.J., Fowlie, M.K. & Blackwell, P.G. (2001) Social organization of cooperatively breeding long-tailed tits: Kinship and spatial dynamics. *Journal of Animal Ecology* **70**, 820–830.

Hatchwell, B.J., Ross, D.J., Chaline, N., Fowlie, M.K. & Burke, T. (2002) Parentage in the cooperative breeding system of long-tailed tits, *Aegithalos caudatus*. *Animal Behaviour* **64**, 55–63.

Hatchwell, B.J., Russell, A.F., Ross, D.J. & Fowlie, M.K. (2000) Divorce in cooperatively breeding long-tailed tits: a consequence of inbreeding avoidance? *Proceedings of the Royal Society of London. Series B: Biological Sciences* **267.1445**, 813–819.

Haydock, J. & Koenig, W.D. (2002) Reproductive skew in the polygynandrous acorn woodpecker. *PNAS* **99**, 7178–7183.

Haydock, J. & Koenig, W.D. (2003) Patterns of reproductive skew in the polygynandrous acorn woodpecker. *The American Naturalist* **162**, 277–289.

Haydock, J., Koenig, W.D. & Stanback, M.T. (2001) Shared parentage and incest avoidance in the cooperatively breeding acorn woodpecker. *Molecular Ecology* **10**, 1515–1525.

Haydock, J., Parker, P.G. & Rabenold, K.N. (1996) Extra-pair paternity uncommon in the cooperatively breeding bicolored wren. *Behavioral Ecology and Sociobiology* **38**, 1–16.

Heinsohn, R., Dunn, P., Legge, S. & Double, M. (2000) Coalitions of relatives and reproductive skew in cooperatively breeding white-winged choughs. *Proceedings of the Royal Society B: Biological Sciences* **267**, 243–249.

Hughes, J.M., Hesp, J.D.E., Kallioinen, R., Kempster, M., Lange, C.L., Hedstrom, K.E., Mather, P.B., Robinson, A. & Wellbourn, M.J. (1996) Differences in social behaviour between populations of the Australian magpie *Gymnorhina tibicen*. *Emu* **96**, 65–70. CSIRO.

Hughes, J.M., Mather, P.B., Toon, A., Ma, J., Rowley, I. & Russell, E. (2003) High levels of extra-group paternity in a population of Australian magpies *Gymnorhina tibicen*: evidence from microsatellite analysis. *Molecular Ecology* **12**, 3441–3450.

James, P.C. & Oliphant, L.W. (1986) Extra birds and helpers at the nests of Richardson’s merlin. *The Condor* **88**, 533–534.

Joste, N., Ligon, D. & Stacey, P.B. (1985) Shared paternity in the acorn woodpecker (*Melanerpes formicivorus*). *Behavioral Ecology and Sociobiology* **17**, 39–41.

Kingma, S.A., Hall, M.L. & Peters, A. (2011) Multiple Benefits Drive Helping Behavior in a Cooperatively Breeding Bird: An Integrated Analysis. *The American Naturalists* **177**, 486–495.

Kingma, S.A., Hall, M.L. & Peters, A. (2013) Breeding synchronization facilitates extrapair mating for inbreeding avoidance. *Behavioral Ecology* **24**, 1390–1397.

Kingma, S.A., Hall, M.L., Segelbacher, G. & Peters, A. (2009) Radical loss of an extreme extra-pair mating system. *BMC Ecology* **9**, 1–11.

Komdeur, J. (2005) No evidence for adaptive suppression of joint laying by dominant female Seychelles warblers: an experimental study. *Behaviour* **142**, 1669–1684.

Langen, T.A. (1996a) The mating system of the White-throated Magpie-jay *Calocitta formosa* and Greenwood’s hypothesis for sex-biased dispersal. *Ibis* **138**, 506–513. Blackwell Publishing Ltd.

Langen, T.A. (1996b) The mating system of the White-throated Magpie-jay *Calocitta formosa* and Greenwood’s hypothesis for sex-biased dispersal. *Ibis* **138**, 506–513.

Legge, S. & Cockburn, A. (2000) Social and mating system of cooperatively breeding laughing kookaburras (*Dacelo novaeguineae*). *Behavioral Ecology and Sociobiology* **47**, 220–229.

Lennartz, M.R., Hooper, R.G. & Harlow, R.F. (1987) Sociality and cooperative breeding of red-cockaded woodpeckers, *Picoides borealis*. *Behavioral Ecology and Sociobiology* **20**, 77–88.

Ligon, David.J. & Ligon, S.H. (1990) Green Woodhoopoes: life history traits and sociality. In *Cooperative breeding in birds: Long-term studies of ecology and behavior* (eds P.B. Stacey & W.D. Koenig), pp. 33–65. Cambridge University Press, Cambridge.

Li, S.-H. (1997) The genetic analysis of paternity pattern in a natural population of Mexican jays (Aphelocoma ultramarina). Ph.D. dissertation, Albany: State University of New York.

Li, S.H. & Brown, J.L. (2000) High frequency of extrapair fertilization in a plural breeding bird, the Mexican jay, revealed by DNA microsatellites. *Animal Behaviour* **60**, 867–877.

Loyau, A. & Schmeller, D.S. (2012) Mixed reproductive strategies of the Common moorhen on a microscale as revealed by genetic data. *Comptes Rendus - Biologies* **335**, 673–679.

Lundy, K.J., Parker, P.G. & Zahavi, A. (1998) Reproduction by subordinates in cooperatively breeding Arabian babblers is uncommon but predictable. *Behavioral Ecology and Sociobiology* **43**, 173–180.

Magrath, R.D. & Whittingham, L.A. (1997) Subordinate males are more likely to help if unrelated to the breeding female in cooperatively breeding white-browed scrubwrens. *Behavioral Ecology and Sociobiology* **41**, 185–192.

McRae, S.B. (1996) Family values: costs and benefits of communal nesting in the moorhen. *Animal Behaviour* **52**, 225–245.

Mikami, K., Yamaguchi, N., Noske, R.A. & Eguchi, K. (2021) Male and female helpers of Grey-crowned Babblers *Pomatostomus temporalis rubecula* acquire breeding positions in different ways, and don’t avoid incest. *Ornithol Sci* **20**, 3–13.

Ben Mocha, Y., Scemama de Gialluly, S. & Markman, S. (undated) What is cooperative breeding in mammals and birds? Removing definitional barriers for comparative research. *Major revisions in Biological Reviews*.

Napper, C.J. & Hatchwell, B.J. (2016) Social dynamics in nonbreeding flocks of a cooperatively breeding bird: causes and consequences of kin associations. *Animal Behaviour* **122**, 23–35. Academic Press.

Nelson-Flower, M.J., Flower, T.P. & Ridley, A.R. (2018) Sex differences in the drivers of reproductive skew in a cooperative breeder. *Molecular Ecology* **27**, 2435–2446. Blackwell Publishing Ltd.

Nelson-Flower, M.J., Hockey, P.A.R., O’Ryan, C., English, S., Thompson, A.M., Bradley, K., Rose, R. & Ridley, A.R. (2013) Costly reproductive competition between females in a monogamous cooperatively breeding bird. *Proceedings of the Royal Society B: Biological Sciences* **280**. Royal Society.

Nelson-Flower, M.J., Hockey, P.A.R., O’Ryan, C., Raihani, N.J., Du Plessis, M.A. & Ridley, A.R. (2011) Monogamous dominant pairs monopolize reproduction in the cooperatively breeding pied babbler. *Behavioral Ecology* **22**, 559–565.

North, A.J. (1901) *Nest and eggs of birds found breeding in Australia and Tasmania. Volume I*. Australian Museum, Sydney.

Öst, M. (1999) Within-season and between-year variation in the structure of Common Eider broods. *Condor* **101**, 598–606.

Öst, M., Clark, C.W., Kilpi, M. & Ydenberg, R. (2007) Parental effort and reproductive skew in coalitions of brood rearing female common eiders. *American Naturalist* **169**, 73–86.

Öst, M., Ydenberg, R., Kilpi, M. & Lindström, K. (2003) Condition and coalition formation by brood-rearing common eider females. *Behavioral Ecology* **14**, 311–317.

Painter, J.N., Crozier, R.H., Poiani, A., Robertson, R.J. & Clarke, M.F. (2000) Complex social organization reflects genetic structure and relatedness in the cooperatively breeding bell miner, *Manorina melanophrys*. *Molecular Ecology* **9**, 1339–1347.

Piper, W.H. (1994) Courtship, Copulation, Nesting Behavior and Brood Parasitism in the Venezuelan Stripe-Backed Wren. *The Condor* **96**, 654–671.

Piper, W.H. & Slater, G. (1993) Polyandry and Incest Avoidance in the Cooperative Stripe-Backed Wren of Venezuela. *Behaviour* **124**, 227–247.

Põldmaa, T., Montgomerie, R. & Boag, P. (1995) Mating system of the cooperatively breeding noisy miner *Manorina melanocephala*, as revealed by DNA profiling. *Behavioral Ecology and Sociobiology* **37**, 137–143.

Rabenold, P.P., Rabenold, K.N., Piper, W.H., Haydock, J. & Zack, S.W. (1990) Shared paternity revealed by genetic analysis in cooperatively breeding tropical wrens. *Nature* **348**, 538‐540.

Raihani, N.J. & Clutton-Brock, T.H. (2010) Higher reproductive skew among birds than mammals in cooperatively breeding species. *Biology letters* **6**, 630–632.

Raj Pant, S., Komdeur, J., Burke, T.A., Dugdale, H.L. & Richardson, D.S. (2019) Socio-ecological conditions and female infidelity in the Seychelles warbler. *Behavioral Ecology* **30**, 1254–1264. Oxford University Press.

Richardson, D., Komdeur, J. & Burke, T. (2003) Altruism and infidelity among warblers. *Nature* **422**, 580.

Richardson, D.S., Burke, T. & Komdeur, J. (2002) Direct Benefits and the Evolution of Female-Biased Cooperative Breeding in Seychelles Warblers. *Evolution* **56**, 2313–2321.

Richardson, D.S., Jury, F.L., Blaakmeer, K., Komdeur, J. & Burke, T. (2001) Parentage assignment and extra-group paternity in a cooperative breeder: The Seychelles warbler (*Acrocephalus sechellensis*). *Molecular Ecology* **10**, 2263–2273.

Rowley, I. (1965) The life history of the Superb Blue Wren, Malurus cyaneus. *Emu* **64**, 251–297.

Rowley, I. (1978) Communal Activities Among White‐Winged Choughs Corcorax Melanorhamphus. *Ibis* **120**, 178–197.

Rowley, I., Russell, E., Payne, R.B. & Payne, L.L. (1989) Plural Breeding in the Splendid Fairy‐wren, *Malurus splendens* (Aves: Maluridae), a Cooperative Breeder. *Ethology* **83**, 229–247.

Rubenstein, D.R. (2007) Female extrapair mate choice in a cooperative breeder: trading sex for help and increasing offspring heterozygosity. *Proceedings of the Royal Society B: Biological Sciences* **274**, 1895–1903.

Rubenstein, D.R. (2016) Superb starlings: cooperation and conflict in an unpredictable environment. In *Cooperative Breeding in Vertebrates: Studies of Ecology, Evolution, and Behavior* (eds W.D. Koenig & J.L. Dickinson), pp. 181–196. Cambridge University Press.

Seddon, N., Amos, W., Adcock, G., Johnson, P., Kraaijeveld, K., Kraaijeveld-Smit, F.J.L., Lee, W., Senapathi, G.D., Mulder, R.A. & Tobias, J.A. (2005) Mating system, philopatry and patterns of kinship in the cooperatively breeding subdesert mesite *Monias benschi*. *Molecular Ecology* **14**, 3573–3583.

Shen, S.F., Emlen, S.T., Koenig, W.D. & Rubenstein, D.R. (2017) The ecology of cooperative breeding behaviour. *Ecology Letters* **20**, 708–720.

Sloane, S.A. (1996) Incidence and origins of supernumeraries at bushtit (*Psaltriparus minimus*) nests. *The Auk* **113**, 757–770.

Sodhi, N.S. (1989) Attempted Polygyny by a Merlin. *The Wilson Bulletin* **101**, 506–507.

Sodhi, N.S. (1991) Pair copulations, extra-pair copulations, and intraspecific nest intrusions in Merlin. *The Condor* **93**, 433–437.

Temple, H.J., Hoffman, J.I. & Amos, W. (2009) Group structure, mating system and extra-group paternity in the co-operatively breeding White-breasted Thrasher *Ramphocinclus brachyurus*. *Ibis* **151**, 99–112.

Townsend, A.K., Bowman, R., Fitzpatrick, J.W., Dent, M. & Lovette, I.J. (2011a) Genetic monogamy across variable demographic landscapes in cooperatively breeding Florida scrub-jays. *Behavioral Ecology* **22**, 464–470.

Townsend, A.K., Clark, A.B. & McGowan, K.J. (2011b) Injury and paternity loss in cooperatively breeding American Crows. *Journal of Field Ornithology* **82**, 415–421.

Townsend, A.K., Clark, A.B., McGowan, K.J. & Lovette, I.J. (2009) Reproductive partitioning and the assumptions of reproductive skew models in the cooperatively breeding American crow. *Animal Behaviour* **77**, 503–512.

Waldeck, P., Kilpi, M., Öst, M. & Andersson, M. (2004) Brood parasitism in a population of common eider (*Somateria mollissima*). *Behaviour* **141**, 725–739.

Walters, J., Doerr, P. & Carter, J.I. (1988) The cooperative breeding system of the red-cockaded woodpecker. *Ethology* **78**, 275–305.

Warkentin, I.G., Curzon, A.D., Carter, R.E., Wetton, J.H., James, C., P., Oliphant, L.W. & Parkin, D.T. (1994) No evidence for extrapair fertilizations in the merlin revealed by DNA fingerprinting. *Molecular Ecology* **3**, 229–234.

Warkentin, I.G., Lieske, D.J., Espie, R.H.M. & James, P. (2013) Close Inbreeding and Dispersal in Merlins: Further Examination. *Journal of Raptor Research* **47**, 69–74.

Warrington, M.H., Rollins, L.A., Raihani, N.J., Russell, A.F. & Griffith, S.C. (2013) Genetic monogamy despite variable ecological conditions and social environment in the cooperatively breeding apostlebird. *Ecology and Evolution* **3**, 4669–4682.

Warrington, M.H., Rollins, L.A., Russell, A.F. & Griffith, S.C. (2015) Sequential polyandry through divorce and re-pairing in a cooperatively breeding bird reduces helper-offspring relatedness. *Behavioral Ecology and Sociobiology* **69**, 1311–1321. Springer Verlag.

Webster, M.S., Tarvin, K.A., Tuttle, E.M. & Pruett-Jones, S. (2004) Reproductive promiscuity in the splendid fairy-wren: effects of group size and auxiliary reproduction. *Behavioral Ecology* **15**, 907–915.

Webster, M.S., Tarvin, K.A., Tuttle, E.M. & Pruett-Jones, S. (2007) Promiscuity drives sexual selection in a socially monogamous bird. *Evolution* **61**, 2205–2211.

Weinman, L.R., Solomon, J.W. & Rubenstein, D.R. (2015) A comparison of single nucleotide polymorphism and microsatellite markers for analysis of parentage and kinship in a cooperatively breeding bird. *Molecular Ecology Resources* **15**, 502–511.

Whittingham, L.A., Dunn, P.O. & Magrath, R.D. (1997) Relatedness, polyandry and extra-group paternity in the cooperatively-breeding white-browed scrubwren (*Sericornis frontalis*). *Behavioral Ecology and Sociobiology* **40**, 261–270.

Williams, D.A. (2004) Female control of reproductive skew in cooperatively breeding brown jays (*Cyanocorax morio*). *Behavioral Ecology and Sociobiology* **55**, 370–380.

Windsor, R.L., Tringali, A., Dent, M. & Bowman, R. (2021) Rare occurrences of polygyny in the monogamous Florida Scrub-Jay (*Aphelocoma coerulescens*): Synthesis of 97 combined years of population monitoring across 3 populations. *Wilson Journal of Ornithology* **133**, 484–490.

Woxvold, I.A. & Mulder, R.A. (2008) Mixed mating strategies in cooperatively breeding apostlebirds *Struthidea cinerea*. *Journal of Avian Biology* **39**, 50–56.

Wrege, P.H. & Emlen, S.T. (1987) Biochemical determination of parental uncertainty in white-fronted bee-eaters. *Behavioral Ecology and Sociobiology* **20**, 153–160.
